# Role of copper during microglial inflammation

**DOI:** 10.1101/2024.09.18.613750

**Authors:** Laura Craciun, Sandra E. Muroy, Kaoru Saijo

## Abstract

Copper plays crucial roles in various physiological functions of the nervous and immune systems. Dysregulation of copper homeostasis is linked to several diseases, including neurodegenerative diseases. Since dysfunctional microglial immunity can contribute to such diseases, we investigated the role of copper in microglial immunity. We found that both increased and decreased copper levels induced by chemical treatments suppresses lipopolysaccharide (LPS)-mediated inflammation in microglial cells, as determined by RT-qPCR analysis. RNA sequencing (RNA-seq) analysis confirmed that increased copper level reduces the inflammatory response to LPS; however, it also showed that decreased copper level affects genes involved in cell proliferation, transcription, and autophagosome regulation. These findings suggest that copper is vital for maintaining normal immune function in microglia, and both copper excess and deficiency can disrupt microglial immunity.

## INTRODUCTION

Copper is the third most abundant metal in the human body^1–4^. It regulates critical physiological functions in many systems, including the nervous and immune systems^5–11^. Cellular levels of copper are precisely maintained by copper transporters. Copper transporter 1 (CTR1), encoded by the solute carrier family 31 member 1 (*SLC31A1*) gene, is located on the plasma membrane and predominantly mediates copper intake^12,13^. In contrast, the ATPases, ATPase copper transporting alpha and beta (ATP7A and ATP7B, respectively), are the main copper exporters. They are located on the Golgi apparatus and sort copper into the lumen of the secretory compartment to be loaded onto cuproenzymes, such as ceruloplasmin^14–16^. ATP7A can also move to the plasma membrane for copper export when intracellular levels of copper are high^17–19^. Although ATP7A and ATP7B perform similar functions, they are not biologically redundant^20,21^. Mutations in these copper transporters result in rare genetic disorders, such as Wilson’s disease (caused by alterations in *ATP7B*) and Menkes disease (caused by defective *ATP7A*), which are associated with various neurological and other symptoms^6,17,22–24^. Neurological symptoms of Menkes disease are characterized by low muscle tone (floppy child) and seizures^24,25^. Individuals who are diagnosed with Wilson’s disease develop symptoms similar to Parkinson’s disease, such as tremors and rigidity. They also showed some neuropsychiatric symptoms, including anxiety and hallucinations^26,27^. Copper dysregulation is also linked to many neurodegenerative diseases, such as Alzheimer’s disease, Parkinson’s disease, and amyotrophic lateral sclerosis (ALS) ^6,22,28–30^. Overall, this evidence suggests that copper is critical for maintaining normal physiologic functions in the human body.

Microglial cells are resident innate immune cells in the central nervous system (CNS) whose immune functions protect the CNS from infection and injury. In addition, microglia play critical roles in brain development and maintaining homeostasis in the CNS^31–33^. Dysfunctional microglia-mediated immune responses are observed in many neurological diseases, including neurodegenerative diseases^34–36^. Although the impact of copper dysregulation in microglia is not well understood, many reports suggest that copper plays critical roles in other innate immune cells, such as macrophages. Some reports indicate that copper acts in a proinflammatory manner by increasing the production of cytokines, reactive oxygen species (ROS), and reactive nitrogen species (RNS), as well as recruiting phagocytic cells^37–40^. In addition, it has been shown that copper changes the epigenetic status of immune cells and promotes inflammation^37,41^. However, other reports have suggested that copper suppresses inflammation and improves symptoms associated with chronic inflammatory diseases^42,43^. Therefore, exactly how copper regulates innate and microglial immunity is still unclear.

In the current study, we investigated the roles of copper in microglial cells using an *in vitro* system and found that both high and low levels of copper suppressed the transcription of proinflammatory mediators upon short-term lipopolysaccharide (LPS) treatment, as determined by RT-qPCR. We then performed bulk RNA-sequencing (RNA-seq) to better understand copper-mediated regulation of microglial immunity. Our results showed that excessive amounts of copper reduced the transcription of proinflammatory genes, while copper depletion attenuated genes involved in autophagy. These findings suggest that copper mediates essential immune functions in microglial cells *in vitro*. They also indicate that a more mechanistic understanding of copper-mediated regulation of microglial immunity may provide important insights into the development and progression of copper dysregulation-associated neuroinflammatory diseases.

## RESULTS

### Decreasing copper levels reduces LPS-mediated inflammation

In order to study the impact of copper levels on microglial inflammation, we used an *in vitro* model consisting of SIM-A9 cells, a mouse microglial cell line, that were stimulated by lipopolysaccharide (LPS), a Toll-like receptor (TLR)4 ligand that induces an innate immune response characterized by the production of proinflammatory genes^44,45^. We first treated SIM-A9 cells with the copper chelator tetrathiomolybdate (TTM)^46,47^ for 1 hour to decrease copper levels, and then stimulated them with LPS for 6 hours (Figure 1A). We subsequently measured the induction of the proinflammatory genes interleukin-1 beta (*Il-1β*; Figure 1B), interleukin-6 (*Il-6*; Figure 1C), and inducible nitric oxide synthase (*Nos2*; Figure 1D) using RT-qPCR analysis. We found that the LPS-induced expression of these proinflammatory mRNAs was repressed by pretreatment with TTM. Since the duration of the TTM effect *in vitro* is unknown, we also sought to establish a line of SIM-A9 cells with reduced levels of intracellular copper. To generate these SIM-A9 cells, we targeted *Slc31a* for knockdown in order to decrease CTR1 numbers and ultimately decrease copper import into cells (Supplemental Figure 1) ^12,13^. We used two different doxycycline-inducible short-hairpin RNAs (shRNAs) to reduce the expression of *Slc31a1* in SIM-A9 cells, *Slc31a1* knockdown 1 (*Slc31a1*^KD^^1^) and *Slc31a1* knockdown 2 (*Slc31a1*^KD2^), and a non-targeting control. After confirming the lentivirus-mediated transduction of the shRNAs into the cells, we treated them with doxycycline (Dox) for 48 hours to reduce the expression of *Slc31a1*, followed by LPS stimulation for 6 hours to test the induction of proinflammatory mediators (Supplemental Figure 2A). As shown in Supplemental Figure 2B, we determined that both shRNAs significantly reduced the mRNA levels of *Slc31a1* compared to the control. We found that reduced expression of *Slc31a1* resulted in significantly decreased expression levels of *Il-1β* (Supplemental Figure 2C), *Il-6* (Supplemental Figure 2D), and *Nos2* (Supplemental Figure 2E). These data suggest that decreasing the concentration of copper in cells represses the induction of proinflammatory mediators upon LPS stimulation.

**Figure 1.**
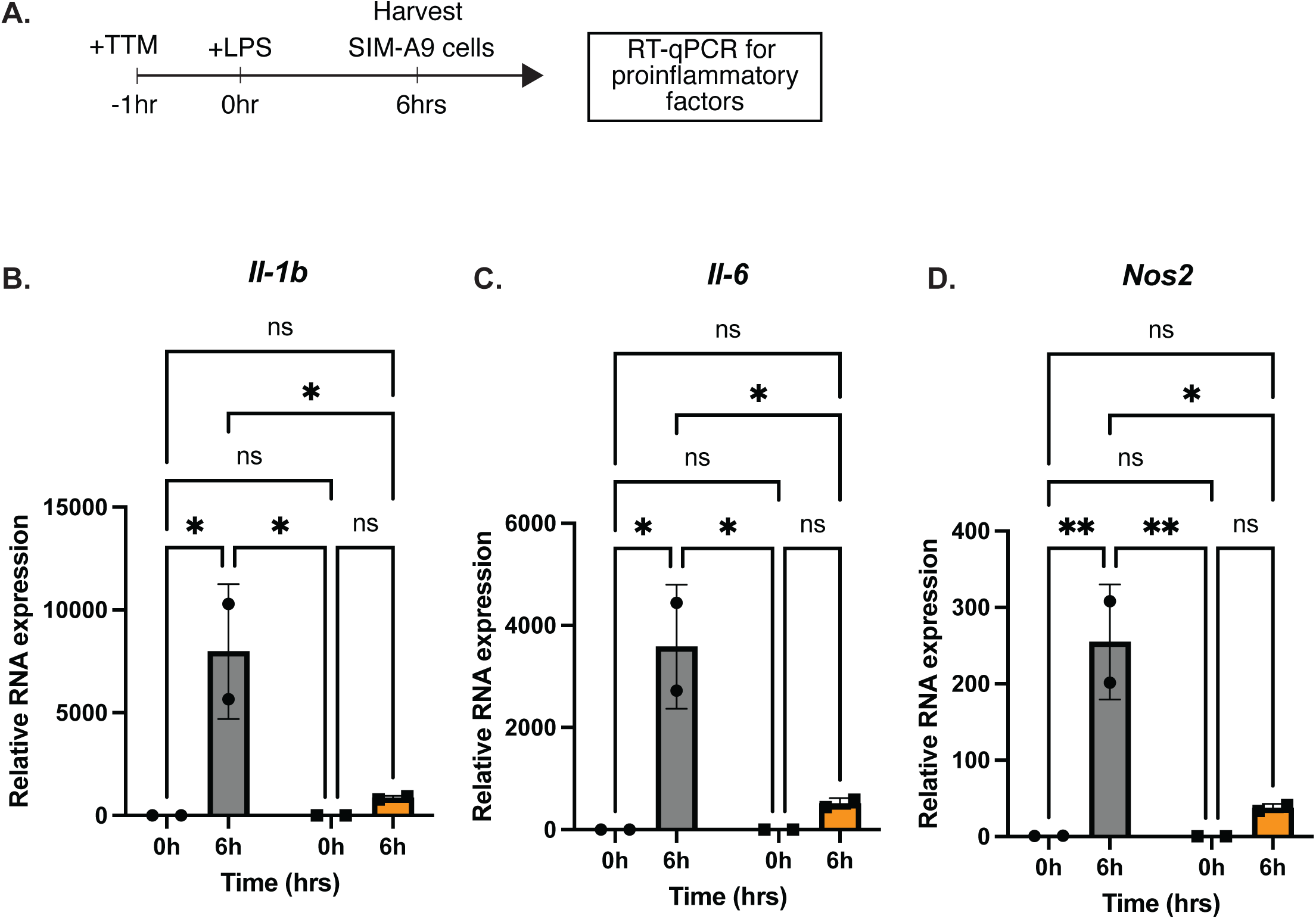
TTM-treated SIM-A9 cells (decreased copper) show reduced transcription of proinflammatory factors. **A)** Experimental scheme is shown. mRNA expressions of **B)** *Il-1β*, **C)** *Il-6*, and **D)** *Nos2* were determined by RT-qPCR. Gray bars are representative of control treatment (PBS) and orange bars are representative of TTM treatment. Data shown is representative of three biological replicates. Results are shown as averages ± standard deviation (SD). Data were analyzed using two-way ANOVA with Tukey’s multiple comparison test. \**p*<0.05, ** *p*<0.01, and ns: not significant.

### Increasing copper levels reduces LPS-mediated inflammation

Our previous work showed that microglial cells *in vitro* accumulate intracellular copper over time upon treatment with LPS or a TLR9 ligand CpG oligodeoxynucleotides^48^. Therefore, we next tested how increasing intracellular copper levels affects LPS-mediated inflammatory gene expression in microglia. For these experiments, SIM-A9 cells were pre-treated with diacetyl bis *(N*(4)-methylthiosemicarbazonato)-Cu^II^ (Cu^II^(atsm))^49^ for 1 hour to increase the copper concentration inside the cells, followed by treatment with LPS for 6 hours, and then RT-qPCR analysis to quantify the mRNA expression of proinflammatory genes (Figure 2A). We found that the LPS-induced expressions of *Il-1β* (Figure 2B), *Il-6* (Figure 2C), and *Nos2* (Figure 2D) mRNAs were significantly repressed when the cells were pretreated with Cu^II^(atsm). We also observed that the induction of interleukin-1 alpha (*Il-1α)* expression was repressed by Cu^II^(atsm) (Supplemental Figure 3A). Interestingly, the expression of tumor necrosis factor alpha (*Tnfα*) mRNA was not altered (Supplemental Figure 3B), suggesting that in our experimental setting, Cu^II^(atsm) suppresses the LPS-mediated induction of some pro-inflammatory mediators, but not all. In addition, we attempted to establish SIM-A9 cell lines wherein *Atp7a*, which encodes a transporter that exports copper from cells (Supplemental Figure 1)^24^, was knocked-down in order to increase intracellular copper concentrations. We attempted to use two different doxycycline-inducible short-hairpin RNAs (shRNAs) to reduce the expression of *Atp7a* in SIM-A9 cells, *Atp7a* knockdown 1 (*Atp7a* ^KD1^) and *Atp7a* knockdown 2 (*Atp7a* ^KD2^), and a non-targeting control. However, we were unable to establish cell lines with sufficient knockdown of this gene, as neither cell line had 50% or less reduced expression of *Atp7a* compared to the control, so we could not replicate the results of our Cu^II^(atsm) experiments using this methodology (Supplemental Figure 4).

**Figure 2.**
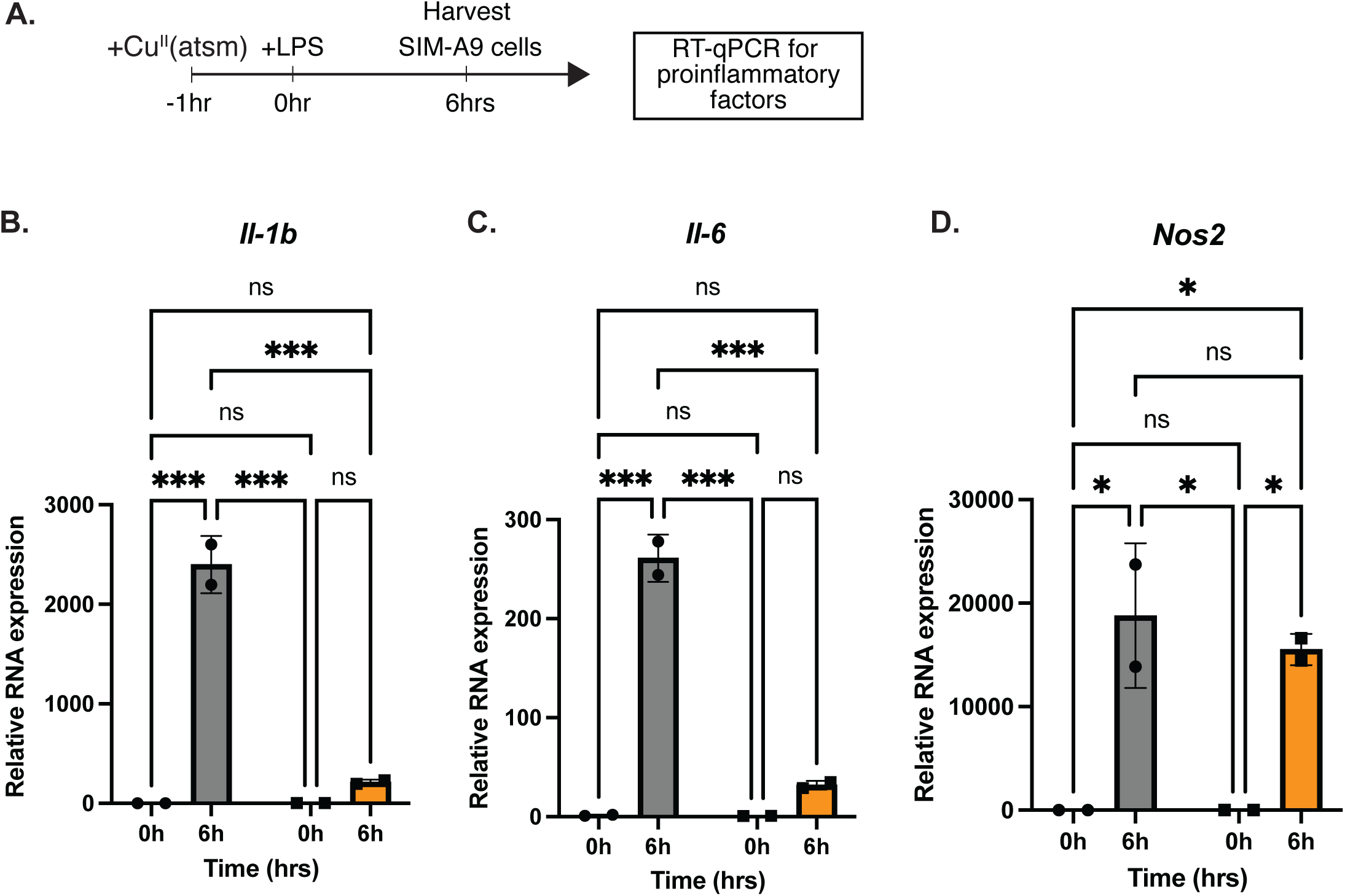
Cu^II^(atsm)-treated SIM-A9 cells (increased copper) show repressed proinflammatory gene expression. **A)** Experimental scheme is shown. mRNA expressions of **B)** *Il-1β*, **C)** *Il-6*, and **D)** *Nos*2 were determined by RT-qPCR. Gray bars are representative of control treatment (DMSO) and orange bars are representative of Cu^II^(atsm) treatment. Data shown is representative of three biological replicates. Results are shown as averages ± standard deviation (SD). Data were analyzed using two-way ANOVA with Tukey’s multiple comparison test. \**p*<0.05, *** *p*<0.001, and ns: not significant.

### RNA-sequencing analysis of the copper-mediated modulation of LPS responses

Since we found that both increasing and decreasing the levels of copper in cells suppresses the induction of proinflammatory mediators upon LPS stimulation, we decided to perform bulk RNA-sequencing (RNA-seq) to examine global gene expression changes mediated by copper during microglial inflammation. For these studies, we treated SIM-A9 cells with Cu^II^(atsm) or TTM for 1 hour, followed by 12 hours of LPS stimulation. We chose to treat the cells with LPS for 12 hours before harvesting in order to capture more changes in LPS-mediated gene expression. For the SIM-A9 cells treated with Cu^II^(atsm), we identified 118 genes that were significantly upregulated and 73 genes that were significantly downregulated at a threshold of p adjusted < 0.05 (Supplemental Table 1), as indicated in the volcano plot in Figure 3A. We confirmed that Cu^II^(atsm) suppressed genes, including *Il-1β*, which we also observed in our RT-qPCR analysis. Gene ontology (GO) analysis of these differentially expressed genes (DEGs) indicated that the downregulated genes were enriched in immune response and interferon-beta responsive genes (Figure 3B). We did not identify any significant terms in upregulated genes by GO analysis, however nitric oxide signaling was slightly activated (Supplemental Table 1). The pretreatment of cells with TTM indicated that 1005 genes were significantly upregulated, and 638 genes were significantly downregulated at a threshold of p adjusted < 0.05 and log2 fold change greater than 1 or less than -1 respectively, as indicated in the volcano plot (Figure 3C). This includes the altered expression of autophagosome regulation genes, which are LPS targets (Supplemental Table 2). Unlike our RT-qPCR results, we did not detect any inflammatory genes that were significantly repressed by TTM with 12 hours of LPS treatment. Subsequent GO analysis showed that genes that were upregulated with TTM pretreatment included ones that regulate transcription and autophagosomes (Figure 3D). Pretreatment with TTM also significantly repressed genes involved in DNA replication, proliferation, and energy generation (Figure 3D). Overall, our RNA-seq results showed that increased copper concentrations repress the expression of proinflammatory mediators, while decreased copper concentrations activate autophagosomes and repress cell proliferation.

**Figure 3.**
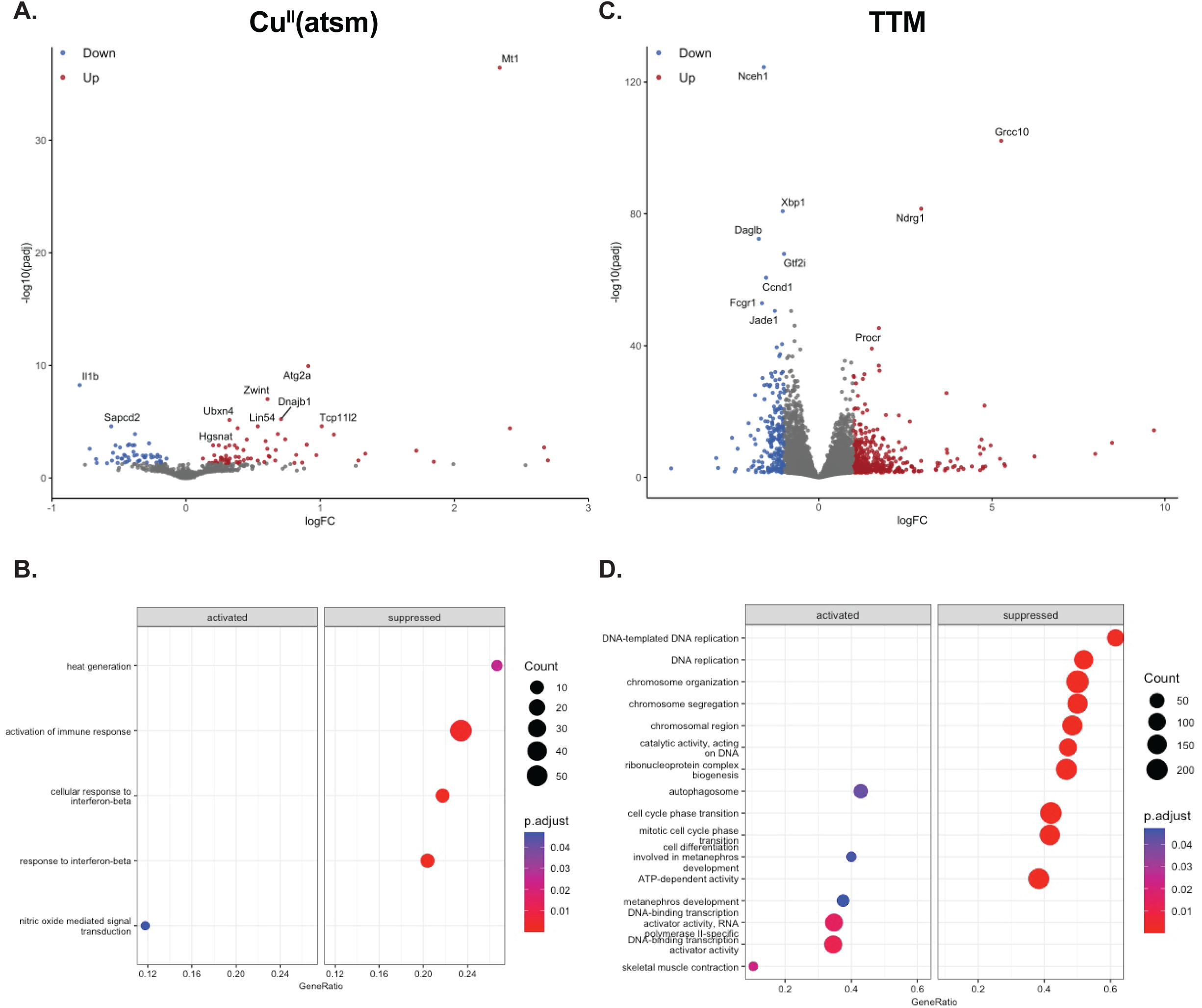
Increased and decreased copper concentrations regulate distinctive subsets of genes upon LPS stimulation. **A)** Volcano plot shown indicates differentially expressed genes (DEGs) in samples treated with 1 µM Cu^II^ (atsm) and LPS for 12 hours compared to vehicle control (DMSO) samples treated with LPS for 12 hours. Right side indicates increased expression and left side indicates decreased expression. Red and blue circles indicate adjusted *p*-values (*p-adj*) less than or equal to 0.05 and log_2F_C values either greater than 1 or less than -1. The gray circles are not significant. **B)** GO analysis of the DEGs identified in A is shown. **C)** Volcano plot shown indicates DEGs in samples treated with 20 µM TTM and LPS for 12 hours compared to vehicle control samples treated with LPS for 12 hours. Right side indicates increased expression and left side indicates decreased expression. Red and blue circles indicate *p-adj* less than or equal to 0.05 and log_2_FC values either greater than 1 or less than -1. The gray circles are not significant. **D)** GO analysis of the DEGs identified in C is shown.

### Copper regulates critical immune functions induced by LPS

To better understand LPS-mediated immune functions impacted by increased and decreased copper concentrations, we directly compared the expression of genes in the Cu^II^(atsm) condition relative to the TTM condition (in other words, we used the TTM treatment as the reference level for differential gene expression analysis) (Supplemental Table 3). The left side of the volcano plot in Figure 4A indicates the DEGs whose expressions were decreased by Cu^II^(atsm) relative to TTM, and the right side of the plot indicates the DEGs whose expressions were increased by Cu^II^(atsm) relative to TTM at a threshold of p adjusted < 0.05 and log2 fold change greater than -1 or less than 1, respectively. GO analysis of these DEGs suggested that Cu^II^(atsm) activates catabolism and peroxisomes compared to TTM (Figure 4B). In addition, we found that Cu^II^(atsm) represses interferon-beta responsive genes, cytokine signaling, and phagocytosis compared to TTM (Figure 4B). These data suggest that both increased and decreased copper concentrations attenuate critical microglial immune functions *in vitro*.

**Figure 4.**
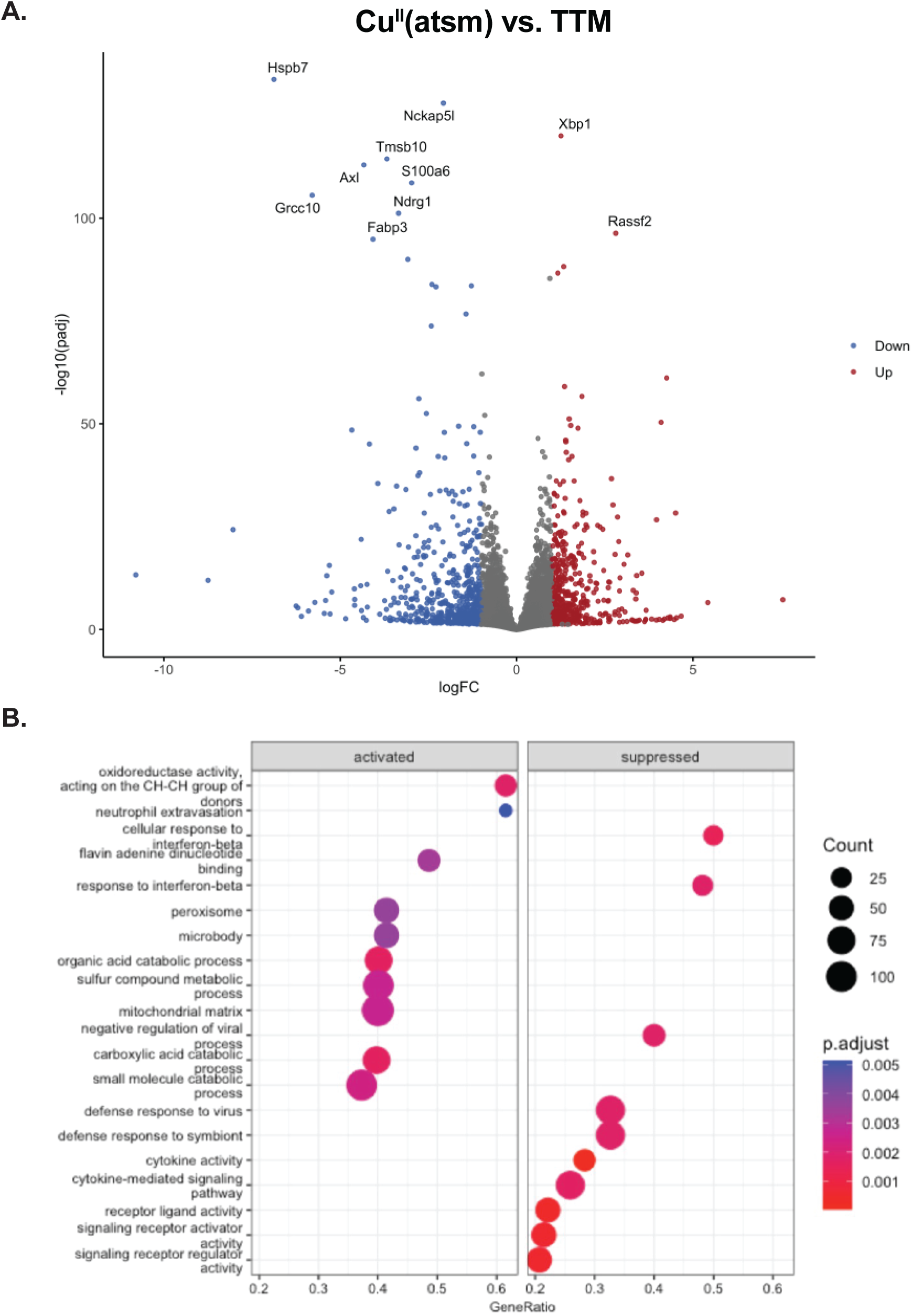
Copper regulates various immune functions in microglial cells. **A)** Volcano plots indicating the DEGs identified in samples treated with 1 µM Cu^II^ (atsm) and 12 hours of LPS compared to those treated with 20 µM TTM and 12 hours of LPS are shown. The right side indicates DEGs whose expressions were increased by Cu^II^(atsm) compared to TTM and the left side indicates DEGs whose expressions were decreased by Cu^II^(atsm) compared to TTM, Red and blue circles indicate *p-adj* less than or equal to 0.05 and log_2_FC values either greater than 1 or less than -1. The gray circles are not significant. **B)** GO analysis of the DEGs identified in A is shown.

## DISCUSSION

Copper dysregulation often contributes to the onset and progression of neurological disorders, including not only congenital diseases^6,17,22–24^, but also neurodegenerative diseases^6,22,28–30^. The role of copper as an anti- or proinflammatory agent in innate immune cells, such as macrophages, remains controversial^28,42,50,51^, and even less is known about copper regulation in microglia. Since dysfunctional microglia are linked to neurodegenerative diseases, the current study investigated the roles of copper in microglial immune functions *in vitro*.

Initially, we used RT-qPCR to examine the role of copper in the induction of LPS-induced proinflammatory mediators by treating mouse microglial cell lines with chemicals to either increase (Cu^II^(atsm)) or decrease (TTM) intracellular copper concentrations. Unexpectedly, both conditions significantly repressed the acute induction of proinflammatory LPS target genes. To confirm these results, we attempted to generate cells with low and high levels of copper by knocking down the genes essential for copper import and export, respectively. Although we failed to establish a cell line with sufficient knockdown of *Atp7a* to reduce copper export from the cells (Supplemental Figure 4), we did successfully generate *Slc31a1* knockdown cell lines to reduce copper import. RT-qPCR analysis of the LPS response in these cells revealed data consistent with that of cells treated with the copper chelator TTM (Figure 1; Supplemental Figure 2). Based on our examination of a limited number of LPS target genes, we realized a more global analysis of gene expression was necessary to better understand the roles of copper in microglial immunity.

To examine overall gene expression changes in microglial cells mediated by increased and decreased copper concentrations, we performed bulk RNA-seq. Consistent with our RT-qPCR analysis, our RNA-seq analysis showed that Cu^II^(atsm) treatment, which increases copper levels, repressed the expression of proinflammatory mediators, such as cytokines and interferon-beta responsive genes (Figure 3A and B; Figure 4). TTM treatment did still repress some immune-associated genes, such as *Cx3cr1*, *Cd38*, and *Fcgr1* (Supplemental Table 2), that mediate migration and phagocytosis^52–54^); however, it did not drastically change the expression levels of these genes. Instead, we found that TTM significantly suppressed the expression of genes regulating cell proliferation and activated the expression of genes involved in transcription and autophagosome formation (Figure 3C and D; Figure 4). Thus, copper is necessary for the normal function of the innate immune system, and dysregulation of cellular copper levels disrupts microglial homeostasis.

To explain the discrepancies between the results of the RT-qPCR and bulk RNA-seq analyses in copper-deficient cells, we considered several possibilities. Although 6 hours of LPS treatment can be a good time point to detect the induction of many proinflammatory genes, 12 hours of LPS treatment might maximize or saturate the induction of LPS-target genes. Some genes require a longer time to express or increase their expression levels; thus, 12 hours of LPS treatment may increase the chance of detecting more pro-inflammatory genes. However, it is also possible that at this later time point, some inflammatory genes may have started to be downregulated because of the termination of the inflammatory response in microglial cells. Thus, a precise time course analysis of the gene expression in microglial cells may help us account for the differences we observed between the RT-qPCR and bulk RNA-seq results. Additionally, microglia stimulated by LPS are known to accumulate copper^48^; therefore, chemical treatments may attenuate the effects of LPS. Since we failed to establish cell lines in which we could increase intracellular copper concentrations, we may need to try other methods, such as CRISPRi, in order to generate efficient knockouts of copper transporters.

Among the many copper-mediated functions, copper-mediated redox regulation is particularly interesting in terms of its physiopathology in humans. Copper is involved in generating ROS and RNS^55–58^. Since dysregulations in ROS and RNS were often associated with neurodegenerative diseases, copper dyshomeostasis can greatly contribute to the pathogenesis of neurodegenerative diseases, such as Parkinson’s disease and Alzheimer’s disease. Consequently, copper dysregulation has been reported in these diseases and restoration of copper homeostasis may help improve the symptoms of neurodegenerative diseases. Interestingly, these diseases are also associated with chronic neuroinflammation. Therefore, it is possible that dysregulated copper in microglia could be a contributor to this pathology. In addition, copper has been directly associated with another neurodegenerative disease, amyotrophic lateral sclerosis (ALS)^59,60^. Copper levels and expression levels of copper associated proteins were shown to be altered in spinal cords of patients with sporadic ALS. Thus, Cu^II^(atsm) is currently in phase 2/3 of clinical trials for treatment of this form of ALS^61–63^.

Overall, our results indicate that copper regulates various aspects of microglial immune function. This work can be used to understand how dysregulation of copper homeostasis may lead to neurological diseases involving microglia and neuroinflammation. Further studies are needed to help establish treatments for neurodegenerative diseases targeting copper homeostasis.

## MATERIALS AND METHODS

### Cell Culture

SIM-A9, a mouse microglial cell line, was kindly provided by Dr. Nagamoto-Combs (10.1016/j.jneumeth.2014.05.021.) and maintained in DMEM/F12 medium (Corning) supplemented with 10% fetal bovine serum (FBS; Hyclone), 5% horse serum (Hyclone), and 1% penicillin-streptomycin (ThermoFisher Scientific).

### Compound treatments

Lipopolysaccharide (LPS; 0.1 µg/ml; Sigma-Aldrich), Cu^II^(atsm) (1 µM; Sigma-Aldrich), and TTM (20 µM; Sigma-Aldrich) were used to treat SIM-A9 cells, as indicated in the figure legends. For controls, we used the solvents for the drugs: DMSO (Siga-Aldrich) for Cu^II^(atsm) and DPBS (ThermoFisher) for TTM.

### RNA-extraction and RT-qPCR

Total RNA was isolated using the Quick-RNA Miniprep Kit (Zymo Research). Complementary DNA (cDNA) was synthesized using the SuperScript III Kit (ThermoFisher). Quantitative PCR (qPCR) was performed using SYBR Green Master Mix (Roche) on the QuantStudio6 Real-Time PCR System (ThermoFisher). Primers for the RT-qPCR were obtained from PrimerBank-MGH-PGA, and their sequences are summarized in Supplemental Table 4. RT-qPCR assays were performed with biological triplicates, and the results shown are representative data. Cycle threshold (Ct) values of the target genes were normalized to the housekeeping gene hypoxanthine phosphoribosyltransferase (*Hprt*), and fold changes were calculated using the ΔΔCt method. Data shown are averages of technical duplicates ± standard deviation (SD).

### Bulk RNA-sequencing

SIM-A9 cells were treated with compounds and LPS, as shown in Figure 1A and 2A, and RNAs were extracted, as described above. RNA-seq libraries were prepared and sequenced using NovaSeq PE 150 by Novogene.

### Bioinformatic analysis

Bioinformatic analysis of the RNA-seq data was performed as described in Goto *et al*^64^. Briefly, reads were trimmed with Cutadapt 3.4 to remove the 3’ adapter sequences. Quality control was then performed using FastQC 0.11.7. Reads were aligned to the GRCm38-mm10 *mus musculus* reference genome using Spliced Transcripts alignment to a Reference aligner 2.7.1a. TPMCalculator 0.0.4 was used to generate count data, and we performed gene expression analysis with DEseq2 1.32.0. Genes with a false discovery rate (FDR) < 0.1 were considered differentially expressed. Gene set enrichment analysis (GSEA) was performed on the sets of differentially expressed genes to determine enriched biological themes. Volcano plots were generated, and the GO analyses were performed using RStudio (R 3.6.0+).

### Statistical analysis

Data from the RT-qPCR assays were analyzed using two-way ANOVA with Tukey’s multiple comparison test as a post-hoc analysis using Prism 10 software (GraphPad). The results of the Tukey’s multiple comparison tests are shown in the figures, with *p* < 0.05 considered to be significant.

## Supporting information

Supplemental information, figure compressed, table 1-5

## DATA AVAILABILITY

RNA-seq data will be posted on the GEO.

## AUTHOR CONTRIBUTION

L.C. designed and performed the experiments and analyzed data. S.E.M. performed bioinformatical analysis. K.S. conceived the project and wrote the manuscript with inputs from all authors.

## ACKNOWLEDGEMENTS

We thank Joseph Moreno and Cindy Wong for assisting with the experiments and analyzing data. We greatly appreciate Dr. Chris Chang’s lab, especially Drs. Chris Chang, Tong Xiao, Aidan Pezacki, and Lakshmipriya Krishnamoorthy, for the reagents and helpful discussions. The authors also thank Drs. Christian Schemdt for designing shRNAs and Amy Sullivan for editing the manuscript. This work was supported by the U.C. Dissertation Year Fellowship for L.C. and, NIH R21AG073735 and R21HD107388 for K.S.

## Notes

### Competing Interest Statement

The authors have declared no competing interest.

