## Supplemental information, figure compressed, table 1-5 for "Role of copper during microglial inflammation": Supplemental information-r3.pdf

### SUPPLEMETAL INFORMATION

### SUPPLEMENTAL FIGURE LEGENDS

**Supplemental Figure 1.** Schematic of copper transporters in microglia. Copper enters cells mainly through copper importer Ctr1. Atp7a is a copper exporter that is responsible for incorporating copper onto proteins at the Golgi apparatus or for exporting copper through the plasma membrane. Other cell types use Atb7b. These copper importers can also be found on the cellular membrane.

**Supplemental Figure 2. *Slc31a1* knockdown SIM-A9 cells show decreased expression of proinflammatory genes. A)** Experimental scheme is shown. **B)** shRNA knockdown efficiency shown by measuring *Slc31a1* mRNA levels in control (white bar), $Slc31a1^{KD1}$ (light blue bar) and $Slc31a1^{KD2}$ (dark blue bar) SIM-A9 cells lines. mRNA expression of **C)** *Il-1β,* **D)** *Il-6* and **E)** *iNOS* in SIM-A9 cell lines in response to LPS treatment. Data shown is representative of three biological replicates. Results are shown as averages ± standard deviation (SD). Data were analyzed using two-way ANOVA with Tukey's multiple comparison test. *$p$<0.05, ** $p$<0.01, *** $p$<0.001, ****$p$<0.0001, and ns: not significant.

**Supplemental Figure 3. $Cu^{II}$(atsm)- treated SIM-A9 cells inducing repression in additional proinflammatory genes.** mRNA expressions of **A)** *Il-1α* and **B)** *Tnfα* were determined by RT-qPCR. Gray bars are representative of control treatment (DMSO) and orange bars are representative of $Cu^{II}$(atsm) treatment. Data shown is representative of three biological replicates. Results are shown as averages ± standard deviation (SD). Data were analyzed using two-way ANOVA with Tukey's multiple comparison test. *$p$<0.05, *** $p$<0.001, and ns: not significant.

**Supplemental Figure 4.** shRNA knockdown efficiency shown by measuring *Atp7a* mRNA levels in control (white bar), *Atp7a*$^{KD1}$ (light blue bar) and *Atp7a*$^{KD2}$ (dark blue bar) SIM-A9 cells lines. Two-way ANOVA followed by Tukey's multiple comparisons. All data are mean ± SD. $^{*}p<0.05$, ns: not significant.

### SUPPLEMENTAL METHODS AND MATIERALS

#### shRNA preparation

Tet-on inducible lentivirus vector pTRIPZ (Dharmacon) was used to generated for knockdown *Slc31a1* (*Ctr1*) gene. We used two different shRNA sequences and used luciferase sequence as a non-targeting control. Sequences for shRNA were listed in the Supplemental Table 5. Lentiviruses were packaged using co-transfection vectors including pCMV-VSV-G (Addgene plasmid #8454) and pCMV-dR8.2 (Addgene plasmid #8455) into HEK 293T cells using Lipofectamine 3000 reagent (Life Technologies) according to the manufacturer's instructions. Viral supernatant was harvested after 48 hours and added to SIM-A9 cells. 48 hours post-transduction, puromycin (8 µg/ml) was added to select for transduced cells. Successful transduction was checked after 24~48 hours treatment of 0.5 µg/ml Doxyxyxline using 1) presence of RFP+ cells via FACS on an Aria Fusion (BD Biosciences, UC Berkeley Cancer Research Laboratory) and 2) percent knockdown determined via RT-qPCR. Following confirmation, each cell line was expanded and subcultured for immune stimulation experiments, as described.
