## Supplemental information, figure compressed, table 1-5 for "Role of copper during microglial inflammation": Supplemental Table 1-compressed.pdf

| Gene_Id | baseMean | log2FoldChar | lfcSE | pvalue | padj | external_gene |
|---|---|---|---|---|---|---|
| ENSMUSG00( | 49.2832222 | 9.6829582 | 1.29250735 | 1.00E-16 | 5.28E-15 | |
| ENSMUSG00( | 20.4914399 | 8.47706557 | 1.35189417 | 1.07E-12 | 3.18E-11 | Gm24407 |
| ENSMUSG00( | 6.00297631 | 7.98712439 | 2.40735197 | 4.03E-09 | 6.85E-08 | Gm26316 |
| ENSMUSG00( | 9.92069839 | 6.22303154 | 1.14568657 | 2.86E-08 | 4.10E-07 | Gm23971 |
| ENSMUSG00( | 3.82095865 | 5.38272113 | 1.62979469 | 4.42E-05 | 3.17E-04 | Gm25099 |
| ENSMUSG00( | 9.05335719 | 5.35999158 | 2.10611056 | 1.15E-05 | 9.38E-05 | Vipr2 |
| ENSMUSG00( | 144.76251 | 5.27023126 | 0.2404979 | 8.19E-107 | 6.86E-103 | Grcc10 |
| ENSMUSG00( | 7.67095727 | 5.23626343 | 1.07282633 | 1.79E-07 | 2.14E-06 | Snord118 |
| ENSMUSG00( | 10.8293119 | 4.96198319 | 0.74185958 | 6.91E-12 | 1.85E-10 | Gm22513 |
| ENSMUSG00( | 5.04313252 | 4.80590653 | 1.50972016 | 6.94E-05 | 4.72E-04 | |
| ENSMUSG00( | 28.6627552 | 4.78148135 | 0.47462438 | 9.95E-25 | 1.62E-22 | B930036N10I |
| ENSMUSG00( | 10.8932933 | 4.75843316 | 0.77341427 | 9.77E-11 | 2.18E-09 | Rnu2-10 |
| ENSMUSG00( | 3.98580401 | 4.73586928 | 2.14645915 | 8.92E-05 | 5.91E-04 | 2310016D23I |
| ENSMUSG00( | 2.34478118 | 4.73216839 | 2.40486647 | 3.38E-04 | 0.00191888 | Gm43577 |
| ENSMUSG00( | 13.4700385 | 4.69656036 | 0.65134975 | 8.95E-14 | 3.16E-12 | Muc3 |
| ENSMUSG00( | 7.59233705 | 4.66086051 | 0.83763038 | 4.20E-09 | 7.09E-08 | Rnu12 |
| ENSMUSG00( | 3.18604494 | 4.65481123 | 2.280659 | 0.0014186 | 0.00672603 | Gm24830 |
| ENSMUSG00( | 13.653457 | 4.63783569 | 0.72733163 | 1.97E-11 | 5.00E-10 | Gm23849 |
| ENSMUSG00( | 4.33022969 | 4.42535358 | 1.35130475 | 6.76E-05 | 4.62E-04 | Gm47920 |
| ENSMUSG00( | 3.74219327 | 4.18460781 | 1.53316397 | 4.85E-04 | 0.00264089 | Gm13836 |
| ENSMUSG00( | 4.37788333 | 4.11602263 | 1.23535657 | 1.31E-04 | 8.30E-04 | |
| ENSMUSG00( | 2.33420655 | 4.03817265 | 2.51883009 | 0.00120828 | 0.00583436 | Gm17597 |
| ENSMUSG00( | 16.7112159 | 4.00858754 | 0.81551733 | 2.08E-07 | 2.46E-06 | Sstr5 |
| ENSMUSG00( | 8.47988092 | 3.9343243 | 1.01995616 | 4.67E-05 | 3.32E-04 | Ccdc17 |
| ENSMUSG00( | 3.38890975 | 3.88242458 | 1.21034256 | 1.20E-04 | 7.69E-04 | Gm22265 |
| ENSMUSG00( | 18.6181038 | 3.80806058 | 0.89486949 | 1.21E-06 | 1.22E-05 | Arl4d |
| ENSMUSG00( | 4.70213295 | 3.71936903 | 1.20604707 | 3.00E-04 | 0.00173008 | Gm47271 |
| ENSMUSG00( | 18.2609247 | 3.70982853 | 0.67390504 | 2.10E-09 | 3.72E-08 | |
| ENSMUSG00( | 9.73663889 | 3.69923831 | 0.61151458 | 1.55E-10 | 3.35E-09 | Rnu5g |
| ENSMUSG00( | 80.0651833 | 3.6894914 | 0.33934635 | 1.02E-28 | 2.26E-26 | Rdm1 |
| ENSMUSG00( | 3.18872353 | 3.65444022 | 1.29681433 | 5.26E-04 | 0.00283876 | Tmem82 |
| ENSMUSG00( | 11.0617081 | 3.64521015 | 2.2946963 | 0.00194254 | 0.00881137 | Gm25360 |
| ENSMUSG00( | 5.86367263 | 3.48489175 | 0.89222851 | 6.51E-06 | 5.65E-05 | 2900060B14I |
| ENSMUSG00( | 4.33610158 | 3.43060485 | 0.99417582 | 5.20E-05 | 3.66E-04 | Rpl31-ps17 |
| ENSMUSG00( | 7.12607925 | 3.42145613 | 0.86045506 | 4.43E-06 | 3.99E-05 | Rnu1a1 |
| ENSMUSG00( | 3.2798158 | 3.38476879 | 1.5244653 | 0.00122847 | 0.00591989 | Gm23444 |
| ENSMUSG00( | 3.49950583 | 3.36959472 | 1.33993044 | 6.38E-04 | 0.00335936 | Gm22973 |
| ENSMUSG00( | 2.62904709 | 2.96603929 | 1.34191552 | 0.00234963 | 0.01034142 | Gm7527 |
| ENSMUSG00( | 281.856925 | 2.95912346 | 0.15159265 | 5.06E-86 | 2.83E-82 | Ndrg1 |
| ENSMUSG00( | 15.8178141 | 2.95808086 | 0.66961278 | 9.22E-07 | 9.61E-06 | Myh3 |

| | | | | | | |
|---|---|---|---|---|---|---|
| ENSMUSG00( | 2.22618388 | 2.90170936 | 1.83923338 | 0.00398053 | 0.01623689 | Ankrd66 |
| ENSMUSG00( | 2.55023084 | 2.8772361 | 1.39184636 | 0.00283607 | 0.0122134 | Gm26616 |
| ENSMUSG00( | 13.3227788 | 2.76589992 | 0.97790385 | 5.10E-04 | 0.00276178 | Plcd1 |
| ENSMUSG00( | 2.31336279 | 2.75632272 | 1.7958883 | 0.00468163 | 0.01864754 | Gm24497 |
| ENSMUSG00( | 3.73615545 | 2.74509183 | 1.54643137 | 0.00317218 | 0.01348076 | Oscar |
| ENSMUSG00( | 4.49251875 | 2.7055965 | 0.84784887 | 1.04E-04 | 6.77E-04 | |
| ENSMUSG00( | 6.18452756 | 2.684109 | 0.79129975 | 5.46E-05 | 3.81E-04 | |
| ENSMUSG00( | 15.0381179 | 2.68133678 | 0.83017645 | 8.45E-05 | 5.63E-04 | A530064D06I |
| ENSMUSG00( | 45.7512785 | 2.63546967 | 0.30110827 | 1.44E-19 | 1.14E-17 | Gm2774 |
| ENSMUSG00( | 21.0314542 | 2.56140709 | 0.63347914 | 2.61E-06 | 2.47E-05 | Gnat3 |
| ENSMUSG00( | 13.455433 | 2.53381433 | 0.95610498 | 9.23E-04 | 0.00462981 | Gm10384 |
| ENSMUSG00( | 18.3911973 | 2.51431809 | 0.6314768 | 4.60E-06 | 4.12E-05 | Trp53inp1 |
| ENSMUSG00( | 31.2903564 | 2.50662617 | 0.40785931 | 4.67E-11 | 1.11E-09 | Rgs9 |
| ENSMUSG00( | 8.48893027 | 2.44426713 | 0.62294386 | 5.53E-06 | 4.87E-05 | Slc24a1 |
| ENSMUSG00( | 3.85529565 | 2.43671765 | 1.49627196 | 0.00457632 | 0.01831627 | Gm44424 |
| ENSMUSG00( | 2.34452361 | 2.41730282 | 3.14342234 | 0.00608029 | 0.02319893 | Gm10642 |
| ENSMUSG00( | 16.4376089 | 2.40238527 | 0.40481824 | 1.92E-10 | 4.09E-09 | Cyp1a2 |
| ENSMUSG00( | 4.10250986 | 2.4023652 | 1.4000003 | 0.00249581 | 0.01093343 | Gm4610 |
| ENSMUSG00( | 39.5018444 | 2.37382627 | 0.49180494 | 1.07E-07 | 1.34E-06 | Klf2 |
| ENSMUSG00( | 3.71940066 | 2.3737063 | 1.24192211 | 0.00221782 | 0.00985468 | Gm25280 |
| ENSMUSG00( | 21.6752289 | 2.34174761 | 0.50580808 | 2.15E-07 | 2.53E-06 | Tmem179 |
| ENSMUSG00( | 6.0147848 | 2.33750923 | 0.87795957 | 4.54E-04 | 0.00249333 | BC030343 |
| ENSMUSG00( | 4.02127928 | 2.33235967 | 0.98818644 | 9.73E-04 | 0.00483947 | |
| ENSMUSG00( | 5.06912872 | 2.33172405 | 1.43665239 | 0.00593207 | 0.02271613 | Gm37653 |
| ENSMUSG00( | 3.46679258 | 2.32762858 | 1.03669362 | 0.00148142 | 0.00697444 | Gm13783 |
| ENSMUSG00( | 4.06823044 | 2.31843465 | 1.16543867 | 0.00283528 | 0.01221312 | Tmem274 |
| ENSMUSG00( | 8.37302565 | 2.31426244 | 0.78469954 | 1.67E-04 | 0.00103085 | Ppbp |
| ENSMUSG00( | 36.7041716 | 2.31402023 | 0.33912693 | 6.27E-13 | 1.95E-11 | AV356131 |
| ENSMUSG00( | 39.9028581 | 2.31150531 | 0.24944449 | 1.43E-21 | 1.58E-19 | |
| ENSMUSG00( | 4.25926637 | 2.27519639 | 0.82916571 | 3.25E-04 | 0.00185552 | C630004M23 |
| ENSMUSG00( | 3.723935 | 2.27343447 | 1.29907289 | 0.00259546 | 0.01133774 | 1700015I17Ri |
| ENSMUSG00( | 3.68911567 | 2.24311079 | 1.05645478 | 0.00188181 | 0.00857766 | Gpr61 |
| ENSMUSG00( | 8.57044092 | 2.24291156 | 0.9759901 | 0.00140393 | 0.00666579 | Gm43127 |
| ENSMUSG00( | 10.8609295 | 2.2213964 | 0.52829657 | 1.56E-06 | 1.54E-05 | Xcr1 |
| ENSMUSG00( | 3.56810074 | 2.21236436 | 1.31212468 | 0.00334982 | 0.01409279 | Prss22 |
| ENSMUSG00( | 2.59567415 | 2.19516904 | 1.67601765 | 0.00746506 | 0.02757252 | Gm2395 |
| ENSMUSG00( | 5.31300055 | 2.18482586 | 1.2873598 | 0.00639215 | 0.02417971 | Gm34256 |
| ENSMUSG00( | 20.9536808 | 2.15965676 | 0.44593571 | 8.60E-08 | 1.10E-06 | 2210417A02I |
| ENSMUSG00( | 27.3010827 | 2.15483027 | 0.36917653 | 3.17E-10 | 6.50E-09 | Chd5 |
| ENSMUSG00( | 36.6870974 | 2.13800866 | 0.45366182 | 1.30E-07 | 1.60E-06 | B330016D10I |
| ENSMUSG00( | 8.0908135 | 2.11170805 | 0.59370326 | 2.33E-05 | 1.78E-04 | Gm30873 |

| | | | | | | |
|---|---|---|---|---|---|---|
| ENSMUSG00( | 11.7708271 | 2.1063286 | 0.53538475 | 5.71E-06 | 5.01E-05 | Gm49041 |
| ENSMUSG00( | 10.472551 | 2.09797255 | 0.60071573 | 2.67E-05 | 2.01E-04 | Gm47175 |
| ENSMUSG00( | 7.90519534 | 2.09133634 | 0.64766208 | 8.22E-05 | 5.50E-04 | Gm16061 |
| ENSMUSG00( | 10.2747359 | 2.07739732 | 0.6754613 | 1.23E-04 | 7.90E-04 | 6330409D20I |
| ENSMUSG00( | 14.0756791 | 2.07267221 | 0.52792811 | 5.08E-06 | 4.50E-05 | Gm9887 |
| ENSMUSG00( | 4.59593435 | 2.06853271 | 0.94175333 | 0.00158735 | 0.0074025 | Gm44280 |
| ENSMUSG00( | 6.1030919 | 2.04991988 | 0.68122436 | 1.55E-04 | 9.70E-04 | |
| ENSMUSG00( | 12.392277 | 2.04985304 | 0.7747312 | 4.88E-04 | 0.00265685 | Klhl3 |
| ENSMUSG00( | 15.1938851 | 2.04747299 | 0.44060566 | 2.09E-07 | 2.47E-06 | Cstdc4 |
| ENSMUSG00( | 73.2838305 | 2.02992312 | 0.35477317 | 7.38E-10 | 1.42E-08 | Tnks1bp1 |
| ENSMUSG00( | 3.82003622 | 2.02667325 | 1.35096347 | 0.00531582 | 0.02071069 | |
| ENSMUSG00( | 6.0776563 | 2.02164001 | 1.0358492 | 0.00227738 | 0.01008728 | Map1a |
| ENSMUSG00( | 7.52047194 | 2.01020017 | 0.99086648 | 0.00143969 | 0.00680472 | mt-Te |
| ENSMUSG00( | 348.792929 | 2.00921689 | 0.40739897 | 4.49E-08 | 6.11E-07 | Gm49774 |
| ENSMUSG00( | 18.0654445 | 2.00840445 | 0.44199608 | 3.35E-07 | 3.80E-06 | Jph3 |
| ENSMUSG00( | 5.18996467 | 2.00516693 | 0.84963614 | 8.02E-04 | 0.00410564 | Adprhl1 |
| ENSMUSG00( | 15.3972457 | 1.99905664 | 0.59280015 | 3.76E-05 | 2.74E-04 | Samd11 |
| ENSMUSG00( | 61.7549874 | 1.99313479 | 0.31538015 | 1.47E-11 | 3.79E-10 | Cdcp1 |
| ENSMUSG00( | 4.13020562 | 1.97302197 | 1.13756932 | 0.00376112 | 0.0155041 | A930012O16I |
| ENSMUSG00( | 5.93675501 | 1.96646359 | 0.8652956 | 0.00105929 | 0.00521861 | 1700014D04I |
| ENSMUSG00( | 9.56580818 | 1.94448834 | 0.82832897 | 0.00111213 | 0.00543741 | A930024E05I |
| ENSMUSG00( | 106.722441 | 1.93988035 | 0.35800793 | 4.47E-09 | 7.52E-08 | Tsc22d3 |
| ENSMUSG00( | 102.11229 | 1.93540602 | 0.20864505 | 1.08E-21 | 1.20E-19 | Gm5526 |
| ENSMUSG00( | 8.1841196 | 1.90672034 | 1.01420588 | 0.00272521 | 0.01181493 | Gm39473 |
| ENSMUSG00( | 26.3967611 | 1.90620921 | 0.39068331 | 6.32E-08 | 8.33E-07 | Lncpint |
| ENSMUSG00( | 13.6554264 | 1.90530844 | 0.56216516 | 3.87E-05 | 2.81E-04 | Nqo1 |
| ENSMUSG00( | 7.57711269 | 1.89720456 | 0.77292081 | 7.08E-04 | 0.00369075 | Gm37850 |
| ENSMUSG00( | 21.826918 | 1.87351249 | 0.4541178 | 2.08E-06 | 2.01E-05 | Gdnf |
| ENSMUSG00( | 9.41003692 | 1.86792985 | 0.68579139 | 2.88E-04 | 0.00166806 | Gm12867 |
| ENSMUSG00( | 62.6507769 | 1.86610994 | 0.26192046 | 6.79E-14 | 2.43E-12 | Gpr68 |
| ENSMUSG00( | 5.12486269 | 1.86471939 | 0.73978581 | 5.52E-04 | 0.00295954 | Pira12 |
| ENSMUSG00( | 8.92946049 | 1.85937301 | 0.77948483 | 7.11E-04 | 0.00370224 | Sec16b |
| ENSMUSG00( | 10.8394517 | 1.85835314 | 0.56190281 | 5.00E-05 | 3.54E-04 | Gm5475 |
| ENSMUSG00( | 2.76606514 | 1.85295123 | 2.7181923 | 0.0095014 | 0.03390613 | Gm50203 |
| ENSMUSG00( | 7.24168558 | 1.84821198 | 0.64225125 | 2.33E-04 | 0.0013846 | |
| ENSMUSG00( | 40.9929537 | 1.83986241 | 0.67282233 | 2.47E-04 | 0.00145173 | Egr2 |
| ENSMUSG00( | 36.1786741 | 1.83656609 | 0.25329124 | 2.82E-14 | 1.06E-12 | Prkg2 |
| ENSMUSG00( | 7.57970367 | 1.82686773 | 0.84013222 | 0.00171202 | 0.00789387 | Fam187b |
| ENSMUSG00( | 13.5982855 | 1.82350193 | 0.42713511 | 1.23E-06 | 1.24E-05 | |
| ENSMUSG00( | 15.0405587 | 1.8168629 | 0.4989918 | 1.66E-05 | 1.32E-04 | Catip |
| ENSMUSG00( | 65.7004009 | 1.80901553 | 0.27971396 | 6.29E-12 | 1.70E-10 | Unc13a |

| | | | | | | |
|---|---|---|---|---|---|---|
| ENSMUSG00( | 19.2760278 | 1.80640941 | 0.40100157 | 3.98E-07 | 4.47E-06 | Itgb3 |
| ENSMUSG00( | 10.3842986 | 1.79965156 | 0.64756925 | 2.98E-04 | 0.00171912 | A230103J11R |
| ENSMUSG00( | 2.85056679 | 1.79537528 | 1.79716913 | 0.010774 | 0.03773262 | Gm42576 |
| ENSMUSG00( | 5.70880723 | 1.79110463 | 0.81302025 | 0.00129274 | 0.00619754 | Gm42930 |
| ENSMUSG00( | 13.6334538 | 1.7870696 | 0.50909627 | 2.46E-05 | 1.87E-04 | |
| ENSMUSG00( | 9.21711972 | 1.78160955 | 0.60905146 | 2.05E-04 | 0.00123957 | 5830454E08F |
| ENSMUSG00( | 4.73365417 | 1.77875831 | 0.85894331 | 0.00166927 | 0.00771795 | Camkk1 |
| ENSMUSG00( | 3.85815101 | 1.77778435 | 1.01680229 | 0.00320261 | 0.01358939 | Ly75 |
| ENSMUSG00( | 10.4922916 | 1.7748727 | 0.79128212 | 9.66E-04 | 0.0048107 | Gm43361 |
| ENSMUSG00( | 5.23554079 | 1.76446354 | 1.29114904 | 0.00433051 | 0.01746476 | Gm17056 |
| ENSMUSG00( | 34.2504697 | 1.76280799 | 0.28859771 | 6.71E-11 | 1.54E-09 | Il12rb1 |
| ENSMUSG00( | 7.82269414 | 1.7602062 | 0.73571042 | 8.57E-04 | 0.00435328 | Mgam |
| ENSMUSG00( | 235.792171 | 1.74831454 | 0.14221962 | 6.83E-36 | 4.24E-33 | Piwil2 |
| ENSMUSG00( | 9.14480187 | 1.74362701 | 0.52137838 | 4.76E-05 | 3.38E-04 | A630072M18I |
| ENSMUSG00( | 4680.66242 | 1.73217498 | 0.11896266 | 3.46E-49 | 4.83E-46 | Procr |
| ENSMUSG00( | 617.430774 | 1.72863165 | 0.13744471 | 1.88E-37 | 1.26E-34 | |
| ENSMUSG00( | 35.1256599 | 1.72030795 | 0.32898602 | 9.93E-09 | 1.57E-07 | Smarcd3 |
| ENSMUSG00( | 10.4181382 | 1.71773512 | 0.55034375 | 9.74E-05 | 6.38E-04 | Gm46637 |
| ENSMUSG00( | 49.2614612 | 1.71044356 | 0.38592214 | 5.42E-07 | 5.90E-06 | Vgf |
| ENSMUSG00( | 7.94024168 | 1.70755149 | 0.61095824 | 2.91E-04 | 0.00168147 | |
| ENSMUSG00( | 20.6729394 | 1.70655799 | 0.35476297 | 1.01E-07 | 1.27E-06 | B930059L03F |
| ENSMUSG00( | 11.570161 | 1.70650821 | 0.44024121 | 6.61E-06 | 5.71E-05 | Tnnc1 |
| ENSMUSG00( | 7.65583488 | 1.70563991 | 0.67423449 | 5.37E-04 | 0.0028886 | Gm29367 |
| ENSMUSG00( | 9.30315929 | 1.69981493 | 0.64556429 | 4.53E-04 | 0.00248596 | Gm44389 |
| ENSMUSG00( | 120.073713 | 1.69000378 | 0.20944714 | 4.71E-17 | 2.59E-15 | Plekhg2 |
| ENSMUSG00( | 37.9948787 | 1.67924793 | 0.44759722 | 1.03E-05 | 8.53E-05 | Ifnlr1 |
| ENSMUSG00( | 49.3289584 | 1.6781188 | 0.29622412 | 9.41E-10 | 1.76E-08 | Gm7204 |
| ENSMUSG00( | 24.8672695 | 1.67114965 | 0.55331594 | 1.22E-04 | 7.80E-04 | Gm10382 |
| ENSMUSG00( | 11.3949283 | 1.66315547 | 0.78307101 | 0.00142692 | 0.00675392 | |
| ENSMUSG00( | 28.4296895 | 1.65961331 | 0.35379482 | 1.74E-07 | 2.09E-06 | Gm10252 |
| ENSMUSG00( | 40.1176609 | 1.65920737 | 0.40842283 | 2.68E-06 | 2.53E-05 | Rhpn2 |
| ENSMUSG00( | 5.4478865 | 1.6565026 | 1.213182 | 0.00504131 | 0.01983982 | 1700060J05R |
| ENSMUSG00( | 17.1837146 | 1.6558781 | 0.42957861 | 6.57E-06 | 5.69E-05 | Pira2 |
| ENSMUSG00( | 74.2138913 | 1.65332005 | 0.87414433 | 0.00419753 | 0.01697745 | Mgst1 |
| ENSMUSG00( | 8.98211589 | 1.64547264 | 0.57075177 | 2.18E-04 | 0.00130777 | Gm12500 |
| ENSMUSG00( | 132.245783 | 1.64229907 | 0.25189259 | 4.56E-12 | 1.25E-10 | Ccbe1 |
| ENSMUSG00( | 6.10266733 | 1.62546433 | 0.66914269 | 7.60E-04 | 0.00391473 | Gm1141 |
| ENSMUSG00( | 14.7286692 | 1.61885285 | 0.46652132 | 3.38E-05 | 2.49E-04 | Rnasek |
| ENSMUSG00( | 28.1540583 | 1.61832545 | 0.31137193 | 1.38E-08 | 2.13E-07 | |
| ENSMUSG00( | 14.7478426 | 1.60220841 | 0.47874807 | 4.72E-05 | 3.35E-04 | Slc25a34 |
| ENSMUSG00( | 30.027624 | 1.60064365 | 0.33014186 | 8.91E-08 | 1.13E-06 | 2900093K20F |

| | | | | | | |
|---|---|---|---|---|---|---|
| ENSMUSG00( | 70.2741936 | 1.58778814 | 0.21311311 | 6.34E-15 | 2.64E-13 | Gm39469 |
| ENSMUSG00( | 44.707348 | 1.58636054 | 0.28383864 | 1.55E-09 | 2.80E-08 | Gm10425 |
| ENSMUSG00( | 22.2812522 | 1.58491053 | 0.45257751 | 2.63E-05 | 1.99E-04 | Gm49169 |
| ENSMUSG00( | 146.135429 | 1.5781023 | 0.23096851 | 6.33E-13 | 1.96E-11 | Slc6a9 |
| ENSMUSG00( | 29.0413386 | 1.57611698 | 0.37544726 | 1.61E-06 | 1.58E-05 | Slc6a19 |
| ENSMUSG00( | 79.048181 | 1.57457554 | 0.19654002 | 8.15E-17 | 4.34E-15 | Gm43661 |
| ENSMUSG00( | 15.0428897 | 1.57360829 | 0.65121539 | 8.34E-04 | 0.0042476 | Gm8818 |
| ENSMUSG00( | 83.7974553 | 1.57134323 | 0.21324293 | 1.14E-14 | 4.50E-13 | |
| ENSMUSG00( | 12.0081316 | 1.56970927 | 0.68001212 | 0.00103961 | 0.00512919 | Marchf3 |
| ENSMUSG00( | 6.26445499 | 1.56740811 | 0.7441777 | 0.00163082 | 0.00756103 | |
| ENSMUSG00( | 116.512656 | 1.55189657 | 0.42957613 | 1.59E-05 | 1.26E-04 | Chac1 |
| ENSMUSG00( | 152.388136 | 1.54338463 | 0.2170783 | 8.81E-14 | 3.12E-12 | Tcp11l2 |
| ENSMUSG00( | 8282.78368 | 1.53618065 | 0.26321657 | 3.44E-10 | 7.02E-09 | Srxn1 |
| ENSMUSG00( | 8.17545689 | 1.53616787 | 0.71299951 | 0.00141903 | 0.00672603 | Gm43255 |
| ENSMUSG00( | 31343.9559 | 1.52879761 | 0.11244703 | 7.23E-43 | 7.57E-40 | Prdx1 |
| ENSMUSG00( | 17.7930452 | 1.52328391 | 0.429037 | 2.25E-05 | 1.73E-04 | Gm26811 |
| ENSMUSG00( | 7.85482383 | 1.5219716 | 0.73416152 | 0.00161728 | 0.00751094 | Gm50080 |
| ENSMUSG00( | 11.6553149 | 1.51045181 | 0.60712548 | 6.17E-04 | 0.0032627 | A430108G06I |
| ENSMUSG00( | 81.4977826 | 1.50950173 | 0.61366837 | 5.86E-04 | 0.00311976 | Eid3 |
| ENSMUSG00( | 5.55766054 | 1.5074755 | 1.4806666 | 0.00744119 | 0.0274965 | Gm48372 |
| ENSMUSG00( | 2813.75778 | 1.49820115 | 0.65505208 | 9.66E-04 | 0.00481221 | Mmp13 |
| ENSMUSG00( | 2990.18826 | 1.49820082 | 0.22155309 | 9.17E-13 | 2.76E-11 | Ubc |
| ENSMUSG00( | 13.7668604 | 1.49695372 | 0.50128146 | 1.50E-04 | 9.41E-04 | Pgf |
| ENSMUSG00( | 48.7710652 | 1.49138948 | 0.23483995 | 1.51E-11 | 3.90E-10 | Gm42067 |
| ENSMUSG00( | 12.9865488 | 1.48911613 | 0.46728294 | 9.62E-05 | 6.32E-04 | Gm4285 |
| ENSMUSG00( | 6585.2084 | 1.4767014 | 0.21500141 | 5.82E-13 | 1.82E-11 | Mmp12 |
| ENSMUSG00( | 566.042047 | 1.46946942 | 0.45292874 | 6.27E-05 | 4.32E-04 | Gdf15 |
| ENSMUSG00( | 9.51645228 | 1.4635411 | 0.55409052 | 4.38E-04 | 0.00241336 | Gm37170 |
| ENSMUSG00( | 27.4508289 | 1.46339459 | 0.65880044 | 0.00147217 | 0.00693675 | Egr3 |
| ENSMUSG00( | 29696.1205 | 1.46127783 | 0.18442606 | 2.23E-16 | 1.12E-14 | Sqstm1 |
| ENSMUSG00( | 17.8815813 | 1.45411269 | 0.43658344 | 5.29E-05 | 3.71E-04 | Gm24727 |
| ENSMUSG00( | 12.4862183 | 1.4531721 | 0.48882537 | 1.64E-04 | 0.00101377 | Tnfsfm13 |
| ENSMUSG00( | 39.7769034 | 1.45119598 | 0.25456707 | 8.80E-10 | 1.67E-08 | Gm12968 |
| ENSMUSG00( | 8.49769385 | 1.45055325 | 0.63047463 | 0.00106813 | 0.00524983 | Gm29686 |
| ENSMUSG00( | 9.68396479 | 1.44971947 | 0.79667973 | 0.00327164 | 0.01381936 | Aldh5a1 |
| ENSMUSG00( | 12.760557 | 1.44691545 | 0.7408892 | 0.00201795 | 0.00910659 | Gldc |
| ENSMUSG00( | 41.7329307 | 1.43632021 | 0.29327797 | 7.49E-08 | 9.71E-07 | Leng9 |
| ENSMUSG00( | 26.3849474 | 1.43150258 | 0.36070312 | 5.02E-06 | 4.45E-05 | 1500002C15I |
| ENSMUSG00( | 16.4470544 | 1.4287442 | 0.43607003 | 6.41E-05 | 4.41E-04 | Pira1 |
| ENSMUSG00( | 7.5649521 | 1.42475278 | 0.84385643 | 0.00355224 | 0.01480689 | Gm45779 |
| ENSMUSG00( | 51.0480042 | 1.42461849 | 0.44962409 | 8.53E-05 | 5.68E-04 | Gm46189 |

| | | | | | | |
|---|---|---|---|---|---|---|
| ENSMUSG00( | 11.5299052 | 1.41458933 | 0.53554275 | 5.12E-04 | 0.00276723 | |
| ENSMUSG00( | 3.77920007 | 1.41432643 | 1.60575153 | 0.01125405 | 0.03915993 | Gm45051 |
| ENSMUSG00( | 13.4527438 | 1.40826344 | 0.50262396 | 2.89E-04 | 0.00167324 | 1110025M09I |
| ENSMUSG00( | 182.818453 | 1.4074029 | 0.2334906 | 1.13E-10 | 2.49E-09 | Slc1a4 |
| ENSMUSG00( | 138.273539 | 1.39961651 | 0.2594861 | 4.80E-09 | 8.04E-08 | Casz1 |
| ENSMUSG00( | 23.5962191 | 1.39813882 | 0.36634225 | 9.06E-06 | 7.59E-05 | 0610040B10F |
| ENSMUSG00( | 325.667199 | 1.39520453 | 0.14425097 | 3.10E-23 | 4.12E-21 | Il36g |
| ENSMUSG00( | 411.77443 | 1.39467469 | 0.19963646 | 1.90E-13 | 6.44E-12 | Gprc5a |
| ENSMUSG00( | 25.529325 | 1.39160469 | 0.3711448 | 1.14E-05 | 9.33E-05 | Gm41611 |
| ENSMUSG00( | 25.7737757 | 1.38219354 | 0.31039988 | 5.74E-07 | 6.22E-06 | Mid1-ps1 |
| ENSMUSG00( | 5.5616354 | 1.37593557 | 0.80356583 | 0.00356549 | 0.01484999 | Gm44649 |
| ENSMUSG00( | 31.1856859 | 1.37398603 | 0.44362836 | 1.08E-04 | 7.05E-04 | Disp3 |
| ENSMUSG00( | 12.8591611 | 1.37120809 | 0.61113527 | 0.00114499 | 0.00557532 | |
| ENSMUSG00( | 7.77592664 | 1.36706877 | 0.7999796 | 0.00320994 | 0.01361013 | 2810029C07F |
| ENSMUSG00( | 58.9093555 | 1.36395888 | 0.83123175 | 0.00332202 | 0.01398636 | Ptpn14 |
| ENSMUSG00( | 299.271209 | 1.36312433 | 0.16560911 | 1.39E-17 | 8.42E-16 | Cldn12 |
| ENSMUSG00( | 113.573253 | 1.36077615 | 0.19949409 | 6.70E-13 | 2.07E-11 | Gm21399 |
| ENSMUSG00( | 231.861119 | 1.3596897 | 0.18618513 | 2.04E-14 | 7.88E-13 | Crebrf |
| ENSMUSG00( | 278.552832 | 1.35850594 | 0.28993764 | 1.89E-07 | 2.26E-06 | Perm1 |
| ENSMUSG00( | 46.772883 | 1.35304621 | 0.32071178 | 1.62E-06 | 1.59E-05 | Acoxl |
| ENSMUSG00( | 4.31995108 | 1.35253515 | 1.53066989 | 0.01207572 | 0.04149403 | Mir670hg |
| ENSMUSG00( | 24.2708576 | 1.35203651 | 0.43693519 | 1.13E-04 | 7.30E-04 | |
| ENSMUSG00( | 12.8736403 | 1.35002012 | 0.66257351 | 0.00218097 | 0.00971414 | Uckl1os |
| ENSMUSG00( | 19.7095779 | 1.34970689 | 0.39259091 | 3.58E-05 | 2.63E-04 | Rnf43 |
| ENSMUSG00( | 155.975958 | 1.34911537 | 0.17347746 | 5.53E-16 | 2.63E-14 | 5830432E09F |
| ENSMUSG00( | 435.06754 | 1.34298292 | 0.20458873 | 3.71E-12 | 1.03E-10 | Hvcn1 |
| ENSMUSG00( | 30.1929916 | 1.33688411 | 0.38024902 | 2.79E-05 | 2.09E-04 | Mok |
| ENSMUSG00( | 1008.11654 | 1.33397843 | 0.13182365 | 3.66E-25 | 6.40E-23 | Chd2 |
| ENSMUSG00( | 15.3574448 | 1.33366724 | 0.49191014 | 3.78E-04 | 0.00211456 | Gm42141 |
| ENSMUSG00( | 5842.22435 | 1.32669465 | 0.43443626 | 1.32E-04 | 8.37E-04 | Plk2 |
| ENSMUSG00( | 22.0091427 | 1.32645702 | 0.36889457 | 2.21E-05 | 1.70E-04 | Gm9917 |
| ENSMUSG00( | 60.7668602 | 1.32500883 | 0.27610139 | 1.13E-07 | 1.41E-06 | Card6 |
| ENSMUSG00( | 157.741145 | 1.32130591 | 0.26250233 | 3.23E-08 | 4.54E-07 | Fzd5 |
| ENSMUSG00( | 478.389848 | 1.31733989 | 0.10898108 | 9.69E-35 | 4.78E-32 | Carhsp1 |
| ENSMUSG00( | 20.8515038 | 1.31587211 | 0.56112439 | 9.18E-04 | 0.00460954 | |
| ENSMUSG00( | 6.56145371 | 1.31373611 | 1.40372323 | 0.00709103 | 0.02634195 | Gm44187 |
| ENSMUSG00( | 10.8062001 | 1.31242347 | 0.66728284 | 0.00215866 | 0.00963172 | Gemin4 |
| ENSMUSG00( | 5.27865781 | 1.31143935 | 0.94100275 | 0.00564951 | 0.02179339 | Gm31462 |
| ENSMUSG00( | 400.838084 | 1.30738933 | 0.22312976 | 3.20E-10 | 6.56E-09 | Atf5 |
| ENSMUSG00( | 27.3254238 | 1.30623343 | 0.33525744 | 6.81E-06 | 5.86E-05 | 9130230N09F |
| ENSMUSG00( | 179.145899 | 1.30560597 | 0.2313111 | 1.20E-09 | 2.20E-08 | |

| | | | | | | |
|---|---|---|---|---|---|---|
| ENSMUSG00( | 11.7791111 | 1.30300303 | 0.64371027 | 0.00194255 | 0.00881137 | Gm47246 |
| ENSMUSG00( | 5.50281075 | 1.30253237 | 0.84253873 | 0.00490161 | 0.01937188 | Gm19265 |
| ENSMUSG00( | 22.3586104 | 1.29891229 | 0.38622145 | 4.95E-05 | 3.50E-04 | Gm32401 |
| ENSMUSG00( | 40.6868215 | 1.297053 | 0.4254322 | 1.33E-04 | 8.43E-04 | Pomc |
| ENSMUSG00( | 14.5241157 | 1.29347584 | 0.43787019 | 1.99E-04 | 0.00120455 | Sall2 |
| ENSMUSG00( | 17.6448256 | 1.29168949 | 0.44642037 | 2.29E-04 | 0.0013638 | Zfpm2 |
| ENSMUSG00( | 5.23135974 | 1.29084271 | 0.89522429 | 0.00521065 | 0.02039137 | Gm28731 |
| ENSMUSG00( | 151.282608 | 1.29038081 | 0.27594629 | 2.07E-07 | 2.44E-06 | Kctd7 |
| ENSMUSG00( | 38.5213663 | 1.28566174 | 0.29609205 | 9.78E-07 | 1.01E-05 | |
| ENSMUSG00( | 109.932722 | 1.2840678 | 0.2620732 | 6.78E-08 | 8.90E-07 | Zfp469 |
| ENSMUSG00( | 11.650041 | 1.28235393 | 0.7456769 | 0.00364459 | 0.01509425 | 9330159M07I |
| ENSMUSG00( | 6.95991484 | 1.28103529 | 1.00580717 | 0.00596058 | 0.02280967 | Gm19967 |
| ENSMUSG00( | 426.304844 | 1.27231596 | 0.10774352 | 3.08E-33 | 1.23E-30 | St7l |
| ENSMUSG00( | 29.3416069 | 1.27139901 | 0.41120387 | 1.25E-04 | 7.99E-04 | Rnft2 |
| ENSMUSG00( | 163.23289 | 1.26889928 | 0.13361144 | 1.92E-22 | 2.41E-20 | Mxd1 |
| ENSMUSG00( | 157.396644 | 1.2654497 | 0.22326314 | 1.09E-09 | 2.01E-08 | Card14 |
| ENSMUSG00( | 16.3565305 | 1.26292692 | 0.44797784 | 2.91E-04 | 0.00168193 | Gm26901 |
| ENSMUSG00( | 22.3128461 | 1.26005089 | 0.39293245 | 8.45E-05 | 5.63E-04 | Trim66 |
| ENSMUSG00( | 153.089487 | 1.25984352 | 0.23554409 | 6.48E-09 | 1.06E-07 | Rragd |
| ENSMUSG00( | 82.1870733 | 1.25912204 | 0.25699033 | 7.24E-08 | 9.44E-07 | Hrh1 |
| ENSMUSG00( | 1020.99788 | 1.25821961 | 0.18441928 | 6.87E-13 | 2.11E-11 | Mfsd6 |
| ENSMUSG00( | 33.0582246 | 1.25526292 | 0.31299722 | 5.08E-06 | 4.51E-05 | Rdh12 |
| ENSMUSG00( | 67.7524845 | 1.25440409 | 0.4830197 | 4.95E-04 | 0.0026909 | Acss2 |
| ENSMUSG00( | 65.6603149 | 1.24923619 | 0.355275 | 2.84E-05 | 2.13E-04 | Arnt2 |
| ENSMUSG00( | 2.92830387 | 1.24880483 | 1.47127024 | 0.00921858 | 0.03311735 | Gm12403 |
| ENSMUSG00( | 23.9551183 | 1.2484324 | 0.41294129 | 1.55E-04 | 9.68E-04 | Phkg1 |
| ENSMUSG00( | 4.60831054 | 1.24764596 | 1.28263265 | 0.00799696 | 0.02927911 | Gm37069 |
| ENSMUSG00( | 12.7743725 | 1.24647966 | 0.55699765 | 0.00134955 | 0.00644045 | Samd15 |
| ENSMUSG00( | 10.2600165 | 1.24513235 | 0.81134014 | 0.0054011 | 0.02097025 | Gm20072 |
| ENSMUSG00( | 38.3707982 | 1.24341293 | 0.55830856 | 0.00124858 | 0.00600127 | Wwc1 |
| ENSMUSG00( | 148.609074 | 1.24132065 | 0.17287284 | 5.65E-14 | 2.07E-12 | Zfp811 |
| ENSMUSG00( | 10.866533 | 1.24070547 | 0.53274945 | 0.00106947 | 0.00525334 | Gm44116 |
| ENSMUSG00( | 16.2823333 | 1.24003188 | 0.47930044 | 5.42E-04 | 0.00290967 | Slc26a1 |
| ENSMUSG00( | 1326.00604 | 1.23956801 | 0.15995905 | 1.10E-15 | 5.13E-14 | Bhlhe40 |
| ENSMUSG00( | 325.599214 | 1.2381561 | 0.21358074 | 4.79E-10 | 9.52E-09 | B430306N03I |
| ENSMUSG00( | 3.94004703 | 1.23797713 | 1.21748465 | 0.00842157 | 0.03065965 | Crtam |
| ENSMUSG00( | 9.34535864 | 1.23683972 | 0.6888216 | 0.00321595 | 0.01362878 | Gm19721 |
| ENSMUSG00( | 13.6026581 | 1.23587576 | 0.48901543 | 6.64E-04 | 0.00347781 | Gm37139 |
| ENSMUSG00( | 3.58519856 | 1.23491247 | 1.71840387 | 0.01091649 | 0.03815197 | Gm6288 |
| ENSMUSG00( | 108.541821 | 1.23363079 | 0.18120394 | 8.03E-13 | 2.44E-11 | Mtmr7 |
| ENSMUSG00( | 7.7457879 | 1.2293086 | 0.73517025 | 0.00376331 | 0.01550931 | Gm45548 |

| | | | | | | |
|---|---|---|---|---|---|---|
| ENSMUSG00( | 96.1687049 | 1.22389318 | 0.1969894 | 4.16E-11 | 9.98E-10 | 1500015A07F |
| ENSMUSG00( | 10.6558269 | 1.22374471 | 0.85512029 | 0.00518043 | 0.02030627 | Gm49785 |
| ENSMUSG00( | 237.49999 | 1.22238186 | 0.31132384 | 6.14E-06 | 5.36E-05 | Lat |
| ENSMUSG00( | 17.6476084 | 1.21964828 | 0.52081127 | 0.00107848 | 0.0052883 | Kif21a |
| ENSMUSG00( | 52.0849957 | 1.21825851 | 0.3560964 | 4.57E-05 | 3.26E-04 | Smagp |
| ENSMUSG00( | 278.028875 | 1.21520517 | 0.15194509 | 1.29E-16 | 6.67E-15 | Rnf167 |
| ENSMUSG00( | 9.87582636 | 1.21453142 | 0.62294244 | 0.00243486 | 0.0106832 | 4930595D18F |
| ENSMUSG00( | 86.8020425 | 1.21444235 | 0.20127873 | 1.30E-10 | 2.84E-09 | Wwtr1 |
| ENSMUSG00( | 9.77162861 | 1.2138647 | 0.55701013 | 0.00155575 | 0.00727737 | Hunk |
| ENSMUSG00( | 10.6962591 | 1.21147971 | 0.55340628 | 0.00147951 | 0.00696741 | Gfra1 |
| ENSMUSG00( | 31.1416599 | 1.21078204 | 0.39514104 | 1.42E-04 | 8.95E-04 | Gm56053 |
| ENSMUSG00( | 12.2569174 | 1.2102702 | 0.59387366 | 0.00209338 | 0.00938886 | Gm15911 |
| ENSMUSG00( | 1064.6904 | 1.20987821 | 0.21836229 | 2.34E-09 | 4.10E-08 | Errfi1 |
| ENSMUSG00( | 8.5209313 | 1.20950666 | 1.45049103 | 0.00928016 | 0.03327955 | Gm13091 |
| ENSMUSG00( | 150.171981 | 1.20744261 | 0.15039056 | 7.93E-17 | 4.25E-15 | Sgtb |
| ENSMUSG00( | 199.327924 | 1.2069722 | 0.14108201 | 1.26E-18 | 8.72E-17 | Trmt12 |
| ENSMUSG00( | 59.8484377 | 1.20519108 | 0.36824339 | 6.83E-05 | 4.66E-04 | Brd3os |
| ENSMUSG00( | 4.99905983 | 1.20477845 | 0.8949054 | 0.00660275 | 0.02484182 | Platr1 |
| ENSMUSG00( | 331.487678 | 1.20202716 | 0.25801353 | 2.24E-07 | 2.63E-06 | Atp13a4 |
| ENSMUSG00( | 21.3979276 | 1.19929729 | 0.43480187 | 3.68E-04 | 0.00206438 | Ggn |
| ENSMUSG00( | 77.4850353 | 1.19769852 | 0.27809932 | 1.23E-06 | 1.24E-05 | Pde8b |
| ENSMUSG00( | 5.71169634 | 1.19580365 | 1.6692554 | 0.00833088 | 0.03034923 | Gm48678 |
| ENSMUSG00( | 133.894746 | 1.1945857 | 0.1744359 | 6.40E-13 | 1.98E-11 | Rfxank |
| ENSMUSG00( | 176.487542 | 1.18804913 | 0.21090988 | 1.42E-09 | 2.57E-08 | Celsr1 |
| ENSMUSG00( | 56.0264312 | 1.18791398 | 0.30520455 | 7.29E-06 | 6.24E-05 | 2610008E11F |
| ENSMUSG00( | 132.443033 | 1.18767605 | 0.17213603 | 4.70E-13 | 1.49E-11 | |
| ENSMUSG00( | 49.9752902 | 1.18433582 | 0.40271734 | 2.02E-04 | 0.00121831 | Src |
| ENSMUSG00( | 44.1293965 | 1.18316914 | 0.35309912 | 5.27E-05 | 3.70E-04 | Gm26520 |
| ENSMUSG00( | 10.2252766 | 1.18086983 | 0.73652037 | 0.00461351 | 0.01845515 | Mmp23 |
| ENSMUSG00( | 2.78851696 | 1.18052121 | 8.29667305 | 0.00990049 | 0.03512098 | Tbr1 |
| ENSMUSG00( | 36.6410405 | 1.17588696 | 0.3065627 | 9.36E-06 | 7.80E-05 | St14 |
| ENSMUSG00( | 742.491271 | 1.1752927 | 0.38803641 | 1.47E-04 | 9.25E-04 | Rgs1 |
| ENSMUSG00( | 14.699104 | 1.17341717 | 0.50725651 | 0.00116521 | 0.0056475 | Tomt |
| ENSMUSG00( | 18.3715359 | 1.17281493 | 0.53976441 | 0.00160948 | 0.00748155 | Tubb4a |
| ENSMUSG00( | 193.287377 | 1.17208187 | 0.13742082 | 1.28E-18 | 8.82E-17 | Tmem156 |
| ENSMUSG00( | 2242.11606 | 1.17078316 | 0.17954646 | 5.77E-12 | 1.56E-10 | Blvrb |
| ENSMUSG00( | 14.6755084 | 1.16792467 | 0.70074032 | 0.00383479 | 0.01574585 | Rasgrp4 |
| ENSMUSG00( | 106.04009 | 1.16658967 | 0.2661067 | 8.70E-07 | 9.10E-06 | Phf1 |
| ENSMUSG00( | 593.850205 | 1.16532554 | 0.17291722 | 1.98E-12 | 5.74E-11 | Gadd45b |
| ENSMUSG00( | 38.9618688 | 1.1620909 | 0.38743102 | 1.76E-04 | 0.00108149 | 4921531C22F |
| ENSMUSG00( | 25.3048672 | 1.16147307 | 0.57378114 | 0.00205051 | 0.00922865 | Serinc2 |

| | | | | | | |
|---|---|---|---|---|---|---|
| ENSMUSG00( | 18.9455543 | 1.16122322 | 0.59661417 | 0.00263364 | 0.01147724 | Aph1b |
| ENSMUSG00( | 14.5729224 | 1.16028505 | 0.48592956 | 9.86E-04 | 0.00489079 | Gm26981 |
| ENSMUSG00( | 507.286462 | 1.15987191 | 0.18024875 | 9.38E-12 | 2.46E-10 | Rnf144b |
| ENSMUSG00( | 13.482004 | 1.15890388 | 0.4911498 | 0.00107168 | 0.0052611 | C130013H08I |
| ENSMUSG00( | 309.272868 | 1.15541231 | 0.16276217 | 1.03E-13 | 3.64E-12 | Dusp18 |
| ENSMUSG00( | 5.30467581 | 1.15267524 | 0.94380146 | 0.00702323 | 0.02613638 | Gm28874 |
| ENSMUSG00( | 3104.52151 | 1.15233797 | 0.2288474 | 3.90E-08 | 5.37E-07 | Cpeb4 |
| ENSMUSG00( | 516.916405 | 1.15003902 | 0.21502468 | 7.12E-09 | 1.15E-07 | Ypel5 |
| ENSMUSG00( | 5.03645683 | 1.1474466 | 1.0602242 | 0.00877302 | 0.03171806 | Gm45220 |
| ENSMUSG00( | 10.1929201 | 1.14628311 | 0.65705944 | 0.00366956 | 0.01517892 | Gm4793 |
| ENSMUSG00( | 58.7584293 | 1.13674332 | 0.31135865 | 1.89E-05 | 1.48E-04 | Arc |
| ENSMUSG00( | 620.484767 | 1.13651972 | 0.23070073 | 6.07E-08 | 8.03E-07 | Cpeb2 |
| ENSMUSG00( | 3423.32236 | 1.1335576 | 0.11557312 | 9.16E-24 | 1.32E-21 | Slc7a11 |
| ENSMUSG00( | 58.7312245 | 1.13105559 | 0.25590437 | 7.80E-07 | 8.23E-06 | Dclk2 |
| ENSMUSG00( | 519.297775 | 1.13012778 | 0.21059312 | 6.63E-09 | 1.08E-07 | Trib3 |
| ENSMUSG00( | 91.0694701 | 1.1299469 | 0.23553943 | 1.28E-07 | 1.58E-06 | Mrgpre |
| ENSMUSG00( | 448.353855 | 1.12912682 | 0.10544552 | 7.95E-28 | 1.63E-25 | Bcl2l2 |
| ENSMUSG00( | 27.6143719 | 1.12462546 | 0.36357176 | 1.37E-04 | 8.68E-04 | 4833445I07Ri |
| ENSMUSG00( | 31.087753 | 1.12419744 | 0.33839896 | 6.35E-05 | 4.37E-04 | Gm28809 |
| ENSMUSG00( | 67.7959927 | 1.12089687 | 0.33936321 | 6.67E-05 | 4.57E-04 | 4930589L23R |
| ENSMUSG00( | 18.8840142 | 1.11979151 | 0.40561954 | 3.93E-04 | 0.00218923 | Gm16740 |
| ENSMUSG00( | 361.172602 | 1.11707458 | 0.22350694 | 4.72E-08 | 6.40E-07 | Sertad1 |
| ENSMUSG00( | 2129.23764 | 1.11212422 | 0.16003046 | 4.49E-13 | 1.44E-11 | Cd83 |
| ENSMUSG00( | 90.0128367 | 1.11048652 | 0.24780566 | 6.12E-07 | 6.58E-06 | Ahr |
| ENSMUSG00( | 86.9928064 | 1.11002293 | 0.17852095 | 4.40E-11 | 1.06E-09 | Tnfsf12 |
| ENSMUSG00( | 810.603435 | 1.10914148 | 0.16481267 | 1.51E-12 | 4.43E-11 | Zswim4 |
| ENSMUSG00( | 55.2773428 | 1.10813272 | 0.25703479 | 1.31E-06 | 1.31E-05 | Gm7160 |
| ENSMUSG00( | 14.9552072 | 1.10721309 | 0.56847893 | 0.00272473 | 0.01181493 | Gm10509 |
| ENSMUSG00( | 130.500207 | 1.10450874 | 0.23190043 | 1.75E-07 | 2.11E-06 | Clec10a |
| ENSMUSG00( | 149.325805 | 1.10450824 | 0.28527812 | 8.16E-06 | 6.92E-05 | Epha2 |
| ENSMUSG00( | 42.7099392 | 1.10415328 | 0.24424295 | 5.07E-07 | 5.56E-06 | 4833427F10R |
| ENSMUSG00( | 51.2919868 | 1.10324611 | 0.25363664 | 1.16E-06 | 1.18E-05 | Gm15726 |
| ENSMUSG00( | 8.28862812 | 1.10303014 | 0.71686209 | 0.0054641 | 0.02118541 | F420014N23R |
| ENSMUSG00( | 102.044753 | 1.10016227 | 0.30639192 | 2.43E-05 | 1.85E-04 | Gm28875 |
| ENSMUSG00( | 39.5581889 | 1.09988129 | 0.39678388 | 3.63E-04 | 0.00204146 | Lif |
| ENSMUSG00( | 11.4733792 | 1.09867058 | 0.46455553 | 0.00112966 | 0.00550867 | |
| ENSMUSG00( | 372.04963 | 1.09744997 | 0.26255081 | 2.30E-06 | 2.20E-05 | |
| ENSMUSG00( | 10.0218019 | 1.0974185 | 0.58957651 | 0.00328495 | 0.01387211 | 1700030J22R |
| ENSMUSG00( | 76.3970819 | 1.09543853 | 0.23213703 | 2.10E-07 | 2.48E-06 | Kcnb1 |
| ENSMUSG00( | 21.4805229 | 1.09484619 | 0.44328553 | 8.33E-04 | 0.00424326 | 4933440N22R |
| ENSMUSG00( | 95.7248669 | 1.09293885 | 0.18335927 | 2.27E-10 | 4.77E-09 | B3galt4 |

| | | | | | | |
|---|---|---|---|---|---|---|
| ENSMUSG00( | 246.557283 | 1.0913926 | 0.18001943 | 1.26E-10 | 2.75E-09 | Mettl7a1 |
| ENSMUSG00( | 1723.3214 | 1.0881862 | 0.15745586 | 4.67E-13 | 1.48E-11 | Rusc2 |
| ENSMUSG00( | 60.7731256 | 1.08560105 | 0.26436993 | 3.20E-06 | 2.98E-05 | 4632427E13F |
| ENSMUSG00( | 12.8054984 | 1.08525495 | 0.62521757 | 0.0038823 | 0.01590198 | Gm5112 |
| ENSMUSG00( | 3.82260938 | 1.07913018 | 2.47006164 | 0.01083853 | 0.0379111 | |
| ENSMUSG00( | 19.6970018 | 1.07868258 | 0.6367981 | 0.00424287 | 0.01712779 | Tinagl1 |
| ENSMUSG00( | 995.074294 | 1.07776078 | 0.15450781 | 4.14E-13 | 1.34E-11 | Tmsb10 |
| ENSMUSG00( | 428.276326 | 1.07425398 | 0.09315975 | 7.61E-32 | 2.60E-29 | Cdc42ep4 |
| ENSMUSG00( | 45.9492038 | 1.07145229 | 0.24504941 | 1.05E-06 | 1.08E-05 | 1600020E01F |
| ENSMUSG00( | 383.708677 | 1.07086864 | 0.20163783 | 9.71E-09 | 1.54E-07 | Slc39a4 |
| ENSMUSG00( | 438.018513 | 1.07018771 | 0.24414187 | 8.80E-07 | 9.19E-06 | Sesn2 |
| ENSMUSG00( | 123.058741 | 1.06939367 | 0.21450957 | 5.86E-08 | 7.79E-07 | Fam131a |
| ENSMUSG00( | 53.3927459 | 1.06895443 | 0.33624899 | 1.07E-04 | 6.94E-04 | Zfp189 |
| ENSMUSG00( | 362.061042 | 1.0676792 | 0.13554857 | 3.29E-16 | 1.61E-14 | Fam83h |
| ENSMUSG00( | 20.4912469 | 1.06743121 | 0.4251654 | 8.12E-04 | 0.00415039 | Usp27x |
| ENSMUSG00( | 14.5052142 | 1.067133 | 0.52422783 | 0.00235018 | 0.01034142 | Gm37127 |
| ENSMUSG00( | 6.60623832 | 1.06694242 | 0.77453123 | 0.00682937 | 0.02556821 | Espn |
| ENSMUSG00( | 21.2179885 | 1.06554806 | 0.45274028 | 0.00121263 | 0.00585027 | Clcn2 |
| ENSMUSG00( | 303.996531 | 1.06536729 | 0.18380603 | 5.71E-10 | 1.12E-08 | Trp53inp2 |
| ENSMUSG00( | 184.871366 | 1.06462164 | 0.21199041 | 4.39E-08 | 6.00E-07 | Nupr1 |
| ENSMUSG00( | 181.580422 | 1.06410727 | 0.13943859 | 2.26E-15 | 9.98E-14 | Prkab2 |
| ENSMUSG00( | 9.80614784 | 1.06255032 | 0.64710383 | 0.00471595 | 0.01875201 | |
| ENSMUSG00( | 10.9733682 | 1.06189161 | 0.80085752 | 0.00667483 | 0.02506237 | Gm28424 |
| ENSMUSG00( | 892.124617 | 1.06158208 | 0.16718912 | 1.79E-11 | 4.56E-10 | Jarid2 |
| ENSMUSG00( | 158.818681 | 1.05897172 | 0.17707817 | 2.02E-10 | 4.29E-09 | Serac1 |
| ENSMUSG00( | 24.0965181 | 1.05840534 | 0.41058222 | 6.70E-04 | 0.00350841 | Rab2b |
| ENSMUSG00( | 399.054092 | 1.05670162 | 0.5064036 | 0.00193853 | 0.0088003 | Osgin1 |
| ENSMUSG00( | 12.1059741 | 1.05585457 | 0.73963489 | 0.00613615 | 0.02338544 | Sptbn4 |
| ENSMUSG00( | 19.9128586 | 1.05437717 | 0.47214007 | 0.00154639 | 0.00724169 | 5031434O11F |
| ENSMUSG00( | 197.308721 | 1.05414343 | 0.14620376 | 5.17E-14 | 1.90E-12 | Zfp7 |
| ENSMUSG00( | 2139.46268 | 1.05404385 | 0.1075839 | 7.43E-24 | 1.11E-21 | Slc39a10 |
| ENSMUSG00( | 95.1030816 | 1.05337564 | 0.2311873 | 4.48E-07 | 4.98E-06 | Atosa |
| ENSMUSG00( | 13.5379258 | 1.05140699 | 0.46984246 | 0.00157325 | 0.00734491 | |
| ENSMUSG00( | 793.429875 | 1.05081333 | 0.15841484 | 3.05E-12 | 8.51E-11 | Hmga2 |
| ENSMUSG00( | 222.089137 | 1.05004955 | 0.33257508 | 1.16E-04 | 7.48E-04 | Gm14221 |
| ENSMUSG00( | 128.475089 | 1.04987182 | 0.27464699 | 9.14E-06 | 7.64E-05 | Map1b |
| ENSMUSG00( | 11.8660719 | 1.0497694 | 0.66398532 | 0.0051966 | 0.02034586 | A830005F24F |
| ENSMUSG00( | 12.1394757 | 1.04848327 | 0.65252734 | 0.00511328 | 0.02008594 | Gm15787 |
| ENSMUSG00( | 26.1483726 | 1.04799073 | 0.42915663 | 0.00094265 | 0.00471427 | Galnt6os |
| ENSMUSG00( | 2286.22171 | 1.042159 | 0.17972456 | 6.72E-10 | 1.30E-08 | Prr13 |
| ENSMUSG00( | 59.4193617 | 1.03904565 | 0.18813221 | 3.19E-09 | 5.49E-08 | G530011O06F |

| | | | | | | |
|---|---|---|---|---|---|---|
| ENSMUSG00( | 5.62957629 | 1.03862821 | 0.96162696 | 0.0098549 | 0.03498887 | Gm17344 |
| ENSMUSG00( | 10.4635579 | 1.03800709 | 0.66993428 | 0.00559922 | 0.02162427 | Rdh5 |
| ENSMUSG00( | 38.7788373 | 1.03795322 | 0.29160903 | 3.02E-05 | 2.25E-04 | |
| ENSMUSG00( | 1048.24755 | 1.03779297 | 0.17908674 | 5.92E-10 | 1.16E-08 | Vegfa |
| ENSMUSG00( | 7.69106056 | 1.03722723 | 0.80474071 | 0.00757729 | 0.02794395 | Cers4 |
| ENSMUSG00( | 14.4824717 | 1.03434726 | 0.68127888 | 0.00547685 | 0.02122708 | 6430562O15F |
| ENSMUSG00( | 8.12508925 | 1.03089249 | 0.72688801 | 0.00660709 | 0.02485257 | Gm45820 |
| ENSMUSG00( | 7.9893763 | 1.02752882 | 1.02750486 | 0.0089628 | 0.03232122 | Gm17034 |
| ENSMUSG00( | 53.7307746 | 1.02746892 | 0.36375886 | 3.39E-04 | 0.00192422 | Apold1 |
| ENSMUSG00( | 716.591472 | 1.02697167 | 0.11602121 | 7.42E-20 | 6.19E-18 | Acvrl1 |
| ENSMUSG00( | 210.333417 | 1.02519474 | 0.4616892 | 0.00153034 | 0.00718055 | Dusp14 |
| ENSMUSG00( | 503.559813 | 1.02516596 | 0.10479121 | 1.36E-23 | 1.89E-21 | Socs5 |
| ENSMUSG00( | 861.172773 | 1.02465014 | 0.11990631 | 1.25E-18 | 8.71E-17 | Ampd3 |
| ENSMUSG00( | 455.629495 | 1.02452296 | 0.15670382 | 5.69E-12 | 1.54E-10 | Slc66a1 |
| ENSMUSG00( | 378.998706 | 1.02054315 | 0.16317178 | 3.67E-11 | 8.86E-10 | Hivep2 |
| ENSMUSG00( | 157.890852 | 1.02024296 | 0.18517385 | 3.20E-09 | 5.49E-08 | Prkd2 |
| ENSMUSG00( | 109.749655 | 1.01942487 | 0.17594421 | 6.45E-10 | 1.25E-08 | Ly96 |
| ENSMUSG00( | 123.429237 | 1.01757874 | 0.21117974 | 1.32E-07 | 1.62E-06 | Soat2 |
| ENSMUSG00( | 68.2211939 | 1.01681289 | 0.36561067 | 3.88E-04 | 0.00216344 | Adgrg1 |
| ENSMUSG00( | 6391.1105 | 1.01657257 | 0.08556743 | 7.94E-34 | 3.41E-31 | Atf4 |
| ENSMUSG00( | 1290.75528 | 1.01294775 | 0.23138214 | 1.18E-06 | 1.19E-05 | Cd28 |
| ENSMUSG00( | 468.194249 | 1.01197236 | 0.20700682 | 9.39E-08 | 1.19E-06 | Mocos |
| ENSMUSG00( | 43.71121 | 1.00836793 | 0.33710069 | 2.32E-04 | 0.00137661 | Cmtm8 |
| ENSMUSG00( | 14.892603 | 1.00665864 | 0.52299645 | 0.00314632 | 0.01339122 | Gm42611 |
| ENSMUSG00( | 2625.35214 | 1.00601341 | 0.08418828 | 3.26E-34 | 1.56E-31 | Nckap5l |
| ENSMUSG00( | 36.8253433 | 1.00491807 | 0.40447399 | 8.76E-04 | 0.00443162 | H2-DMa |
| ENSMUSG00( | 10.6029497 | 1.0038862 | 0.60683363 | 0.00506831 | 0.01993671 | 1700084E18F |
| ENSMUSG00( | 15.3804954 | 1.0037586 | 0.46515954 | 0.00199283 | 0.00901263 | Trim63 |
| ENSMUSG00( | 36.953323 | 1.00336825 | 0.29992039 | 6.77E-05 | 4.63E-04 | |
| ENSMUSG00( | 18.793003 | 1.00245309 | 0.54532209 | 0.00353596 | 0.01474637 | Gm45477 |
| ENSMUSG00( | 18.8950873 | 1.00218615 | 0.4160955 | 0.00112819 | 0.00550466 | Gm45854 |
| ENSMUSG00( | 323.457879 | 1.00194507 | 0.12501998 | 1.07E-16 | 5.63E-15 | Zfp46 |
| ENSMUSG00( | 13.5167573 | 1.00021236 | 0.50982583 | 0.00296741 | 0.01270718 | Tnfsf13 |
| ENSMUSG00( | 2257.92314 | -1.0024814 | 0.1248159 | 9.17E-17 | 4.85E-15 | Rif1 |
| ENSMUSG00( | 4000.89155 | -1.0028083 | 0.09200778 | 8.14E-29 | 1.84E-26 | Wdr43 |
| ENSMUSG00( | 134.657111 | -1.0031745 | 0.27837464 | 2.65E-05 | 2.00E-04 | Eif4e3 |
| ENSMUSG00( | 85.1093331 | -1.0064356 | 0.33564242 | 2.09E-04 | 0.00125725 | Gm35315 |
| ENSMUSG00( | 259.014513 | -1.0068201 | 0.15796959 | 1.81E-11 | 4.62E-10 | Ngly1 |
| ENSMUSG00( | 2640.56837 | -1.0069614 | 0.05647925 | 4.88E-72 | 1.36E-68 | Gtf2i |
| ENSMUSG00( | 454.426713 | -1.0086575 | 0.15254119 | 3.49E-12 | 9.66E-11 | Alms1 |
| ENSMUSG00( | 1070.30909 | -1.0109137 | 0.10313864 | 1.19E-23 | 1.69E-21 | Irag2 |

| | | | | | | |
|---|---|---|---|---|---|---|
| ENSMUSG00( | 443.134207 | -1.013336 | 0.23174627 | 1.07E-06 | 1.10E-05 | St6gal1 |
| ENSMUSG00( | 778.459127 | -1.0164962 | 0.09052499 | 4.24E-30 | 1.17E-27 | Lrrc75a |
| ENSMUSG00( | 652.878758 | -1.017968 | 0.18131143 | 1.77E-09 | 3.16E-08 | Donson |
| ENSMUSG00( | 1283.83435 | -1.0197845 | 0.13217256 | 1.06E-15 | 4.92E-14 | Brca1 |
| ENSMUSG00( | 141.878677 | -1.0212617 | 0.29413042 | 3.97E-05 | 2.88E-04 | Csf3r |
| ENSMUSG00( | 1244.86943 | -1.0262466 | 0.11256525 | 6.36E-21 | 6.31E-19 | Fkbp5 |
| ENSMUSG00( | 2625.15924 | -1.0277104 | 0.08970552 | 1.60E-31 | 5.37E-29 | Lig1 |
| ENSMUSG00( | 239.977935 | -1.0310011 | 0.18100768 | 9.35E-10 | 1.76E-08 | Kifc3 |
| ENSMUSG00( | 118.660049 | -1.0310569 | 0.37891078 | 4.10E-04 | 0.00226717 | Cirbp |
| ENSMUSG00( | 81.1450103 | -1.0318259 | 0.26780475 | 1.07E-05 | 8.77E-05 | Nemp2 |
| ENSMUSG00( | 222.168575 | -1.0325408 | 0.15408426 | 2.02E-12 | 5.82E-11 | Pidd1 |
| ENSMUSG00( | 658.556451 | -1.0335311 | 0.124861 | 1.69E-17 | 1.01E-15 | Dync2i2 |
| ENSMUSG00( | 1520.90432 | -1.0356071 | 0.11641855 | 9.88E-20 | 8.20E-18 | Rab32 |
| ENSMUSG00( | 172.064145 | -1.0356537 | 0.14997779 | 4.73E-13 | 1.50E-11 | Zfp862-ps |
| ENSMUSG00( | 101.598172 | -1.0365769 | 0.276034 | 1.49E-05 | 1.19E-04 | Tmem241 |
| ENSMUSG00( | 534.214914 | -1.0372419 | 0.1246309 | 8.36E-18 | 5.21E-16 | Nfil3 |
| ENSMUSG00( | 135.938632 | -1.0372759 | 0.29910186 | 4.32E-05 | 3.10E-04 | Bmyc |
| ENSMUSG00( | 533.497854 | -1.0378797 | 0.1242517 | 5.48E-18 | 3.56E-16 | Lacc1 |
| ENSMUSG00( | 3463.52134 | -1.0389454 | 0.12120154 | 8.35E-19 | 5.93E-17 | Vps13c |
| ENSMUSG00( | 819.64745 | -1.0426413 | 0.08506596 | 1.68E-35 | 9.40E-33 | Gpn1 |
| ENSMUSG00( | 2233.48384 | -1.0449452 | 0.05377879 | 3.82E-85 | 1.60E-81 | Xbp1 |
| ENSMUSG00( | 161.63765 | -1.0467441 | 0.18159714 | 8.32E-10 | 1.58E-08 | Kif24 |
| ENSMUSG00( | 79.9568789 | -1.0504083 | 0.31550124 | 9.01E-05 | 5.96E-04 | Dlg3 |
| ENSMUSG00( | 560.405564 | -1.0508284 | 0.20825666 | 3.76E-08 | 5.20E-07 | Mcm8 |
| ENSMUSG00( | 3463.34007 | -1.0550492 | 0.41037226 | 6.22E-04 | 0.00328431 | Itga4 |
| ENSMUSG00( | 46.0810002 | -1.0574528 | 0.44640217 | 0.0012192 | 0.00587858 | Il15ra |
| ENSMUSG00( | 544.680785 | -1.0609489 | 0.11717487 | 1.86E-20 | 1.65E-18 | Parp2 |
| ENSMUSG00( | 462.723792 | -1.0623505 | 0.14660605 | 4.71E-14 | 1.74E-12 | Pomt1 |
| ENSMUSG00( | 1605.68231 | -1.0630572 | 0.07712509 | 2.82E-44 | 3.37E-41 | Dnaja3 |
| ENSMUSG00( | 94.4343267 | -1.0645221 | 0.27585905 | 8.80E-06 | 7.39E-05 | F730043M19I |
| ENSMUSG00( | 9.80103169 | -1.0648865 | 0.70092483 | 0.00663397 | 0.02493688 | Gm9027 |
| ENSMUSG00( | 240.561183 | -1.0652037 | 0.25530577 | 2.45E-06 | 2.33E-05 | Pdpr |
| ENSMUSG00( | 405.198642 | -1.0673244 | 0.25331177 | 2.35E-06 | 2.24E-05 | S1pr1 |
| ENSMUSG00( | 19.2755933 | -1.0682637 | 0.57016981 | 0.00364905 | 0.01510898 | Gm42600 |
| ENSMUSG00( | 376.161148 | -1.0689875 | 0.14098073 | 2.66E-15 | 1.16E-13 | Nagk |
| ENSMUSG00( | 25.2275085 | -1.0692477 | 0.90818732 | 0.0103158 | 0.03638635 | Ptgfrn |
| ENSMUSG00( | 181.613064 | -1.0704356 | 0.35437163 | 1.79E-04 | 0.0010949 | Cysltr1 |
| ENSMUSG00( | 10318.1075 | -1.0764744 | 0.11340096 | 2.13E-22 | 2.60E-20 | Mki67 |
| ENSMUSG00( | 462.601075 | -1.0788599 | 0.14409172 | 6.39E-15 | 2.65E-13 | Gpsm1 |
| ENSMUSG00( | 47.3760552 | -1.0796737 | 0.3448186 | 1.41E-04 | 8.90E-04 | Svil |
| ENSMUSG00( | 223.961155 | -1.0798781 | 0.20209465 | 8.83E-09 | 1.41E-07 | Neil3 |

| | | | | | | |
|---|---|---|---|---|---|---|
| ENSMUSG00( | 732.704738 | -1.0805907 | 0.08889186 | 4.84E-35 | 2.54E-32 | Prkdc |
| ENSMUSG00( | 11.9984382 | -1.0807809 | 1.05410053 | 0.01040431 | 0.03665998 | Gm48239 |
| ENSMUSG00( | 77.082734 | -1.0824806 | 0.30570994 | 3.36E-05 | 2.47E-04 | Lrfn1 |
| ENSMUSG00( | 1352.11552 | -1.0840235 | 0.13454569 | 7.16E-17 | 3.87E-15 | Jag2 |
| ENSMUSG00( | 622.239074 | -1.0894108 | 0.11072301 | 8.32E-24 | 1.22E-21 | Tdp1 |
| ENSMUSG00( | 260.544491 | -1.1030593 | 0.13247031 | 7.88E-18 | 4.95E-16 | Abraxas1 |
| ENSMUSG00( | 396.731294 | -1.1032048 | 0.16410122 | 1.58E-12 | 4.62E-11 | Adprs |
| ENSMUSG00( | 100.680943 | -1.106774 | 0.44466695 | 7.82E-04 | 0.00401909 | Ccdc18 |
| ENSMUSG00( | 476.662845 | -1.1100589 | 0.21681909 | 2.41E-08 | 3.52E-07 | Rad54l |
| ENSMUSG00( | 2300.11198 | -1.1129856 | 0.10214091 | 5.98E-29 | 1.39E-26 | Slc44a2 |
| ENSMUSG00( | 121.655706 | -1.1154187 | 0.21142163 | 1.30E-08 | 2.01E-07 | Tbc1d32 |
| ENSMUSG00( | 57.5852635 | -1.1157003 | 0.38355367 | 2.60E-04 | 0.00152322 | Mapre3 |
| ENSMUSG00( | 985.730383 | -1.1157369 | 0.13355738 | 5.72E-18 | 3.70E-16 | Coro2a |
| ENSMUSG00( | 261.995206 | -1.1163818 | 0.15001439 | 1.07E-14 | 4.26E-13 | Tango6 |
| ENSMUSG00( | 1638.80685 | -1.1195195 | 0.08481671 | 4.36E-41 | 4.06E-38 | Iqgap3 |
| ENSMUSG00( | 1852.4527 | -1.1201748 | 0.11220802 | 1.96E-24 | 3.04E-22 | Pold1 |
| ENSMUSG00( | 636.17304 | -1.1231426 | 0.1924099 | 4.24E-10 | 8.51E-09 | Scamp5 |
| ENSMUSG00( | 1072.34777 | -1.1243586 | 0.10628646 | 3.92E-27 | 7.83E-25 | Timeless |
| ENSMUSG00( | 72.9114469 | -1.1249486 | 0.44377284 | 8.87E-04 | 0.00447376 | Ttc39a |
| ENSMUSG00( | 36.8593034 | -1.125451 | 0.63800061 | 0.00489014 | 0.01933112 | Ssbp2 |
| ENSMUSG00( | 227.905427 | -1.1263322 | 0.18187217 | 5.11E-11 | 1.20E-09 | Tmem260 |
| ENSMUSG00( | 567.160593 | -1.1275158 | 0.13664108 | 1.30E-17 | 7.95E-16 | Mrps35 |
| ENSMUSG00( | 360.247505 | -1.129617 | 0.16313239 | 3.99E-13 | 1.30E-11 | Tmem94 |
| ENSMUSG00( | 113.368366 | -1.1302527 | 0.25123315 | 6.08E-07 | 6.55E-06 | Rad51c |
| ENSMUSG00( | 1301.04895 | -1.137008 | 0.09685968 | 4.10E-33 | 1.60E-30 | Pole |
| ENSMUSG00( | 675.258355 | -1.1372006 | 0.15079134 | 3.66E-15 | 1.60E-13 | Mlh1 |
| ENSMUSG00( | 23.2821418 | -1.1391817 | 0.7103084 | 0.00431136 | 0.0173917 | Cd300e |
| ENSMUSG00( | 339.198886 | -1.1397576 | 0.13669734 | 7.59E-18 | 4.82E-16 | Mmachc |
| ENSMUSG00( | 4255.9678 | -1.1398095 | 0.12494161 | 5.84E-21 | 5.83E-19 | Atad2 |
| ENSMUSG00( | 417.503307 | -1.1401298 | 0.19068334 | 2.05E-10 | 4.34E-09 | Rab3il1 |
| ENSMUSG00( | 290.457906 | -1.1443182 | 0.15902391 | 5.52E-14 | 2.03E-12 | Adarb1 |
| ENSMUSG00( | 1046.86713 | -1.1460949 | 0.08705272 | 1.92E-40 | 1.61E-37 | Adpgk |
| ENSMUSG00( | 92.0268932 | -1.1470609 | 0.30151102 | 1.13E-05 | 9.23E-05 | Mpv17l |
| ENSMUSG00( | 1610.74918 | -1.1483814 | 0.11804117 | 1.34E-23 | 1.87E-21 | Sdc1 |
| ENSMUSG00( | 121.724412 | -1.1544599 | 0.24952745 | 3.00E-07 | 3.43E-06 | Inka1 |
| ENSMUSG00( | 37.2954685 | -1.154874 | 0.48657507 | 0.00108968 | 0.00533856 | Afmid |
| ENSMUSG00( | 1100.93782 | -1.1555631 | 0.10744849 | 7.68E-28 | 1.59E-25 | Cdc6 |
| ENSMUSG00( | 265.150766 | -1.1577206 | 0.216259 | 7.21E-09 | 1.17E-07 | Tln2 |
| ENSMUSG00( | 530.979458 | -1.1589824 | 0.14675944 | 2.52E-16 | 1.26E-14 | Nfix |
| ENSMUSG00( | 900.3995 | -1.1616479 | 0.19617486 | 2.46E-10 | 5.13E-09 | Ccne2 |
| ENSMUSG00( | 280.282666 | -1.163321 | 0.15430198 | 4.48E-15 | 1.92E-13 | Emilin1 |

| | | | | | | |
|---|---|---|---|---|---|---|
| ENSMUSG00( | 75.852775 | -1.1642657 | 0.24795697 | 1.91E-07 | 2.27E-06 | Ccl22 |
| ENSMUSG00( | 72.6996042 | -1.171958 | 0.26314934 | 7.30E-07 | 7.77E-06 | Ttc39aos1 |
| ENSMUSG00( | 445.15948 | -1.1727751 | 0.13732136 | 1.24E-18 | 8.69E-17 | Tle1 |
| ENSMUSG00( | 35.3174561 | -1.1753635 | 0.4813363 | 0.00106856 | 0.0052504 | Casp7 |
| ENSMUSG00( | 142.194398 | -1.1776697 | 0.21008226 | 1.38E-09 | 2.51E-08 | Slc2a3 |
| ENSMUSG00( | 39.1688713 | -1.1780151 | 0.44640877 | 5.24E-04 | 0.0028258 | Shcbp1l |
| ENSMUSG00( | 238.91797 | -1.1786317 | 0.16425962 | 6.17E-14 | 2.24E-12 | Sgsm2 |
| ENSMUSG00( | 496.774405 | -1.1803662 | 0.21181174 | 1.97E-09 | 3.50E-08 | Fbxo5 |
| ENSMUSG00( | 215.787911 | -1.1821661 | 0.16562269 | 8.07E-14 | 2.86E-12 | Gna15 |
| ENSMUSG00( | 531.251919 | -1.1937354 | 0.12221867 | 1.31E-23 | 1.84E-21 | Nol4l |
| ENSMUSG00( | 65.4903268 | -1.194059 | 0.47011178 | 5.75E-04 | 0.00306945 | Gm19026 |
| ENSMUSG00( | 736.433644 | -1.1949328 | 0.11581478 | 7.40E-26 | 1.35E-23 | Cep68 |
| ENSMUSG00( | 558.465427 | -1.1952125 | 0.14876971 | 6.69E-17 | 3.64E-15 | Brip1 |
| ENSMUSG00( | 55.2854882 | -1.1985423 | 0.32641241 | 1.53E-05 | 1.22E-04 | Slfn4 |
| ENSMUSG00( | 702.422239 | -1.202311 | 0.14265829 | 2.88E-18 | 1.93E-16 | Slfn9 |
| ENSMUSG00( | 704.863565 | -1.203573 | 0.10476351 | 1.91E-31 | 6.17E-29 | Pxk |
| ENSMUSG00( | 312.411054 | -1.2060146 | 0.1483637 | 3.93E-17 | 2.19E-15 | Ctdspl |
| ENSMUSG00( | 1399.63318 | -1.2068844 | 0.10688893 | 1.23E-30 | 3.51E-28 | Tk1 |
| ENSMUSG00( | 537.044676 | -1.2080343 | 0.28436189 | 1.61E-06 | 1.58E-05 | Akap6 |
| ENSMUSG00( | 847.602671 | -1.2172677 | 0.0896717 | 2.78E-43 | 3.11E-40 | Trappc6b |
| ENSMUSG00( | 146.192968 | -1.2174891 | 0.19937991 | 9.02E-11 | 2.02E-09 | Zic2 |
| ENSMUSG00( | 445.290483 | -1.2177883 | 0.1499064 | 4.25E-17 | 2.36E-15 | Entpd6 |
| ENSMUSG00( | 726.613786 | -1.2190385 | 0.12545081 | 3.73E-23 | 4.89E-21 | Dock1 |
| ENSMUSG00( | 304.374906 | -1.2207224 | 0.26575908 | 3.33E-07 | 3.78E-06 | Fancb |
| ENSMUSG00( | 77.5851858 | -1.226388 | 0.82418048 | 0.00397792 | 0.01623017 | Osbp2 |
| ENSMUSG00( | 53.8788303 | -1.2273984 | 0.38291128 | 9.61E-05 | 6.31E-04 | Or5v1b |
| ENSMUSG00( | 24.3150407 | -1.2285777 | 0.50916703 | 9.65E-04 | 0.00481032 | 2810414N06F |
| ENSMUSG00( | 227.791043 | -1.2300198 | 0.23625525 | 1.42E-08 | 2.17E-07 | Slitrk5 |
| ENSMUSG00( | 33.2784327 | -1.2324048 | 0.43729729 | 3.18E-04 | 0.00182173 | H1f3 |
| ENSMUSG00( | 830.327955 | -1.256301 | 0.10705921 | 7.79E-33 | 2.97E-30 | Zwilch |
| ENSMUSG00( | 4933.21472 | -1.2593254 | 0.10326243 | 1.97E-35 | 1.07E-32 | Uhrf1 |
| ENSMUSG00( | 600.89754 | -1.2603505 | 0.13245728 | 1.61E-22 | 2.04E-20 | Nmral1 |
| ENSMUSG00( | 1312.60023 | -1.2738284 | 0.08264129 | 1.66E-54 | 3.08E-51 | Jade1 |
| ENSMUSG00( | 1031.09108 | -1.2759088 | 0.13126063 | 2.25E-23 | 3.04E-21 | Kif15 |
| ENSMUSG00( | 823.02468 | -1.2790188 | 0.16087697 | 1.22E-16 | 6.37E-15 | Atr |
| ENSMUSG00( | 123.711606 | -1.2809117 | 0.22817019 | 1.73E-09 | 3.09E-08 | Srl |
| ENSMUSG00( | 1389.35009 | -1.2813743 | 0.17588013 | 2.27E-14 | 8.72E-13 | Dut |
| ENSMUSG00( | 1159.03969 | -1.2855813 | 0.24081024 | 6.55E-09 | 1.07E-07 | H1f2 |
| ENSMUSG00( | 37.8493398 | -1.2953751 | 0.47462814 | 4.43E-04 | 0.00243688 | Shpk |
| ENSMUSG00( | 674.845681 | -1.2954251 | 0.1589853 | 2.77E-17 | 1.58E-15 | Dna2 |
| ENSMUSG00( | 163.29215 | -1.2966389 | 0.18038179 | 5.84E-14 | 2.13E-12 | Mpp7 |

| | | | | | | |
|---|---|---|---|---|---|---|
| ENSMUSG00( | 720.646089 | -1.297895 | 0.13063302 | 2.07E-24 | 3.17E-22 | Cln6 |
| ENSMUSG00( | 82.2878439 | -1.2979966 | 0.32424142 | 4.43E-06 | 3.99E-05 | Evl |
| ENSMUSG00( | 23.2368105 | -1.303667 | 0.53934444 | 0.00105822 | 0.00521487 | 9430091E24F |
| ENSMUSG00( | 45.7414572 | -1.3076576 | 0.41684855 | 1.25E-04 | 7.97E-04 | Nrm |
| ENSMUSG00( | 66.832585 | -1.3171063 | 0.27682744 | 1.55E-07 | 1.88E-06 | Gm13421 |
| ENSMUSG00( | 835.289689 | -1.3213902 | 0.18768494 | 1.57E-13 | 5.41E-12 | Deptor |
| ENSMUSG00( | 472.739201 | -1.3219175 | 0.14671093 | 1.56E-20 | 1.41E-18 | Brca2 |
| ENSMUSG00( | 436.049428 | -1.3233727 | 0.1683767 | 3.13E-16 | 1.54E-14 | Ppm1e |
| ENSMUSG00( | 774.659094 | -1.3335959 | 0.12241911 | 8.46E-29 | 1.89E-26 | Dck |
| ENSMUSG00( | 373.685559 | -1.3421386 | 0.15720551 | 1.22E-18 | 8.52E-17 | Cep78 |
| ENSMUSG00( | 337.011719 | -1.3487127 | 0.15576236 | 4.46E-19 | 3.32E-17 | Tfap4 |
| ENSMUSG00( | 1482.4659 | -1.3487267 | 0.11248168 | 5.91E-34 | 2.68E-31 | Prmt7 |
| ENSMUSG00( | 759.880725 | -1.3495998 | 0.15532482 | 2.32E-19 | 1.77E-17 | Ung |
| ENSMUSG00( | 2997.86098 | -1.3500837 | 0.18665264 | 3.32E-14 | 1.25E-12 | Rrm2 |
| ENSMUSG00( | 48.2621061 | -1.3532029 | 0.63236525 | 0.00222895 | 0.00989891 | Adgb |
| ENSMUSG00( | 65.6207307 | -1.354263 | 0.38050615 | 2.44E-05 | 1.86E-04 | Gm19253 |
| ENSMUSG00( | 781.516355 | -1.3601464 | 0.18099995 | 4.54E-15 | 1.93E-13 | E2f8 |
| ENSMUSG00( | 486.324677 | -1.368689 | 0.21668935 | 2.09E-11 | 5.28E-10 | Ogfrl1 |
| ENSMUSG00( | 676.70891 | -1.3687267 | 0.19795746 | 3.22E-13 | 1.06E-11 | Tcf19 |
| ENSMUSG00( | 535.460446 | -1.3725647 | 0.21663152 | 1.86E-11 | 4.72E-10 | Slc8a1 |
| ENSMUSG00( | 352.122632 | -1.3745819 | 0.2719286 | 2.81E-08 | 4.03E-07 | Pcyox1l |
| ENSMUSG00( | 3816.26623 | -1.3775703 | 0.1136097 | 5.48E-35 | 2.79E-32 | Ptpn6 |
| ENSMUSG00( | 35.8694733 | -1.3796684 | 0.57316434 | 9.97E-04 | 0.00493893 | Pde7b |
| ENSMUSG00( | 165.775801 | -1.3824864 | 0.20744877 | 2.86E-12 | 8.07E-11 | Gatd1 |
| ENSMUSG00( | 115.700914 | -1.3835037 | 0.26060605 | 8.45E-09 | 1.35E-07 | Ticam2 |
| ENSMUSG00( | 143.175151 | -1.3893058 | 0.18415624 | 5.04E-15 | 2.14E-13 | Borcs5 |
| ENSMUSG00( | 194.283029 | -1.389801 | 0.27449468 | 2.86E-08 | 4.10E-07 | Mtfp1 |
| ENSMUSG00( | 563.006763 | -1.3923654 | 0.16931363 | 1.41E-17 | 8.56E-16 | Dennd1a |
| ENSMUSG00( | 246.232903 | -1.3927725 | 0.19505464 | 7.30E-14 | 2.61E-12 | Cadps |
| ENSMUSG00( | 20.971398 | -1.4030396 | 0.52888123 | 4.46E-04 | 0.00245333 | Gm5648 |
| ENSMUSG00( | 669.641615 | -1.4041639 | 0.1228869 | 2.61E-31 | 8.26E-29 | Haus4 |
| ENSMUSG00( | 642.285058 | -1.4055559 | 0.20745483 | 8.40E-13 | 2.54E-11 | Ccne1 |
| ENSMUSG00( | 15.3665845 | -1.4067619 | 0.79756299 | 0.00379477 | 0.01561977 | Sirpb1c |
| ENSMUSG00( | 552.857929 | -1.4092226 | 0.20603827 | 5.23E-13 | 1.64E-11 | Abca1 |
| ENSMUSG00( | 304.360874 | -1.4099293 | 0.15336252 | 2.57E-21 | 2.74E-19 | Gpr180 |
| ENSMUSG00( | 124.759562 | -1.4193068 | 0.18245864 | 4.87E-16 | 2.33E-14 | Rfesd |
| ENSMUSG00( | 307.472995 | -1.4242492 | 0.27706809 | 2.31E-08 | 3.39E-07 | Iqgap2 |
| ENSMUSG00( | 418.037803 | -1.4261811 | 0.12568094 | 6.79E-31 | 2.00E-28 | Scara3 |
| ENSMUSG00( | 785.151718 | -1.4530407 | 0.1789108 | 2.33E-17 | 1.36E-15 | Lfng |
| ENSMUSG00( | 6.87840329 | -1.4536555 | 1.0066071 | 0.00799652 | 0.02927911 | Gm15792 |
| ENSMUSG00( | 8.61438132 | -1.4677496 | 0.54543618 | 4.17E-04 | 0.00230381 | BE692007 |

| | | | | | | |
|---|---|---|---|---|---|---|
| ENSMUSG00( | 24.2272381 | -1.4703517 | 0.49421901 | 1.86E-04 | 0.00113411 | Disc1 |
| ENSMUSG00( | 6.12579353 | -1.4707739 | 0.74785126 | 0.00183544 | 0.00839138 | A4galt |
| ENSMUSG00( | 126.179276 | -1.4713386 | 0.20754897 | 1.33E-13 | 4.60E-12 | Mkx |
| ENSMUSG00( | 296.461509 | -1.4732388 | 0.16848912 | 1.46E-19 | 1.14E-17 | Sla |
| ENSMUSG00( | 234.606185 | -1.4847889 | 0.17912836 | 7.61E-18 | 4.82E-16 | Dus4l |
| ENSMUSG00( | 77.7139807 | -1.4879296 | 0.26679281 | 1.79E-09 | 3.20E-08 | Fbxo48 |
| ENSMUSG00( | 297.686373 | -1.4984213 | 0.13122411 | 1.90E-31 | 6.17E-29 | Phf19 |
| ENSMUSG00( | 2411.28829 | -1.5237732 | 0.09042026 | 9.43E-65 | 2.26E-61 | Ccnd1 |
| ENSMUSG00( | 273.835412 | -1.543871 | 0.17539889 | 1.19E-19 | 9.59E-18 | Ift140 |
| ENSMUSG00( | 382.569342 | -1.5694846 | 0.15017736 | 9.81E-27 | 1.89E-24 | Bard1 |
| ENSMUSG00( | 310.42018 | -1.5706275 | 0.27157267 | 4.63E-10 | 9.23E-09 | Ankle1 |
| ENSMUSG00( | 430.219485 | -1.5731125 | 0.13799025 | 3.06E-31 | 9.52E-29 | Frmd4b |
| ENSMUSG00( | 5.36803811 | -1.5833013 | 1.3622119 | 0.00763309 | 0.02812498 | Pdzd4 |
| ENSMUSG00( | 10.3083782 | -1.5875744 | 0.96644427 | 0.00606434 | 0.02314861 | Clca4a |
| ENSMUSG00( | 4744.499 | -1.587999 | 0.06588623 | 1.60E-129 | 2.68E-125 | Nceh1 |
| ENSMUSG00( | 1075.65381 | -1.6242006 | 0.13705217 | 1.60E-33 | 6.71E-31 | Rassf2 |
| ENSMUSG00( | 79.7856732 | -1.6265989 | 0.4966045 | 8.62E-05 | 5.72E-04 | Pgbd5 |
| ENSMUSG00( | 17.9144764 | -1.6302666 | 0.40515385 | 3.27E-06 | 3.03E-05 | Gbp5 |
| ENSMUSG00( | 1336.46898 | -1.6401258 | 0.10443273 | 6.57E-57 | 1.38E-53 | Fcgr1 |
| ENSMUSG00( | 122.591905 | -1.6540888 | 0.32487654 | 3.13E-08 | 4.44E-07 | Gda |
| ENSMUSG00( | 7.0741033 | -1.6626492 | 1.02151688 | 0.00556735 | 0.02152101 | Rtbdn |
| ENSMUSG00( | 292.763474 | -1.6665436 | 0.18465529 | 1.64E-20 | 1.47E-18 | Engase |
| ENSMUSG00( | 13.0156239 | -1.7024444 | 0.60229696 | 2.23E-04 | 0.00133019 | Slamf6 |
| ENSMUSG00( | 14.0694813 | -1.7248575 | 0.59378417 | 2.54E-04 | 0.00149028 | Gm32089 |
| ENSMUSG00( | 712.984537 | -1.731629 | 0.09408214 | 1.08E-76 | 3.63E-73 | Daglb |
| ENSMUSG00( | 213.334312 | -1.7338669 | 0.41193946 | 1.51E-06 | 1.50E-05 | Fry |
| ENSMUSG00( | 222.19915 | -1.7517034 | 0.2179121 | 6.94E-17 | 3.77E-15 | Cd38 |
| ENSMUSG00( | 20.6848772 | -1.8130817 | 0.78684732 | 0.00143196 | 0.00677586 | Gm43848 |
| ENSMUSG00( | 12.2240609 | -1.8147531 | 1.31196062 | 0.00723365 | 0.02683013 | Sgpp2 |
| ENSMUSG00( | 247.986903 | -1.8239492 | 0.27683604 | 2.89E-12 | 8.15E-11 | Gjb3 |
| ENSMUSG00( | 1137.3713 | -1.835902 | 0.1712743 | 4.29E-28 | 9.33E-26 | Gas7 |
| ENSMUSG00( | 161.02479 | -1.8537357 | 0.26030826 | 6.15E-14 | 2.24E-12 | Ntng2 |
| ENSMUSG00( | 28.4861699 | -1.8701306 | 0.43568898 | 1.40E-06 | 1.40E-05 | Dnhd1 |
| ENSMUSG00( | 12.9539175 | -1.8914304 | 0.56190997 | 3.64E-05 | 2.66E-04 | Gpr31c |
| ENSMUSG00( | 11.3090625 | -1.9370777 | 0.62782215 | 1.28E-04 | 8.15E-04 | Gm47798 |
| ENSMUSG00( | 97.4147424 | -1.9672164 | 0.26158885 | 4.54E-15 | 1.93E-13 | Dcp1b |
| ENSMUSG00( | 1288.84239 | -2.0276304 | 0.23555233 | 4.99E-19 | 3.68E-17 | Cytip |
| ENSMUSG00( | 6.34052067 | -2.0350189 | 0.84306739 | 0.00112895 | 0.00550679 | Tssk4 |
| ENSMUSG00( | 80.2627982 | -2.3453756 | 0.38868889 | 8.31E-11 | 1.88E-09 | Thbs1 |
| ENSMUSG00( | 10.6467466 | -2.3965124 | 0.84871114 | 5.07E-04 | 0.00274752 | Ovgp1 |
| ENSMUSG00( | 5.13644598 | -2.4297344 | 2.7644586 | 0.00439265 | 0.01768981 | 9330179D12I |

| | | | | | | |
|---|---|---|---|---|---|---|
| ENSMUSG00( | 61.8641511 | -2.5109388 | 0.34289026 | 2.52E-14 | 9.61E-13 | AI504432 |
| ENSMUSG00( | 22.8567571 | -2.9128708 | 0.91215327 | 2.31E-04 | 0.00137467 | Vdr |
| ENSMUSG00( | 228.310471 | -2.9672104 | 0.62767109 | 1.06E-07 | 1.33E-06 | Cx3cr1 |
| ENSMUSG00( | 7.524134 | -4.268754 | 2.34265399 | 3.37E-04 | 0.00191564 | Pdzd7 |

e_name

Rik

Rik

Rik

Rik

Rik
ik

Rik

Rik

Rik

Rik
Rik
Rik

ik

Rik

Rik

Rik

ik

Rik

Rik

Rik

Rik

Rik

Rik

Rik

Rik

Rik

Rik

Rik

Rik

Rik

Rik

Rik

ik

Rik

Rik

Rik

ik

Rik

Rik

Rik

Rik

Rik

Rik

Rik

Rik

Rik

Rik

Rik

Rik
