## Supplemental information, figure compressed, table 1-5 for "Role of copper during microglial inflammation": Supplemental Table 2-compressed.pdf

| Gene_Id | baseMean | log2FoldChar | lfcSE | pvalue | padj | external_gene |
|---|---|---|---|---|---|---|
| ENSMUSG00( | 92.8357174 | 3.31685604 | 1.22226107 | 1.69E-04 | 0.02697118 | Fosb |
| ENSMUSG00( | 6.81404826 | 2.6958084 | 0.9339405 | 1.55E-04 | 0.02651804 | Efcab6 |
| ENSMUSG00( | 21.1510933 | 2.66780996 | 0.68939608 | 4.22E-06 | 0.00186988 | Mt2 |
| ENSMUSG00( | 34.6906073 | 2.53080316 | 1.22872144 | 8.26E-04 | 0.07063389 | Rn7sk |
| ENSMUSG00( | 16.7112159 | 2.4142791 | 0.4839895 | 3.10E-08 | 3.86E-05 | Sstr5 |
| ENSMUSG00( | 7535.32109 | 2.33760071 | 0.17669512 | 2.67E-41 | 3.67E-37 | Mt1 |
| ENSMUSG00( | 7.83353928 | 1.99273185 | 0.76727005 | 5.71E-04 | 0.05745655 | |
| ENSMUSG00( | 10.6286764 | 1.8465477 | 0.67187989 | 2.44E-04 | 0.03500886 | H3c6 |
| ENSMUSG00( | 15.0428897 | 1.71631199 | 0.46700667 | 9.72E-06 | 0.003606 | Gm8818 |
| ENSMUSG00( | 49.2614612 | 1.33521571 | 0.38507302 | 1.93E-05 | 0.0066349 | Vgf |
| ENSMUSG00( | 21.1454177 | 1.28412425 | 0.44900879 | 1.61E-04 | 0.02697118 | |
| ENSMUSG00( | 12.1874666 | 1.26546542 | 0.59453419 | 9.93E-04 | 0.08112686 | Sdcbp2 |
| ENSMUSG00( | 59.4193617 | 1.10190947 | 0.23531257 | 1.41E-07 | 1.38E-04 | G530011O06I |
| ENSMUSG00( | 152.388136 | 1.01156393 | 0.19720732 | 1.28E-08 | 2.52E-05 | Tcp11l2 |
| ENSMUSG00( | 524.826934 | 0.97015662 | 0.29127696 | 3.02E-05 | 0.00901695 | AY036118 |
| ENSMUSG00( | 1050.21902 | 0.90998997 | 0.12518268 | 1.65E-14 | 1.13E-10 | Atg2a |
| ENSMUSG00( | 594.968392 | 0.89945342 | 0.21906136 | 1.56E-06 | 0.00106857 | Slc30a1 |
| ENSMUSG00( | 40.2572834 | 0.86598048 | 0.33567129 | 3.20E-04 | 0.04221112 | Mettl27 |
| ENSMUSG00( | 59.7168254 | 0.82240451 | 0.24355275 | 2.82E-05 | 0.00860525 | Dyrk1b |
| ENSMUSG00( | 51.2919868 | 0.80907457 | 0.31877432 | 3.63E-04 | 0.04502376 | Gm15726 |
| ENSMUSG00( | 79.8845852 | 0.77806167 | 0.33383662 | 5.66E-04 | 0.05745655 | Calr3 |
| ENSMUSG00( | 34.2504697 | 0.74052786 | 0.3721945 | 0.00123251 | 0.09241526 | Il12rb1 |
| ENSMUSG00( | 838.417102 | 0.73931527 | 0.1677215 | 3.95E-07 | 3.62E-04 | Abhd4 |
| ENSMUSG00( | 887.302743 | 0.70802801 | 0.12998758 | 2.12E-09 | 5.81E-06 | Dnajb1 |
| ENSMUSG00( | 410.610339 | 0.68342545 | 0.14591196 | 1.18E-07 | 1.25E-04 | Ulk1 |
| ENSMUSG00( | 251.938658 | 0.66785834 | 0.18028339 | 8.40E-06 | 0.00320068 | Cyp4f18 |
| ENSMUSG00( | 111.628225 | 0.66409624 | 0.2365896 | 1.68E-04 | 0.02697118 | Zfp874b |
| ENSMUSG00( | 231.861119 | 0.62137586 | 0.19406647 | 4.98E-05 | 0.01264936 | Crebrf |
| ENSMUSG00( | 1020.99788 | 0.61473779 | 0.18876099 | 4.17E-05 | 0.01080041 | Mfsd6 |
| ENSMUSG00( | 1914.91438 | 0.60584325 | 0.09735121 | 2.82E-11 | 9.68E-08 | Zwint |
| ENSMUSG00( | 307.788112 | 0.6045814 | 0.15942881 | 6.08E-06 | 0.00245382 | Acox3 |
| ENSMUSG00( | 179.53969 | 0.59884242 | 0.24689255 | 4.56E-04 | 0.05087763 | Unc5b |
| ENSMUSG00( | 235.563158 | 0.59127218 | 0.13678617 | 6.49E-07 | 5.24E-04 | Atg14 |
| ENSMUSG00( | 146.026271 | 0.58210309 | 0.219997 | 2.66E-04 | 0.03723395 | Taf7 |
| ENSMUSG00( | 102.11229 | 0.54496923 | 0.22544318 | 4.72E-04 | 0.05227991 | Gm5526 |
| ENSMUSG00( | 766.050354 | 0.53463209 | 0.10473191 | 1.53E-08 | 2.56E-05 | Lin54 |
| ENSMUSG00( | 246.557283 | 0.53408468 | 0.19094467 | 1.76E-04 | 0.02747301 | Mettl7a1 |
| ENSMUSG00( | 516.916405 | 0.49912092 | 0.25294526 | 0.00128753 | 0.09427104 | Ypel5 |
| ENSMUSG00( | 383.708677 | 0.49256546 | 0.23961418 | 0.00110676 | 0.08621611 | Slc39a4 |
| ENSMUSG00( | 237.69961 | 0.48450555 | 0.16041943 | 9.35E-05 | 0.0183103 | Dnajb2 |

| | | | | | | |
|---|---|---|---|---|---|---|
| ENSMUSG00( | 383.102942 | 0.46192666 | 0.15379172 | 9.47E-05 | 0.0183103 | Mtfr2 |
| ENSMUSG00( | 626.832793 | 0.45886798 | 0.20319257 | 7.20E-04 | 0.06543979 | Gss |
| ENSMUSG00( | 1162.69206 | 0.4541024 | 0.1027039 | 4.43E-07 | 3.80E-04 | Mapk9 |
| ENSMUSG00( | 176.498751 | 0.44886658 | 0.19247836 | 6.23E-04 | 0.06042375 | Mical1 |
| ENSMUSG00( | 590.732213 | 0.43573417 | 0.20653102 | 0.00110342 | 0.08621611 | Top3a |
| ENSMUSG00( | 274.498364 | 0.43367432 | 0.1809942 | 5.32E-04 | 0.05595652 | Rnasel |
| ENSMUSG00( | 1086.31053 | 0.43233336 | 0.14911545 | 1.42E-04 | 0.0252276 | Pdlim7 |
| ENSMUSG00( | 1194.43916 | 0.42967004 | 0.11494032 | 8.23E-06 | 0.00320068 | Gabarapl1 |
| ENSMUSG00( | 130.500207 | 0.42934628 | 0.20787731 | 0.0011183 | 0.08621611 | Clec10a |
| ENSMUSG00( | 270.121036 | 0.40817575 | 0.18005318 | 7.59E-04 | 0.06720439 | Ddias |
| ENSMUSG00( | 214.516209 | 0.40738139 | 0.18909299 | 9.05E-04 | 0.07484488 | Tmc8 |
| ENSMUSG00( | 769.045355 | 0.40342017 | 0.15035675 | 2.45E-04 | 0.03500886 | Rnf181 |
| ENSMUSG00( | 868.324281 | 0.3994633 | 0.17129396 | 6.30E-04 | 0.06042375 | Hspa4l |
| ENSMUSG00( | 411.787039 | 0.39562525 | 0.12470969 | 5.72E-05 | 0.01330399 | Fam219a |
| ENSMUSG00( | 325.667199 | 0.38983165 | 0.15314334 | 3.81E-04 | 0.04546435 | Il36g |
| ENSMUSG00( | 767.428019 | 0.3860505 | 0.07760723 | 2.74E-08 | 3.75E-05 | Hgsnat |
| ENSMUSG00( | 956.786803 | 0.38192225 | 0.09887245 | 4.93E-06 | 0.00211464 | P2rx7 |
| ENSMUSG00( | 257.412084 | 0.37812137 | 0.14269793 | 2.74E-04 | 0.03799638 | Zfp287 |
| ENSMUSG00( | 535.215558 | 0.37021222 | 0.11496052 | 5.08E-05 | 0.01268194 | Snx15 |
| ENSMUSG00( | 1281.21259 | 0.3634606 | 0.08978108 | 2.32E-06 | 0.00123422 | Slu7 |
| ENSMUSG00( | 225.926323 | 0.35258803 | 0.14486445 | 5.13E-04 | 0.05584654 | Slc49a4 |
| ENSMUSG00( | 431.604879 | 0.35091393 | 0.16609071 | 0.00107382 | 0.08567478 | Fbxo30 |
| ENSMUSG00( | 899.15283 | 0.3290413 | 0.14407715 | 8.14E-04 | 0.07047506 | Ppard |
| ENSMUSG00( | 702.98489 | 0.3268185 | 0.13429894 | 5.44E-04 | 0.05607956 | Rit1 |
| ENSMUSG00( | 799.678532 | 0.32521813 | 0.10608447 | 9.02E-05 | 0.0183103 | Galnt3 |
| ENSMUSG00( | 2968.06573 | 0.32307975 | 0.06009392 | 3.03E-09 | 6.93E-06 | Ubxn4 |
| ENSMUSG00( | 1797.46091 | 0.32217739 | 0.07982214 | 2.57E-06 | 0.00126202 | Ndel1 |
| ENSMUSG00( | 991.150567 | 0.3165461 | 0.11416193 | 2.13E-04 | 0.03176683 | Syt11 |
| ENSMUSG00( | 4750.05578 | 0.31526046 | 0.11693306 | 2.82E-04 | 0.03863499 | Ncf2 |
| ENSMUSG00( | 616.435537 | 0.31448478 | 0.12278261 | 3.49E-04 | 0.04470675 | Atf7ip |
| ENSMUSG00( | 449.539806 | 0.31404951 | 0.11018155 | 1.64E-04 | 0.02697118 | Zbtb43 |
| ENSMUSG00( | 357.819259 | 0.31271658 | 0.13918557 | 8.17E-04 | 0.07047506 | Sass6 |
| ENSMUSG00( | 786.72745 | 0.29934469 | 0.09978797 | 1.17E-04 | 0.02138924 | Adnp2 |
| ENSMUSG00( | 1275.96242 | 0.29747175 | 0.08912239 | 3.80E-05 | 0.01023421 | Cars |
| ENSMUSG00( | 375.653872 | 0.29738166 | 0.14494269 | 0.00135183 | 0.09815458 | Ptgr1 |
| ENSMUSG00( | 433.388953 | 0.29639935 | 0.11592788 | 3.71E-04 | 0.04502376 | Cep128 |
| ENSMUSG00( | 970.669397 | 0.29443315 | 0.12616439 | 7.02E-04 | 0.06507588 | Impact |
| ENSMUSG00( | 7555.80614 | 0.29308152 | 0.07511169 | 4.11E-06 | 0.00186988 | Akr1b3 |
| ENSMUSG00( | 216.947501 | 0.28911413 | 0.13475393 | 0.00103385 | 0.08296791 | Capn10 |
| ENSMUSG00( | 899.133751 | 0.28755722 | 0.09569191 | 1.24E-04 | 0.0223898 | Ppp2r3d |
| ENSMUSG00( | 594.602641 | 0.26985974 | 0.09440084 | 1.89E-04 | 0.02879562 | Mthfr |

| | | | | | | |
|---|---|---|---|---|---|---|
| ENSMUSG00( | 2137.52978 | 0.26572966 | 0.11212934 | 7.30E-04 | 0.0658813 | Esyt1 |
| ENSMUSG00( | 829.082007 | 0.26520155 | 0.08351855 | 7.74E-05 | 0.01610317 | Slc36a1 |
| ENSMUSG00( | 1029.26241 | 0.26284453 | 0.10226639 | 4.37E-04 | 0.05078734 | Rtel1 |
| ENSMUSG00( | 339.218729 | 0.2591203 | 0.12222014 | 0.00120998 | 0.09173814 | Rab10os |
| ENSMUSG00( | 512.053958 | 0.2591105 | 0.12389704 | 0.00122464 | 0.09233924 | Mynn |
| ENSMUSG00( | 1174.52914 | 0.25865605 | 0.08109701 | 6.98E-05 | 0.01474463 | Marf1 |
| ENSMUSG00( | 1170.05953 | 0.25725916 | 0.09904347 | 4.49E-04 | 0.05087763 | Leng8 |
| ENSMUSG00( | 3899.09317 | 0.25627044 | 0.0875342 | 1.47E-04 | 0.02590411 | Gaa |
| ENSMUSG00( | 722.851679 | 0.25502927 | 0.11186919 | 6.93E-04 | 0.06507588 | Them6 |
| ENSMUSG00( | 487.305394 | 0.24728039 | 0.10928148 | 9.32E-04 | 0.07654948 | Tbc1d17 |
| ENSMUSG00( | 4402.36864 | 0.24305191 | 0.06005563 | 2.34E-06 | 0.00123422 | Trp53 |
| ENSMUSG00( | 986.899473 | 0.24206851 | 0.11639078 | 0.00138787 | 0.09971577 | Slc25a10 |
| ENSMUSG00( | 639.854855 | 0.23694942 | 0.09597806 | 5.86E-04 | 0.05745655 | Tom1l2 |
| ENSMUSG00( | 732.28865 | 0.23300033 | 0.08932995 | 3.66E-04 | 0.04502376 | Dele1 |
| ENSMUSG00( | 4686.40106 | 0.23067761 | 0.0958974 | 7.97E-04 | 0.06966933 | Psmc3 |
| ENSMUSG00( | 380.610919 | 0.23035847 | 0.10585452 | 0.00113416 | 0.08664387 | Exd2 |
| ENSMUSG00( | 549.71908 | 0.22365846 | 0.09258292 | 6.80E-04 | 0.06432747 | Tmc6 |
| ENSMUSG00( | 1446.48973 | 0.21923049 | 0.07584857 | 1.89E-04 | 0.02879562 | E2f6 |
| ENSMUSG00( | 1769.0354 | 0.2187441 | 0.08335798 | 3.59E-04 | 0.04502376 | Srebf1 |
| ENSMUSG00( | 953.85299 | 0.21526661 | 0.09679455 | 0.00113648 | 0.08664387 | Pcx |
| ENSMUSG00( | 1044.88714 | 0.21386534 | 0.08339233 | 4.43E-04 | 0.05087763 | Sidt2 |
| ENSMUSG00( | 2930.33118 | 0.20943661 | 0.06034597 | 3.20E-05 | 0.00934288 | Napa |
| ENSMUSG00( | 4650.46679 | 0.20118281 | 0.04815948 | 1.99E-06 | 0.00123422 | Psmd3 |
| ENSMUSG00( | 957.262149 | 0.19479689 | 0.07473178 | 3.68E-04 | 0.04502376 | Ppp2r2d |
| ENSMUSG00( | 1286.75595 | 0.19335726 | 0.07189498 | 4.17E-04 | 0.04889479 | Rbm5 |
| ENSMUSG00( | 1976.45317 | 0.19323906 | 0.07456798 | 5.26E-04 | 0.05595652 | Ubxn1 |
| ENSMUSG00( | 2766.79835 | 0.18924367 | 0.07250033 | 5.01E-04 | 0.05498719 | Ube4b |
| ENSMUSG00( | 1714.50415 | 0.17493645 | 0.06968964 | 6.29E-04 | 0.06042375 | Retreg2 |
| ENSMUSG00( | 10187.3836 | 0.17360539 | 0.06736347 | 5.78E-04 | 0.05745655 | Psmd2 |
| ENSMUSG00( | 1018.48396 | 0.1722588 | 0.06133559 | 3.41E-04 | 0.04449873 | Dus3l |
| ENSMUSG00( | 1488.41085 | 0.16812415 | 0.06495653 | 5.19E-04 | 0.05595652 | Ranbp10 |
| ENSMUSG00( | 1719.36714 | 0.16576228 | 0.07084175 | 0.00111719 | 0.08621611 | Psmb2 |
| ENSMUSG00( | 1321.9163 | 0.15838783 | 0.06251337 | 7.19E-04 | 0.06543979 | Clpb |
| ENSMUSG00( | 42601.1039 | 0.12568672 | 0.03780899 | 9.13E-05 | 0.0183103 | Hsp90ab1 |
| ENSMUSG00( | 3490.36736 | 0.12425023 | 0.0504219 | 0.00109529 | 0.08621611 | 9530068E07F |
| ENSMUSG00( | 3273.03039 | 0.1214477 | 0.04890808 | 0.00101511 | 0.08242829 | Rnf10 |
| ENSMUSG00( | 20.5614142 | 0.01692574 | 0.05072857 | 5.63E-04 | 0.05745655 | Gm43375 |
| ENSMUSG00( | 26.883157 | -0.0197856 | 0.05244103 | 7.49E-04 | 0.06675598 | Ceacam19 |
| ENSMUSG00( | 40.7481383 | -0.0268231 | 0.05808059 | 8.29E-04 | 0.07063389 | Oas1b |
| ENSMUSG00( | 93.6106657 | -0.0459078 | 0.08860773 | 8.59E-04 | 0.07184485 | Tlr3 |
| ENSMUSG00( | 5788.74515 | -0.1427478 | 0.05985075 | 0.00126084 | 0.09352697 | Adgre1 |

| | | | | | | |
|---|---|---|---|---|---|---|
| ENSMUSG00( | 4364.60126 | -0.1460531 | 0.04624557 | 9.46E-05 | 0.0183103 | G3bp2 |
| ENSMUSG00( | 2100.86368 | -0.1544099 | 0.05787503 | 5.34E-04 | 0.05595652 | Hnrnpll |
| ENSMUSG00( | 10041.2278 | -0.159998 | 0.06300275 | 5.78E-04 | 0.05745655 | Cdv3 |
| ENSMUSG00( | 3320.86849 | -0.1639692 | 0.06797202 | 7.74E-04 | 0.06808968 | Drg1 |
| ENSMUSG00( | 1277.10771 | -0.1808323 | 0.08401642 | 0.00136814 | 0.09881587 | Prkaca |
| ENSMUSG00( | 2693.30709 | -0.1860765 | 0.0651475 | 3.44E-04 | 0.04449873 | Anln |
| ENSMUSG00( | 2669.95503 | -0.1867263 | 0.06640397 | 2.27E-04 | 0.03310926 | Snx18 |
| ENSMUSG00( | 1007.52089 | -0.1881704 | 0.07598366 | 7.18E-04 | 0.06543979 | Mpzl1 |
| ENSMUSG00( | 10937.6648 | -0.1891469 | 0.07206095 | 4.51E-04 | 0.05087763 | Slc15a3 |
| ENSMUSG00( | 1346.64699 | -0.1910688 | 0.08828758 | 0.00109466 | 0.08621611 | Dusp7 |
| ENSMUSG00( | 2582.1001 | -0.1927635 | 0.05838564 | 5.21E-05 | 0.01275521 | Jpt2 |
| ENSMUSG00( | 30551.0878 | -0.19459 | 0.08809568 | 0.00123912 | 0.09241526 | Eif4g2 |
| ENSMUSG00( | 4099.15866 | -0.1964943 | 0.06835937 | 2.01E-04 | 0.03030075 | Gart |
| ENSMUSG00( | 1915.55564 | -0.2124047 | 0.06113723 | 2.62E-05 | 0.00856778 | Rapgef5 |
| ENSMUSG00( | 1259.77617 | -0.2153106 | 0.06662144 | 5.66E-05 | 0.01330399 | Desi2 |
| ENSMUSG00( | 1150.49876 | -0.2167139 | 0.09367186 | 6.58E-04 | 0.0627493 | Gpd1l |
| ENSMUSG00( | 920.685627 | -0.2185159 | 0.08949996 | 7.38E-04 | 0.06614919 | Rpp14 |
| ENSMUSG00( | 672.021371 | -0.2205555 | 0.07992661 | 3.15E-04 | 0.04191041 | St3gal4 |
| ENSMUSG00( | 2100.71437 | -0.2423062 | 0.07404557 | 5.30E-05 | 0.01275521 | Lcp2 |
| ENSMUSG00( | 1258.98949 | -0.2457656 | 0.11747715 | 0.00129148 | 0.09427104 | Aspm |
| ENSMUSG00( | 1606.94619 | -0.2582175 | 0.08102654 | 6.97E-05 | 0.01474463 | Cd47 |
| ENSMUSG00( | 1238.4652 | -0.2621856 | 0.07653952 | 2.82E-05 | 0.00860525 | Tnfaip8 |
| ENSMUSG00( | 369.688753 | -0.2663505 | 0.12166794 | 0.0010255 | 0.08278213 | Glrx2 |
| ENSMUSG00( | 4744.499 | -0.276435 | 0.06582264 | 1.07E-06 | 8.19E-04 | Nceh1 |
| ENSMUSG00( | 319.675431 | -0.2923001 | 0.13042519 | 8.53E-04 | 0.07184485 | Cep89 |
| ENSMUSG00( | 400.968192 | -0.2925799 | 0.10978057 | 3.05E-04 | 0.04103182 | Qpct |
| ENSMUSG00( | 1783.32284 | -0.2992234 | 0.08926391 | 3.41E-05 | 0.00976305 | Fmnl2 |
| ENSMUSG00( | 678.540214 | -0.3055799 | 0.09634003 | 6.49E-05 | 0.01435391 | Metrnl |
| ENSMUSG00( | 392.758995 | -0.3109414 | 0.09858424 | 6.19E-05 | 0.01416876 | Icosl |
| ENSMUSG00( | 245.825019 | -0.3154242 | 0.12756779 | 4.56E-04 | 0.05087763 | Slc25a40 |
| ENSMUSG00( | 806.739449 | -0.321974 | 0.09620774 | 3.52E-05 | 0.00984783 | Kif2c |
| ENSMUSG00( | 339.198886 | -0.3382749 | 0.15373557 | 8.86E-04 | 0.07369938 | Mmachc |
| ENSMUSG00( | 708.65974 | -0.3466359 | 0.11992646 | 0.00016624 | 0.02697118 | Fam241a |
| ENSMUSG00( | 449.008797 | -0.3486669 | 0.12218649 | 1.67E-04 | 0.02697118 | Hacd2 |
| ENSMUSG00( | 1096.21288 | -0.3590614 | 0.12979771 | 2.17E-04 | 0.03205886 | Flrt2 |
| ENSMUSG00( | 563.479642 | -0.3637756 | 0.16409957 | 8.35E-04 | 0.07069184 | Depdc1a |
| ENSMUSG00( | 1336.46898 | -0.3678892 | 0.10750074 | 2.79E-05 | 0.00860525 | Fcgr1 |
| ENSMUSG00( | 2492.01485 | -0.380806 | 0.08005288 | 1.15E-07 | 1.25E-04 | Serpinb6b |
| ENSMUSG00( | 472.677916 | -0.382375 | 0.1630214 | 5.43E-04 | 0.05607956 | Usp18 |
| ENSMUSG00( | 2411.28829 | -0.3850883 | 0.09384916 | 1.83E-06 | 0.00119333 | Ccnd1 |
| ENSMUSG00( | 2139.46268 | -0.3859211 | 0.11620271 | 3.62E-05 | 0.00994241 | Slc39a10 |

| | | | | | | |
|---|---|---|---|---|---|---|
| ENSMUSG00( | 1653.26904 | -0.3877843 | 0.16099446 | 5.26E-04 | 0.05595652 | Slc7a2 |
| ENSMUSG00( | 1959.99687 | -0.3924398 | 0.10979434 | 1.26E-05 | 0.00455476 | Cenpf |
| ENSMUSG00( | 737.086285 | -0.3985835 | 0.09652015 | 2.10E-06 | 0.00123422 | Ifi204 |
| ENSMUSG00( | 353.8994 | -0.4012076 | 0.20186743 | 0.00128199 | 0.09427104 | Cdkn3 |
| ENSMUSG00( | 4468.96981 | -0.4236071 | 0.1018464 | 1.41E-06 | 0.00101587 | Ccna2 |
| ENSMUSG00( | 1352.77871 | -0.42394 | 0.14851086 | 1.51E-04 | 0.02628125 | Pimreg |
| ENSMUSG00( | 297.686373 | -0.4311887 | 0.13787586 | 6.45E-05 | 0.01435391 | Phf19 |
| ENSMUSG00( | 15951.1115 | -0.4336391 | 0.1081516 | 2.30E-06 | 0.00123422 | Tnfrsf1b |
| ENSMUSG00( | 159.937337 | -0.4403634 | 0.16460067 | 2.47E-04 | 0.03500886 | Igtp |
| ENSMUSG00( | 321.538091 | -0.4538051 | 0.13826395 | 4.03E-05 | 0.01064461 | Fam78b |
| ENSMUSG00( | 461.594294 | -0.45477 | 0.17648861 | 2.94E-04 | 0.0399837 | Bcl2a1b |
| ENSMUSG00( | 900.388441 | -0.4610341 | 0.13617058 | 2.51E-05 | 0.00840865 | Bcl2a1d |
| ENSMUSG00( | 267.285197 | -0.47666 | 0.15878366 | 9.68E-05 | 0.018444 | Cdkn2d |
| ENSMUSG00( | 174.820342 | -0.4768638 | 0.20395043 | 5.83E-04 | 0.05745655 | Ifi211 |
| ENSMUSG00( | 374.498484 | -0.480687 | 0.16034864 | 9.85E-05 | 0.01852108 | Ifit2 |
| ENSMUSG00( | 415.923653 | -0.492224 | 0.12618786 | 4.01E-06 | 0.00186988 | Zbed3 |
| ENSMUSG00( | 417.300295 | -0.509784 | 0.20426698 | 3.74E-04 | 0.04502376 | Pif1 |
| ENSMUSG00( | 7139.60807 | -0.5107214 | 0.16439221 | 6.73E-05 | 0.01467005 | Il1a |
| ENSMUSG00( | 5246.57234 | -0.5258508 | 0.12990795 | 2.45E-06 | 0.00124406 | Ccl9 |
| ENSMUSG00( | 415.450825 | -0.5446671 | 0.15460714 | 1.59E-05 | 0.00560536 | Ifi203 |
| ENSMUSG00( | 300.715593 | -0.5526179 | 0.19797951 | 1.74E-04 | 0.02742247 | Gbp2 |
| ENSMUSG00( | 1293.46044 | -0.5576464 | 0.11120791 | 1.68E-08 | 2.56E-05 | Sapcd2 |
| ENSMUSG00( | 222.19915 | -0.5891311 | 0.2368955 | 4.04E-04 | 0.04776166 | Cd38 |
| ENSMUSG00( | 130.510103 | -0.6652506 | 0.26008187 | 3.70E-04 | 0.04502376 | Smtn |
| ENSMUSG00( | 175.975641 | -0.6701916 | 0.22874857 | 1.09E-04 | 0.02028227 | Gbp7 |
| ENSMUSG00( | 293.755827 | -0.717894 | 0.19037421 | 6.04E-06 | 0.00245382 | Ifit3 |
| ENSMUSG00( | 75.852775 | -0.7531934 | 0.3360433 | 6.97E-04 | 0.06507588 | Ccl22 |
| ENSMUSG00( | 6280.84599 | -0.7929231 | 0.11906318 | 1.23E-12 | 5.65E-09 | Il1b |

e_name

Rik

ࢱik
