## Supplemental information, figure compressed, table 1-5 for "Role of copper during microglial inflammation": Supplemental Table 3-compressed.pdf

| Gene_Id | baseMean | log2FoldChar | lfcSE | pvalue | padj | external_gene |
|---|---|---|---|---|---|---|
| ENSMUSG00( | 54.6281499 | 7.54347965 | 2.20766179 | 3.75E-09 | 5.44E-08 | 2310002L09R |
| ENSMUSG00( | 60.1582402 | 5.41831068 | 0.93521412 | 2.15E-08 | 2.75E-07 | BC021767 |
| ENSMUSG00( | 5.79491455 | 4.66099478 | 2.11160013 | 8.88E-05 | 5.61E-04 | Ppef2 |
| ENSMUSG00( | 6.18793027 | 4.5313748 | 2.23583615 | 2.49E-04 | 0.00140471 | AI606473 |
| ENSMUSG00( | 74.2138913 | 4.50698408 | 0.390427 | 2.69E-31 | 4.59E-29 | Mgst1 |
| ENSMUSG00( | 1.91492289 | 4.46849368 | 2.51063159 | 7.07E-04 | 0.0035576 | Slc10a6 |
| ENSMUSG00( | 7.85575179 | 4.35231831 | 2.06841321 | 1.48E-04 | 8.85E-04 | Lrrc17 |
| ENSMUSG00( | 4.43872266 | 4.34863362 | 2.23632837 | 3.50E-04 | 0.00189562 | Hes7 |
| ENSMUSG00( | 6.35196385 | 4.28310671 | 2.16847075 | 2.67E-04 | 0.00149222 | |
| ENSMUSG00( | 307.472995 | 4.25012421 | 0.25150983 | 8.76E-65 | 7.53E-62 | Iqgap2 |
| ENSMUSG00( | 3.36417086 | 4.18307784 | 2.27324472 | 5.29E-04 | 0.00274549 | Vangl1 |
| ENSMUSG00( | 2.30278249 | 4.13701834 | 2.55590811 | 0.00121368 | 0.00572883 | Gm9710 |
| ENSMUSG00( | 190.854256 | 4.08746787 | 0.26716535 | 6.92E-54 | 4.46E-51 | Spink5 |
| ENSMUSG00( | 2.51398847 | 4.03291447 | 2.26956209 | 6.57E-04 | 0.00332442 | Lgr4 |
| ENSMUSG00( | 154.586545 | 3.95986356 | 0.35793556 | 1.28E-29 | 1.94E-27 | Marco |
| ENSMUSG00( | 3.32822467 | 3.95645205 | 2.38285772 | 0.00101735 | 0.00491775 | Cd177 |
| ENSMUSG00( | 5.18375651 | 3.82705584 | 2.11943156 | 5.61E-04 | 0.00288727 | |
| ENSMUSG00( | 20.2127241 | 3.67111097 | 0.84399877 | 2.41E-06 | 2.13E-05 | Gm10134 |
| ENSMUSG00( | 10.9629646 | 3.65932274 | 2.05059658 | 6.01E-04 | 0.00307363 | Gm15513 |
| ENSMUSG00( | 5.3296218 | 3.65297999 | 1.5865051 | 9.55E-04 | 0.00465559 | Gfi1b |
| ENSMUSG00( | 2.16887858 | 3.62413254 | 2.08524681 | 0.00264656 | 0.01126517 | Tmem267 |
| ENSMUSG00( | 82.7281523 | 3.54404446 | 0.45608039 | 5.40E-16 | 2.08E-14 | Hp |
| ENSMUSG00( | 3.07200687 | 3.46991938 | 1.35565528 | 0.00108537 | 0.00520615 | Bpifc |
| ENSMUSG00( | 3.18972504 | 3.43006013 | 2.31092357 | 0.00160825 | 0.00729353 | |
| ENSMUSG00( | 62.1298058 | 3.39553635 | 0.43645794 | 2.08E-15 | 7.47E-14 | Selenbp2 |
| ENSMUSG00( | 2.25627359 | 3.38992132 | 2.4176996 | 0.00200676 | 0.0088353 | Rnf151 |
| ENSMUSG00( | 48.2621061 | 3.38285351 | 0.53721962 | 6.70E-11 | 1.27E-09 | Adgb |
| ENSMUSG00( | 228.310471 | 3.37118009 | 0.62040706 | 2.49E-09 | 3.74E-08 | Cx3cr1 |
| ENSMUSG00( | 2.49450141 | 3.33987402 | 1.69059612 | 0.00205262 | 0.00902404 | Pkp3 |
| ENSMUSG00( | 45.1133873 | 3.30026008 | 0.50852208 | 1.47E-11 | 3.07E-10 | Prnp |
| ENSMUSG00( | 5.13644598 | 3.28537019 | 2.11282427 | 0.00128045 | 0.00599543 | 9330179D12F |
| ENSMUSG00( | 11.2538419 | 3.27020018 | 1.22409494 | 8.68E-04 | 0.0042811 | Pacsin1 |
| ENSMUSG00( | 61.4619316 | 3.15138799 | 0.34481055 | 1.11E-20 | 7.42E-19 | Osbpl10 |
| ENSMUSG00( | 3.95203605 | 3.13023965 | 2.41141725 | 0.00243137 | 0.01048005 | Fam222a |
| ENSMUSG00( | 15.5185356 | 3.07341471 | 0.79504441 | 6.56E-06 | 5.30E-05 | Slc35f1 |
| ENSMUSG00( | 69.4592453 | 3.04445249 | 0.36238814 | 2.79E-18 | 1.44E-16 | Glis3 |
| ENSMUSG00( | 2.12605707 | 3.01374144 | 1.630135 | 0.00280295 | 0.01182501 | Rab33a |
| ENSMUSG00( | 10.1568592 | 2.97616168 | 0.67922196 | 1.09E-06 | 1.03E-05 | Fpr1 |
| ENSMUSG00( | 122.198332 | 2.92401031 | 0.39526622 | 8.08E-15 | 2.70E-13 | Sgms2 |
| ENSMUSG00( | 79.7856732 | 2.91014137 | 0.4538958 | 2.19E-11 | 4.45E-10 | Pgbd5 |

| | | | | | | |
|---|---|---|---|---|---|---|
| ENSMUSG00( | 2.67783227 | 2.90822185 | 1.78718544 | 0.00381317 | 0.01544153 | Tm9sf5 |
| ENSMUSG00( | 10.3083782 | 2.9076966 | 0.81227433 | 5.44E-05 | 3.60E-04 | Clca4a |
| ENSMUSG00( | 60.4805 | 2.88685166 | 0.29092759 | 4.40E-24 | 4.18E-22 | Sptbn1 |
| ENSMUSG00( | 25.2275085 | 2.88677685 | 0.67565729 | 2.19E-06 | 1.95E-05 | Ptgfrn |
| ENSMUSG00( | 161.02479 | 2.85382569 | 0.25462346 | 2.76E-30 | 4.26E-28 | Ntng2 |
| ENSMUSG00( | 1075.65381 | 2.803052 | 0.13256516 | 2.37E-100 | 4.76E-97 | Rassf2 |
| ENSMUSG00( | 122.591905 | 2.78411384 | 0.30464974 | 8.08E-21 | 5.57E-19 | Gda |
| ENSMUSG00( | 110.472745 | 2.78259021 | 0.73833011 | 7.01E-06 | 5.62E-05 | Gsta3 |
| ENSMUSG00( | 11.9207092 | 2.77525711 | 0.79642383 | 3.33E-05 | 2.31E-04 | Gstt4 |
| ENSMUSG00( | 1288.84239 | 2.72736693 | 0.23096255 | 2.59E-33 | 5.02E-31 | Cytip |
| ENSMUSG00( | 535.460446 | 2.68557647 | 0.20615446 | 6.48E-40 | 2.09E-37 | Slc8a1 |
| ENSMUSG00( | 9.67390877 | 2.64493976 | 0.96956712 | 7.71E-04 | 0.00384964 | Cracr2a |
| ENSMUSG00( | 39.7321223 | 2.63158322 | 0.37256748 | 1.17E-13 | 3.29E-12 | Gm20658 |
| ENSMUSG00( | 13.3847058 | 2.6207618 | 0.81726878 | 9.18E-05 | 5.76E-04 | Gm37261 |
| ENSMUSG00( | 8.08815902 | 2.61832375 | 1.10886503 | 0.00258312 | 0.01103154 | Poln |
| ENSMUSG00( | 54.4240787 | 2.61666527 | 0.36853009 | 8.95E-14 | 2.57E-12 | Gstm2 |
| ENSMUSG00( | 15.3633918 | 2.61425815 | 0.94314113 | 3.57E-04 | 0.00192546 | Cpxm1 |
| ENSMUSG00( | 24.6306123 | 2.60211366 | 0.47541808 | 5.84E-09 | 8.19E-08 | Smoc1 |
| ENSMUSG00( | 213.334312 | 2.5993335 | 0.39541878 | 3.05E-12 | 6.97E-11 | Fry |
| ENSMUSG00( | 41.9040199 | 2.59246799 | 0.4914161 | 9.01E-09 | 1.22E-07 | Iqsec3 |
| ENSMUSG00( | 15.3665845 | 2.53150026 | 0.68151302 | 1.77E-05 | 1.30E-04 | Sirpb1c |
| ENSMUSG00( | 16.8804173 | 2.50874226 | 0.68697479 | 2.01E-05 | 1.46E-04 | Rasgrp2 |
| ENSMUSG00( | 26.7187587 | 2.47080315 | 0.45444288 | 4.95E-09 | 7.03E-08 | Tbx15 |
| ENSMUSG00( | 69.6443201 | 2.4684294 | 0.30886338 | 1.06E-16 | 4.46E-15 | Tgm2 |
| ENSMUSG00( | 607.285725 | 2.44327056 | 0.2320132 | 3.47E-27 | 4.17E-25 | Mgst2 |
| ENSMUSG00( | 9.33326725 | 2.43950569 | 0.72444612 | 5.66E-05 | 3.73E-04 | Gm16556 |
| ENSMUSG00( | 3.8847758 | 2.36282309 | 1.58882208 | 0.00562948 | 0.02151419 | Atp8b3 |
| ENSMUSG00( | 21.1510933 | 2.36139752 | 0.67957789 | 2.50E-05 | 1.79E-04 | Mt2 |
| ENSMUSG00( | 367.999237 | 2.36134059 | 0.2213175 | 8.62E-28 | 1.11E-25 | Treml4 |
| ENSMUSG00( | 3.58753855 | 2.35793943 | 1.53220919 | 0.00571131 | 0.02179004 | Htra3 |
| ENSMUSG00( | 10.1575989 | 2.35725446 | 0.95964061 | 0.00157699 | 0.00718109 | Rgs12 |
| ENSMUSG00( | 54.9493308 | 2.30326082 | 0.43263307 | 9.43E-09 | 1.27E-07 | Adrb1 |
| ENSMUSG00( | 7.59011299 | 2.2975867 | 1.4843879 | 0.00555299 | 0.02125334 | H1f5 |
| ENSMUSG00( | 3.99315013 | 2.29131365 | 1.19714985 | 0.00335855 | 0.0138347 | Gm4804 |
| ENSMUSG00( | 781.372269 | 2.28928893 | 0.21280333 | 3.57E-28 | 4.74E-26 | Xdh |
| ENSMUSG00( | 2.76051516 | 2.27595255 | 2.5193511 | 0.00676637 | 0.02508442 | Grhl2 |
| ENSMUSG00( | 168.398051 | 2.26783759 | 0.3276357 | 2.75E-13 | 7.29E-12 | Mrc1 |
| ENSMUSG00( | 198.018869 | 2.24806235 | 0.34770462 | 6.52E-12 | 1.42E-10 | Cmbl |
| ENSMUSG00( | 984.559571 | 2.23918719 | 0.32125387 | 1.83E-13 | 5.01E-12 | Dmpk |
| ENSMUSG00( | 3.5762313 | 2.23208616 | 1.28825969 | 0.00468621 | 0.01837965 | Tnfrsf21 |
| ENSMUSG00( | 131.730368 | 2.19289215 | 0.25468034 | 9.35E-19 | 4.97E-17 | Tmem119 |

| | | | | | | |
|---|---|---|---|---|---|---|
| ENSMUSG00( | 7.452321 | 2.18896734 | 0.86323097 | 8.36E-04 | 0.00413312 | Nim1k |
| ENSMUSG00( | 36.8718595 | 2.18206488 | 0.35357823 | 5.47E-11 | 1.05E-09 | Akr1c12 |
| ENSMUSG00( | 2.85539533 | 2.17781238 | 1.56739781 | 0.00684205 | 0.02532344 | Gm12444 |
| ENSMUSG00( | 405.198642 | 2.1745432 | 0.23443948 | 1.62E-21 | 1.18E-19 | S1pr1 |
| ENSMUSG00( | 3.02112381 | 2.16621347 | 1.95000672 | 0.00611877 | 0.02307663 | Tex19.1 |
| ENSMUSG00( | 20.6848772 | 2.11074301 | 0.74468602 | 4.04E-04 | 0.00215404 | Gm43848 |
| ENSMUSG00( | 79.9568789 | 2.08403591 | 0.28991344 | 6.04E-14 | 1.76E-12 | Dlg3 |
| ENSMUSG00( | 188.42149 | 2.07432644 | 0.27156517 | 2.34E-15 | 8.37E-14 | Mertk |
| ENSMUSG00( | 4.01166633 | 2.05177508 | 1.95478827 | 0.00835891 | 0.03001534 | Rhoj |
| ENSMUSG00( | 19.2665571 | 2.03278075 | 0.74591795 | 4.77E-04 | 0.00250344 | Mlph |
| ENSMUSG00( | 82.2878439 | 2.0173894 | 0.30870524 | 4.80E-12 | 1.07E-10 | Evl |
| ENSMUSG00( | 32.9246436 | 2.00936581 | 0.41777185 | 1.29E-07 | 1.42E-06 | Nuak1 |
| ENSMUSG00( | 36.0625345 | 2.00680837 | 0.37777748 | 1.02E-08 | 1.38E-07 | Rab39 |
| ENSMUSG00( | 297.270384 | 1.99639353 | 0.18645199 | 9.49E-28 | 1.22E-25 | Kank2 |
| ENSMUSG00( | 46.0810002 | 1.99010941 | 0.40397038 | 6.32E-08 | 7.48E-07 | Il15ra |
| ENSMUSG00( | 1204.40891 | 1.98933227 | 0.30814361 | 7.03E-12 | 1.52E-10 | Ppp1r12b |
| ENSMUSG00( | 14.0694813 | 1.98580666 | 0.57210858 | 4.56E-05 | 3.07E-04 | Gm32089 |
| ENSMUSG00( | 12.2240609 | 1.97176124 | 1.20256902 | 0.00554082 | 0.02122473 | Sgpp2 |
| ENSMUSG00( | 835.289689 | 1.96917981 | 0.18228384 | 2.53E-28 | 3.54E-26 | Deptor |
| ENSMUSG00( | 1170.92151 | 1.96755944 | 0.17251312 | 3.16E-31 | 5.19E-29 | Itgal |
| ENSMUSG00( | 53.4658549 | 1.96006573 | 0.36982848 | 8.72E-09 | 1.19E-07 | Apba1 |
| ENSMUSG00( | 94.0611222 | 1.95883929 | 0.71036837 | 5.41E-04 | 0.00279622 | Asb10 |
| ENSMUSG00( | 20.3851543 | 1.95068713 | 0.65727788 | 2.58E-04 | 0.0014508 | Tlr8 |
| ENSMUSG00( | 3.68317132 | 1.93797448 | 1.30732653 | 0.00664741 | 0.02472459 | Kif1a |
| ENSMUSG00( | 48.8827407 | 1.9287666 | 0.37693843 | 2.31E-08 | 2.93E-07 | Id1 |
| ENSMUSG00( | 417.503307 | 1.92263983 | 0.18243402 | 4.98E-27 | 5.92E-25 | Rab3il1 |
| ENSMUSG00( | 69.1565506 | 1.90921767 | 0.3758055 | 2.51E-08 | 3.16E-07 | Clec4a2 |
| ENSMUSG00( | 6.60579252 | 1.9078297 | 1.31120472 | 0.01018503 | 0.03543191 | Gm37795 |
| ENSMUSG00( | 451.046309 | 1.90449032 | 0.21512808 | 5.70E-20 | 3.55E-18 | Cebpa |
| ENSMUSG00( | 65.2902276 | 1.90118748 | 0.34009126 | 1.65E-09 | 2.55E-08 | Ak1 |
| ENSMUSG00( | 45.7439231 | 1.89475472 | 0.41998711 | 6.37E-07 | 6.24E-06 | Asb13 |
| ENSMUSG00( | 41.0297687 | 1.8889584 | 0.31934837 | 2.79E-10 | 4.79E-09 | Fam114a1 |
| ENSMUSG00( | 436.049428 | 1.88132617 | 0.16394111 | 1.47E-31 | 2.55E-29 | Ppm1e |
| ENSMUSG00( | 296.461509 | 1.87841651 | 0.16615029 | 1.05E-30 | 1.68E-28 | Sla |
| ENSMUSG00( | 2276.91235 | 1.85979356 | 0.27287716 | 7.67E-13 | 1.91E-11 | Igf1 |
| ENSMUSG00( | 1520.90432 | 1.85293526 | 0.1143862 | 2.78E-60 | 2.18E-57 | Rab32 |
| ENSMUSG00( | 4.26505608 | 1.84378349 | 1.27587096 | 0.01096211 | 0.03761381 | Gm41041 |
| ENSMUSG00( | 636.17304 | 1.84108313 | 0.1886087 | 1.24E-23 | 1.11E-21 | Scamp5 |
| ENSMUSG00( | 126.179276 | 1.83878777 | 0.20102671 | 6.23E-21 | 4.36E-19 | Mkx |
| ENSMUSG00( | 36.8773308 | 1.83825015 | 0.41559338 | 6.67E-07 | 6.51E-06 | Ccdc125 |
| ENSMUSG00( | 48.8276963 | 1.83211532 | 0.3501916 | 1.38E-08 | 1.81E-07 | Dok4 |

| | | | | | | |
|---|---|---|---|---|---|---|
| ENSMUSG00( | 14.1596668 | 1.83119558 | 0.55751627 | 7.86E-05 | 5.02E-04 | 9830132P13F |
| ENSMUSG00( | 151.139214 | 1.82978711 | 0.2523915 | 3.28E-14 | 1.01E-12 | Mxd4 |
| ENSMUSG00( | 57.5852635 | 1.82320408 | 0.35777031 | 2.79E-08 | 3.49E-07 | Mapre3 |
| ENSMUSG00( | 280.282666 | 1.79381373 | 0.14883303 | 2.12E-34 | 4.51E-32 | Emilin1 |
| ENSMUSG00( | 6.81404826 | 1.78585449 | 0.95421933 | 0.00246433 | 0.01059937 | Efcab6 |
| ENSMUSG00( | 10.6467466 | 1.78483208 | 0.89196918 | 0.00378153 | 0.01532714 | Ovgp1 |
| ENSMUSG00( | 196.320668 | 1.78328767 | 0.22919163 | 5.35E-16 | 2.07E-14 | Mcf2l |
| ENSMUSG00( | 115.700914 | 1.77713084 | 0.25313896 | 1.88E-13 | 5.13E-12 | Ticam2 |
| ENSMUSG00( | 27.5895187 | 1.77579979 | 0.57287535 | 1.08E-04 | 6.66E-04 | F2r |
| ENSMUSG00( | 37.2954685 | 1.77128228 | 0.44834382 | 5.76E-06 | 4.72E-05 | Afmid |
| ENSMUSG00( | 3463.34007 | 1.77101644 | 0.38923238 | 3.37E-07 | 3.46E-06 | Itga4 |
| ENSMUSG00( | 36.2079993 | 1.77101029 | 0.53356812 | 5.63E-05 | 3.72E-04 | Msi1 |
| ENSMUSG00( | 82.8193484 | 1.76869871 | 0.25151464 | 1.90E-13 | 5.15E-12 | Cped1 |
| ENSMUSG00( | 146.192968 | 1.76674519 | 0.19179031 | 3.07E-21 | 2.19E-19 | Zic2 |
| ENSMUSG00( | 214.960435 | 1.76224727 | 0.19190137 | 3.34E-21 | 2.38E-19 | Itga9 |
| ENSMUSG00( | 36.5004742 | 1.75747739 | 0.56983058 | 1.11E-04 | 6.87E-04 | |
| ENSMUSG00( | 292.763474 | 1.74930499 | 0.18276322 | 9.57E-23 | 7.65E-21 | Engase |
| ENSMUSG00( | 19.049598 | 1.73581268 | 0.51207773 | 5.81E-05 | 3.82E-04 | Garnl3 |
| ENSMUSG00( | 597.356514 | 1.7348861 | 0.11513057 | 1.97E-52 | 1.11E-49 | Cyfip2 |
| ENSMUSG00( | 14.8861643 | 1.73042932 | 0.74781028 | 0.00171812 | 0.00772834 | Slc9a9 |
| ENSMUSG00( | 42.4458716 | 1.72030452 | 0.53885975 | 1.00E-04 | 6.25E-04 | |
| ENSMUSG00( | 30.9375024 | 1.71992791 | 0.57543692 | 1.67E-04 | 9.84E-04 | U90926 |
| ENSMUSG00( | 72.9114469 | 1.71305549 | 0.4072817 | 2.67E-06 | 2.32E-05 | Ttc39a |
| ENSMUSG00( | 408.326118 | 1.70793214 | 0.15715102 | 1.82E-28 | 2.60E-26 | Abcd2 |
| ENSMUSG00( | 22.4310282 | 1.70348167 | 0.54740856 | 1.30E-04 | 7.92E-04 | Gbgt1 |
| ENSMUSG00( | 46.8996724 | 1.68923877 | 0.27566809 | 7.34E-11 | 1.38E-09 | Trem3 |
| ENSMUSG00( | 4.64973013 | 1.68917352 | 1.15950542 | 0.00886131 | 0.0315125 | Syt6 |
| ENSMUSG00( | 47.5549918 | 1.68680792 | 0.42947489 | 7.51E-06 | 5.99E-05 | Abhd15 |
| ENSMUSG00( | 23.2821418 | 1.6807191 | 0.62767019 | 4.05E-04 | 0.00216083 | Cd300e |
| ENSMUSG00( | 51.1536985 | 1.66478474 | 0.35633959 | 2.91E-07 | 3.02E-06 | Vangl2 |
| ENSMUSG00( | 132.590446 | 1.66328039 | 0.19865624 | 4.99E-18 | 2.44E-16 | Nxf7 |
| ENSMUSG00( | 140.989878 | 1.66307345 | 0.39640922 | 1.74E-06 | 1.57E-05 | Pdlim4 |
| ENSMUSG00( | 294.779826 | 1.66281432 | 0.19154923 | 2.65E-19 | 1.47E-17 | Irak3 |
| ENSMUSG00( | 164.625664 | 1.66149653 | 0.24001093 | 3.42E-13 | 8.95E-12 | Ppp2r3a |
| ENSMUSG00( | 45.4991094 | 1.6529705 | 0.29907595 | 2.56E-09 | 3.84E-08 | Plb1 |
| ENSMUSG00( | 486.324677 | 1.65048668 | 0.21267212 | 7.53E-16 | 2.87E-14 | Ogfrl1 |
| ENSMUSG00( | 62.9193946 | 1.64966422 | 0.30093824 | 3.94E-09 | 5.69E-08 | Inpp5j |
| ENSMUSG00( | 15.0428897 | 1.6496594 | 0.45848416 | 2.14E-05 | 1.55E-04 | Gm8818 |
| ENSMUSG00( | 1941.89959 | 1.64339725 | 0.15334713 | 6.64E-28 | 8.76E-26 | Sec24d |
| ENSMUSG00( | 17.4002454 | 1.63974758 | 0.70522051 | 0.00102204 | 0.00493645 | Slc7a15 |
| ENSMUSG00( | 151.407571 | 1.63541415 | 0.19992955 | 2.22E-17 | 1.01E-15 | Spsb4 |

| | | | | | | |
|---|---|---|---|---|---|---|
| ENSMUSG00( | 65.4903268 | 1.63201934 | 0.44694575 | 1.67E-05 | 1.23E-04 | Gm19026 |
| ENSMUSG00( | 36.8593034 | 1.63018606 | 0.5693558 | 3.79E-04 | 0.00202902 | Ssbp2 |
| ENSMUSG00( | 12.6020132 | 1.62722286 | 0.69549826 | 0.00113547 | 0.00541053 | Trpv4 |
| ENSMUSG00( | 10.0170783 | 1.62375983 | 0.89592446 | 0.00452847 | 0.01781127 | Gm44509 |
| ENSMUSG00( | 788.498451 | 1.62319238 | 0.12554271 | 2.50E-39 | 7.77E-37 | Ivd |
| ENSMUSG00( | 720.300477 | 1.62290254 | 0.18444157 | 1.06E-19 | 6.38E-18 | Lmo4 |
| ENSMUSG00( | 102.78363 | 1.620384 | 0.27034517 | 1.65E-10 | 2.94E-09 | Bmf |
| ENSMUSG00( | 386.934223 | 1.61361428 | 0.19446784 | 8.28E-18 | 3.92E-16 | Naaa |
| ENSMUSG00( | 12.4289958 | 1.61286971 | 0.5097166 | 1.11E-04 | 6.87E-04 | Naip3 |
| ENSMUSG00( | 85.9370789 | 1.60750971 | 0.28991035 | 2.81E-09 | 4.17E-08 | Sbk1 |
| ENSMUSG00( | 2456.80227 | 1.60426138 | 0.29609735 | 4.28E-09 | 6.13E-08 | Slc43a2 |
| ENSMUSG00( | 27.9942469 | 1.60114297 | 0.43121144 | 1.45E-05 | 1.09E-04 | Fndc9 |
| ENSMUSG00( | 47.3760552 | 1.60033924 | 0.32379918 | 7.37E-08 | 8.59E-07 | Svil |
| ENSMUSG00( | 7535.32109 | 1.59662101 | 0.17722028 | 2.98E-20 | 1.89E-18 | Mt1 |
| ENSMUSG00( | 20.5069348 | 1.59070703 | 0.4452029 | 3.00E-05 | 2.10E-04 | Afap1l1 |
| ENSMUSG00( | 97.4147424 | 1.58107023 | 0.26624674 | 2.94E-10 | 5.04E-09 | Dcp1b |
| ENSMUSG00( | 7.61651893 | 1.57831515 | 0.65486645 | 9.47E-04 | 0.00462388 | Pappa2 |
| ENSMUSG00( | 101.598172 | 1.56873266 | 0.26315444 | 2.18E-10 | 3.81E-09 | Tmem241 |
| ENSMUSG00( | 1067.94032 | 1.56696156 | 0.18338603 | 1.04E-18 | 5.52E-17 | Nacc2 |
| ENSMUSG00( | 47.532219 | 1.56583809 | 0.40555569 | 8.90E-06 | 6.96E-05 | Hsf4 |
| ENSMUSG00( | 632.968651 | 1.56544922 | 0.18393378 | 1.44E-18 | 7.55E-17 | Abcc3 |
| ENSMUSG00( | 28.465981 | 1.56491586 | 0.32051424 | 9.75E-08 | 1.10E-06 | Gm19434 |
| ENSMUSG00( | 18.9822788 | 1.56364648 | 0.56389905 | 3.33E-04 | 0.00181397 | Clec4a1 |
| ENSMUSG00( | 471.698151 | 1.56354527 | 0.1119209 | 2.04E-45 | 8.17E-43 | Ttc21b |
| ENSMUSG00( | 65.6207307 | 1.55530978 | 0.37068801 | 2.10E-06 | 1.88E-05 | Gm19253 |
| ENSMUSG00( | 228.983498 | 1.55154437 | 0.21182942 | 1.91E-14 | 6.10E-13 | Utrn |
| ENSMUSG00( | 11.6605385 | 1.5475904 | 0.83471587 | 0.0028233 | 0.01190254 | Spdye4b |
| ENSMUSG00( | 204.659506 | 1.54083882 | 0.17065769 | 1.57E-20 | 1.04E-18 | Ilvbl |
| ENSMUSG00( | 1137.3713 | 1.5397939 | 0.17168723 | 2.06E-20 | 1.35E-18 | Gas7 |
| ENSMUSG00( | 69.9039702 | 1.53174621 | 0.37604162 | 4.70E-06 | 3.92E-05 | Nlrp1b |
| ENSMUSG00( | 29.9811075 | 1.5290315 | 0.40821829 | 1.54E-05 | 1.15E-04 | Btbd17 |
| ENSMUSG00( | 2137.52978 | 1.52899035 | 0.10080333 | 4.02E-53 | 2.51E-50 | Esyt1 |
| ENSMUSG00( | 132.588979 | 1.51936263 | 0.2044044 | 9.65E-15 | 3.18E-13 | Clec4a3 |
| ENSMUSG00( | 204.79411 | 1.51873572 | 0.31349191 | 9.38E-08 | 1.06E-06 | Aldh1l2 |
| ENSMUSG00( | 145.144176 | 1.51720099 | 0.30786482 | 6.75E-08 | 7.95E-07 | Il1rl1 |
| ENSMUSG00( | 1584.24367 | 1.51357274 | 0.59764572 | 5.72E-04 | 0.00293614 | Fos |
| ENSMUSG00( | 56.0983505 | 1.5059972 | 0.25430876 | 2.80E-10 | 4.80E-09 | Nod1 |
| ENSMUSG00( | 6.81434871 | 1.50551837 | 1.43702338 | 0.0115221 | 0.03918555 | Gm48734 |
| ENSMUSG00( | 132.493196 | 1.50240402 | 0.34057489 | 6.07E-07 | 5.97E-06 | St18 |
| ENSMUSG00( | 15.9982904 | 1.50061229 | 0.56505133 | 5.60E-04 | 0.00288401 | Gm49654 |
| ENSMUSG00( | 133.102258 | 1.49627915 | 0.29755936 | 3.90E-08 | 4.77E-07 | Lrrc20 |

| | | | | | | |
|---|---|---|---|---|---|---|
| ENSMUSG00( | 1997.5767 | 1.48686552 | 0.09638717 | 9.85E-55 | 6.59E-52 | Abcc8 |
| ENSMUSG00( | 7.17998975 | 1.48505512 | 0.84976313 | 0.00512821 | 0.01985486 | Smarca5-ps |
| ENSMUSG00( | 10.9359401 | 1.48266017 | 0.65026787 | 0.00136075 | 0.00631094 | Spic |
| ENSMUSG00( | 669.24881 | 1.48150456 | 0.16733609 | 7.36E-20 | 4.51E-18 | Dglucy |
| ENSMUSG00( | 1496.99483 | 1.4738412 | 0.10665445 | 1.50E-44 | 5.41E-42 | Slc6a12 |
| ENSMUSG00( | 55.5852884 | 1.4621681 | 0.30574036 | 1.52E-07 | 1.65E-06 | Rpl3l |
| ENSMUSG00( | 90.9123084 | 1.45982937 | 0.31280563 | 2.41E-07 | 2.53E-06 | Siglece |
| ENSMUSG00( | 8.35817159 | 1.45718319 | 0.97234685 | 0.00657209 | 0.02449488 | Mfsd6l |
| ENSMUSG00( | 161.822368 | 1.45696017 | 0.23312667 | 3.57E-11 | 7.05E-10 | Trpm2 |
| ENSMUSG00( | 773.440009 | 1.45370811 | 0.17259101 | 3.12E-18 | 1.59E-16 | Cd302 |
| ENSMUSG00( | 121.724412 | 1.44591992 | 0.24363197 | 2.61E-10 | 4.50E-09 | Inka1 |
| ENSMUSG00( | 37.1622668 | 1.44401151 | 0.44787353 | 1.02E-04 | 6.34E-04 | Fbxo10 |
| ENSMUSG00( | 47.5655589 | 1.4390515 | 0.33785099 | 1.86E-06 | 1.67E-05 | Tmod1 |
| ENSMUSG00( | 77.082734 | 1.43154321 | 0.29253193 | 9.60E-08 | 1.09E-06 | Lrfn1 |
| ENSMUSG00( | 215.787911 | 1.42901927 | 0.16274847 | 1.49E-19 | 8.65E-18 | Gna15 |
| ENSMUSG00( | 197.830339 | 1.42726571 | 0.22922571 | 4.10E-11 | 8.00E-10 | Tsku |
| ENSMUSG00( | 96.2277337 | 1.42682524 | 0.33116073 | 1.33E-06 | 1.23E-05 | Slc46a3 |
| ENSMUSG00( | 247.986903 | 1.42237155 | 0.28025703 | 3.23E-08 | 4.01E-07 | Gjb3 |
| ENSMUSG00( | 181.613064 | 1.41868421 | 0.34123717 | 2.41E-06 | 2.13E-05 | Cysltr1 |
| ENSMUSG00( | 16.7153224 | 1.41431881 | 0.65469327 | 0.00186128 | 0.00826734 | Gm14549 |
| ENSMUSG00( | 35.3174561 | 1.41326234 | 0.45520214 | 1.66E-04 | 9.77E-04 | Casp7 |
| ENSMUSG00( | 843.40183 | 1.40722564 | 0.09971062 | 1.96E-46 | 8.25E-44 | Rpap1 |
| ENSMUSG00( | 11.9984382 | 1.40065363 | 0.86586786 | 0.00587547 | 0.02228927 | Gm48239 |
| ENSMUSG00( | 8.73273127 | 1.39782167 | 0.8312911 | 0.00553723 | 0.02121548 | 4732496C06I |
| ENSMUSG00( | 1159.03969 | 1.39705667 | 0.23867541 | 3.00E-10 | 5.14E-09 | H1f2 |
| ENSMUSG00( | 1281.36487 | 1.39700387 | 0.09634845 | 4.89E-49 | 2.33E-46 | Idh1 |
| ENSMUSG00( | 227.647474 | 1.39581548 | 0.17373267 | 1.02E-16 | 4.33E-15 | Map4k2 |
| ENSMUSG00( | 9.44123779 | 1.39542869 | 1.49445645 | 0.01080679 | 0.03720805 | Asb4 |
| ENSMUSG00( | 94.5478712 | 1.39324867 | 0.33929796 | 3.59E-06 | 3.08E-05 | Klhl33 |
| ENSMUSG00( | 712.984537 | 1.39322882 | 0.09524392 | 1.90E-49 | 9.27E-47 | Daglb |
| ENSMUSG00( | 450.377872 | 1.3927444 | 0.14315385 | 2.20E-23 | 1.93E-21 | Plcg1 |
| ENSMUSG00( | 107.415059 | 1.39239161 | 0.22796354 | 9.35E-11 | 1.72E-09 | Igf2bp3 |
| ENSMUSG00( | 563.006763 | 1.39224582 | 0.16865529 | 1.39E-17 | 6.48E-16 | Dennd1a |
| ENSMUSG00( | 78.9221452 | 1.39196254 | 0.41289988 | 5.24E-05 | 3.48E-04 | Aif1 |
| ENSMUSG00( | 161.859937 | 1.3891327 | 0.19233101 | 5.11E-14 | 1.51E-12 | Runx2 |
| ENSMUSG00( | 161.005806 | 1.38177066 | 0.18705197 | 1.39E-14 | 4.54E-13 | Bvht |
| ENSMUSG00( | 8.87146715 | 1.378836 | 0.84081115 | 0.00501245 | 0.01946486 | Fam89a |
| ENSMUSG00( | 3816.26623 | 1.37820393 | 0.11348036 | 5.63E-35 | 1.23E-32 | Ptpn6 |
| ENSMUSG00( | 13.267977 | 1.37721574 | 0.75347983 | 0.00384528 | 0.01554718 | Tmem74 |
| ENSMUSG00( | 1352.11552 | 1.37445571 | 0.13300272 | 3.40E-26 | 3.73E-24 | Jag2 |
| ENSMUSG00( | 100.680943 | 1.37368451 | 0.42457623 | 8.66E-05 | 5.48E-04 | Ccdc18 |

| | | | | | | |
|---|---|---|---|---|---|---|
| ENSMUSG00( | 92.0268932 | 1.37276081 | 0.29311037 | 2.51E-07 | 2.62E-06 | Mpv17l |
| ENSMUSG00( | 144.002713 | 1.37091896 | 0.17633928 | 8.04E-16 | 3.05E-14 | Sfxn5 |
| ENSMUSG00( | 34.6594165 | 1.3693948 | 0.31297949 | 1.07E-06 | 1.01E-05 | Ttc21a |
| ENSMUSG00( | 312.411054 | 1.36795427 | 0.14575357 | 6.94E-22 | 5.18E-20 | Ctdspl |
| ENSMUSG00( | 7.70774026 | 1.36678219 | 1.14906566 | 0.01388151 | 0.04570426 | 2900052L18R |
| ENSMUSG00( | 537.044676 | 1.36262846 | 0.28104071 | 1.01E-07 | 1.14E-06 | Akap6 |
| ENSMUSG00( | 1312.60023 | 1.3613251 | 0.08212984 | 1.06E-62 | 8.72E-60 | Jade1 |
| ENSMUSG00( | 30.8161867 | 1.35710418 | 0.64707066 | 0.00310871 | 0.01294569 | Abca6 |
| ENSMUSG00( | 29.9538957 | 1.35195905 | 0.39514725 | 5.66E-05 | 3.73E-04 | |
| ENSMUSG00( | 574.624935 | 1.34966199 | 0.11812533 | 2.92E-31 | 4.84E-29 | Hadh |
| ENSMUSG00( | 445.290483 | 1.34939539 | 0.14844393 | 9.55E-21 | 6.47E-19 | Entpd6 |
| ENSMUSG00( | 53.7494897 | 1.34885401 | 0.56573045 | 0.00101821 | 0.00492059 | Zfyve28 |
| ENSMUSG00( | 25.8122041 | 1.34640798 | 0.37348153 | 2.57E-05 | 1.83E-04 | Ripor3 |
| ENSMUSG00( | 79.8845852 | 1.34044732 | 0.30039276 | 6.66E-07 | 6.51E-06 | Calr3 |
| ENSMUSG00( | 163.29215 | 1.33779451 | 0.17843214 | 7.00E-15 | 2.37E-13 | Mpp7 |
| ENSMUSG00( | 65.9557394 | 1.33696396 | 0.33637302 | 6.00E-06 | 4.89E-05 | Ccdc9b |
| ENSMUSG00( | 4744.499 | 1.33695357 | 0.06604138 | 3.85E-92 | 5.79E-89 | Nceh1 |
| ENSMUSG00( | 138.79403 | 1.3348763 | 0.21062651 | 2.41E-11 | 4.86E-10 | Tcam1 |
| ENSMUSG00( | 34.7111846 | 1.33452062 | 0.40627737 | 9.01E-05 | 5.67E-04 | Lmtk3 |
| ENSMUSG00( | 303.444715 | 1.33389342 | 0.15841695 | 3.85E-18 | 1.92E-16 | Prkce |
| ENSMUSG00( | 379.76369 | 1.33239124 | 0.11578405 | 1.29E-31 | 2.26E-29 | Limk1 |
| ENSMUSG00( | 341.096025 | 1.33111049 | 0.13491487 | 5.75E-24 | 5.36E-22 | Sall3 |
| ENSMUSG00( | 1904.79492 | 1.33001 | 0.10288108 | 3.00E-39 | 9.18E-37 | Myo1f |
| ENSMUSG00( | 426.84451 | 1.32382526 | 0.14167297 | 9.25E-22 | 6.85E-20 | Add3 |
| ENSMUSG00( | 63.9469297 | 1.3197366 | 0.25039358 | 1.39E-08 | 1.82E-07 | Ogdhl |
| ENSMUSG00( | 215.403038 | 1.31789215 | 0.15316175 | 7.74E-19 | 4.17E-17 | Etfrf1 |
| ENSMUSG00( | 279.468358 | 1.31745838 | 0.13319192 | 4.71E-24 | 4.46E-22 | Nectin2 |
| ENSMUSG00( | 12.8959871 | 1.31569031 | 0.53448348 | 0.00100319 | 0.00485866 | Gm44154 |
| ENSMUSG00( | 16.5022855 | 1.31174664 | 0.60611835 | 0.00218523 | 0.00954664 | Zfp963 |
| ENSMUSG00( | 413.568351 | 1.31046351 | 0.15560232 | 3.58E-18 | 1.79E-16 | Nr1h3 |
| ENSMUSG00( | 868.051464 | 1.30632298 | 0.13735496 | 1.83E-22 | 1.42E-20 | Kif21b |
| ENSMUSG00( | 5.59347868 | 1.30332841 | 1.09507555 | 0.01244475 | 0.04171878 | F630028O10R |
| ENSMUSG00( | 485.847959 | 1.300231 | 0.14894316 | 2.44E-19 | 1.36E-17 | Fcgr2b |
| ENSMUSG00( | 66.832585 | 1.29911199 | 0.27419764 | 2.23E-07 | 2.35E-06 | Gm13421 |
| ENSMUSG00( | 72.6996042 | 1.2984821 | 0.25723924 | 4.69E-08 | 5.68E-07 | Ttc39aos1 |
| ENSMUSG00( | 13.6962076 | 1.29781038 | 0.46062233 | 3.91E-04 | 0.00209077 | Slc7a14 |
| ENSMUSG00( | 552.857929 | 1.29574397 | 0.20580866 | 2.74E-11 | 5.49E-10 | Abca1 |
| ENSMUSG00( | 235.575459 | 1.29514941 | 0.23652315 | 3.88E-09 | 5.62E-08 | Ralgps1 |
| ENSMUSG00( | 40.4925647 | 1.29327593 | 0.32698519 | 7.47E-06 | 5.95E-05 | Sgcb |
| ENSMUSG00( | 153.930895 | 1.29321792 | 0.18611648 | 4.48E-13 | 1.15E-11 | Cd300lg |
| ENSMUSG00( | 81.1198595 | 1.29238886 | 0.28837401 | 7.56E-07 | 7.31E-06 | Abi3 |

| | | | | | | |
|---|---|---|---|---|---|---|
| ENSMUSG00( | 239.977935 | 1.29184229 | 0.17587812 | 2.54E-14 | 7.92E-13 | Kifc3 |
| ENSMUSG00( | 534.214914 | 1.28759534 | 0.12303767 | 1.23E-26 | 1.40E-24 | Nfil3 |
| ENSMUSG00( | 44.9730154 | 1.28442647 | 0.33344567 | 1.17E-05 | 8.94E-05 | Wasf1 |
| ENSMUSG00( | 118.660049 | 1.28437851 | 0.36506963 | 3.49E-05 | 2.41E-04 | Cirbp |
| ENSMUSG00( | 161.762341 | 1.2837547 | 0.23082032 | 2.43E-09 | 3.66E-08 | Arhgef19 |
| ENSMUSG00( | 188.97801 | 1.28201806 | 0.28172524 | 4.57E-07 | 4.58E-06 | |
| ENSMUSG00( | 314.915161 | 1.27996926 | 0.16717004 | 1.89E-15 | 6.81E-14 | Map3k5 |
| ENSMUSG00( | 8.96568635 | 1.27291145 | 1.0217617 | 0.00767625 | 0.02789105 | Igsf3 |
| ENSMUSG00( | 69.5815331 | 1.26889217 | 0.33674643 | 1.62E-05 | 1.20E-04 | Zbtb16 |
| ENSMUSG00( | 48.9236978 | 1.26540844 | 0.34283051 | 2.01E-05 | 1.46E-04 | Wipi1 |
| ENSMUSG00( | 33.2784327 | 1.26437054 | 0.42775295 | 2.69E-04 | 0.00150214 | H1f3 |
| ENSMUSG00( | 390.979124 | 1.26184095 | 0.1298231 | 2.61E-23 | 2.26E-21 | Ninl |
| ENSMUSG00( | 16.8836452 | 1.26027498 | 0.65584508 | 0.00343785 | 0.0140954 | Zc3h6 |
| ENSMUSG00( | 2233.48384 | 1.2596216 | 0.05324624 | 1.52E-124 | 9.18E-121 | Xbp1 |
| ENSMUSG00( | 669.641615 | 1.258091 | 0.1230754 | 1.79E-25 | 1.88E-23 | Haus4 |
| ENSMUSG00( | 5.01663027 | 1.2579775 | 0.95095256 | 0.00839988 | 0.03013249 | Gm16565 |
| ENSMUSG00( | 11.8461408 | 1.25510092 | 0.55492774 | 0.00191962 | 0.0085014 | Fcrlb |
| ENSMUSG00( | 594.369342 | 1.25312154 | 0.12272226 | 2.00E-25 | 2.08E-23 | Pde3b |
| ENSMUSG00( | 247.376904 | 1.25271874 | 0.17734081 | 1.67E-13 | 4.59E-12 | Nt5dc1 |
| ENSMUSG00( | 16.7112159 | 1.25215287 | 0.46685846 | 5.49E-04 | 0.00283493 | Sstr5 |
| ENSMUSG00( | 487.520377 | 1.25017653 | 0.22699605 | 3.75E-09 | 5.44E-08 | Abca9 |
| ENSMUSG00( | 147.60347 | 1.24865284 | 0.2302936 | 6.56E-09 | 9.10E-08 | Tbc1d8 |
| ENSMUSG00( | 37.5616759 | 1.24840366 | 0.39542544 | 1.38E-04 | 8.31E-04 | Hpse |
| ENSMUSG00( | 2593.73491 | 1.24574757 | 0.2268242 | 3.87E-09 | 5.60E-08 | H6pd |
| ENSMUSG00( | 361.043484 | 1.24232293 | 0.28393367 | 1.06E-06 | 1.00E-05 | Siglec1 |
| ENSMUSG00( | 15.3060995 | 1.24059363 | 0.51458227 | 0.00127193 | 0.00595865 | 9030407P20F |
| ENSMUSG00( | 174.732063 | 1.24042105 | 0.22274604 | 2.67E-09 | 3.99E-08 | Nmnat3 |
| ENSMUSG00( | 295.243493 | 1.23901821 | 0.15454994 | 1.18E-16 | 4.94E-15 | Lrrcc1 |
| ENSMUSG00( | 338.176252 | 1.23530758 | 0.16383112 | 5.00E-15 | 1.73E-13 | Nsun3 |
| ENSMUSG00( | 79.8799985 | 1.23384912 | 0.22638298 | 5.15E-09 | 7.28E-08 | |
| ENSMUSG00( | 67.7524845 | 1.23310141 | 0.43770982 | 3.57E-04 | 0.00192648 | Acss2 |
| ENSMUSG00( | 256.592951 | 1.23156752 | 0.1437705 | 1.10E-18 | 5.82E-17 | Ints6l |
| ENSMUSG00( | 20.2088543 | 1.22905245 | 0.87408402 | 0.01086291 | 0.03733726 | Gm2415 |
| ENSMUSG00( | 1055.10161 | 1.22795138 | 0.0960855 | 1.82E-38 | 5.30E-36 | Slc12a7 |
| ENSMUSG00( | 631.447743 | 1.22515273 | 0.25272674 | 1.13E-07 | 1.25E-06 | Plxdc1 |
| ENSMUSG00( | 28.8613382 | 1.22408472 | 0.44656106 | 5.50E-04 | 0.00283745 | Gm28417 |
| ENSMUSG00( | 146.135429 | 1.21942224 | 0.18901125 | 1.15E-11 | 2.42E-10 | Slc6a9 |
| ENSMUSG00( | 262.029313 | 1.2188654 | 0.16533284 | 2.08E-14 | 6.58E-13 | Abhd8 |
| ENSMUSG00( | 959.771817 | 1.21873708 | 0.11691525 | 1.94E-26 | 2.19E-24 | Arhgap19 |
| ENSMUSG00( | 60.5466372 | 1.21820125 | 0.30025543 | 5.22E-06 | 4.31E-05 | Selenbp1 |
| ENSMUSG00( | 102.102351 | 1.21722366 | 0.21543617 | 1.80E-09 | 2.77E-08 | Tmem102 |

| | | | | | | |
|---|---|---|---|---|---|---|
| ENSMUSG00( | 25.7156242 | 1.21650235 | 0.51548259 | 0.00128419 | 0.0060093 | Tspan17 |
| ENSMUSG00( | 273.835412 | 1.2152729 | 0.17762662 | 8.51E-13 | 2.10E-11 | Ift140 |
| ENSMUSG00( | 56.1197521 | 1.21513418 | 0.38635239 | 1.38E-04 | 8.33E-04 | Sbsn |
| ENSMUSG00( | 267.648798 | 1.21398329 | 0.18402778 | 4.43E-12 | 9.93E-11 | Ephx1 |
| ENSMUSG00( | 33.4745612 | 1.21295571 | 0.35331735 | 6.05E-05 | 3.96E-04 | Hcn3 |
| ENSMUSG00( | 429.279792 | 1.21047745 | 0.13812578 | 2.10E-19 | 1.19E-17 | Gngt2 |
| ENSMUSG00( | 12.5892347 | 1.20697885 | 0.57815863 | 0.00272429 | 0.01154977 | Il1r1 |
| ENSMUSG00( | 785.151718 | 1.2037492 | 0.17947274 | 2.05E-12 | 4.79E-11 | Lfng |
| ENSMUSG00( | 261.995206 | 1.19870928 | 0.14830922 | 7.07E-17 | 3.06E-15 | Tango6 |
| ENSMUSG00( | 91.9249591 | 1.19801452 | 0.3406645 | 4.07E-05 | 2.77E-04 | Nectin3 |
| ENSMUSG00( | 479.698511 | 1.19728138 | 0.17837693 | 2.02E-12 | 4.74E-11 | Cdk19 |
| ENSMUSG00( | 42.0563195 | 1.19534516 | 0.3026445 | 7.84E-06 | 6.22E-05 | Gm14040 |
| ENSMUSG00( | 753.871353 | 1.19068362 | 0.22473451 | 1.18E-08 | 1.57E-07 | Xylt2 |
| ENSMUSG00( | 37.9053108 | 1.1859393 | 0.37497786 | 1.48E-04 | 8.87E-04 | Arpin |
| ENSMUSG00( | 329.863666 | 1.18576488 | 0.19190476 | 7.33E-11 | 1.38E-09 | Zfp395 |
| ENSMUSG00( | 117.278183 | 1.18519738 | 0.30429946 | 1.03E-05 | 7.88E-05 | Tlr5 |
| ENSMUSG00( | 1076.35083 | 1.18387053 | 0.14217754 | 8.84E-18 | 4.16E-16 | Rcan1 |
| ENSMUSG00( | 378.361789 | 1.18377797 | 0.2022911 | 4.97E-10 | 8.20E-09 | Stard4 |
| ENSMUSG00( | 22.9161624 | 1.18200714 | 0.38621768 | 2.05E-04 | 0.00117767 | Eef1akmt3 |
| ENSMUSG00( | 121.655706 | 1.18184848 | 0.20797198 | 1.60E-09 | 2.49E-08 | Tbc1d32 |
| ENSMUSG00( | 432.010897 | 1.18042633 | 0.15786529 | 8.88E-15 | 2.95E-13 | Fgd2 |
| ENSMUSG00( | 498.685105 | 1.17912266 | 0.12916627 | 8.57E-21 | 5.85E-19 | Dmxl2 |
| ENSMUSG00( | 335.641085 | 1.17750874 | 0.11335903 | 3.17E-26 | 3.50E-24 | Ufl1 |
| ENSMUSG00( | 152.396088 | 1.17597094 | 0.18207208 | 1.15E-11 | 2.43E-10 | Ube2e2 |
| ENSMUSG00( | 337.011719 | 1.17562295 | 0.15610324 | 5.92E-15 | 2.04E-13 | Tfap4 |
| ENSMUSG00( | 462.601075 | 1.17542993 | 0.14297069 | 2.04E-17 | 9.35E-16 | Gpsm1 |
| ENSMUSG00( | 592.916291 | 1.17511536 | 0.12072633 | 2.47E-23 | 2.16E-21 | Fzd7 |
| ENSMUSG00( | 136.200108 | 1.17418208 | 0.27854332 | 2.31E-06 | 2.05E-05 | Mob3b |
| ENSMUSG00( | 15.9707884 | 1.17243708 | 0.55077856 | 0.00241771 | 0.01042615 | Atp10b |
| ENSMUSG00( | 206.033249 | 1.17192556 | 0.24619925 | 2.07E-07 | 2.19E-06 | Khk |
| ENSMUSG00( | 4401.90045 | 1.17098039 | 0.20060567 | 4.91E-10 | 8.12E-09 | Tlr2 |
| ENSMUSG00( | 1333.4455 | 1.17082558 | 0.12125449 | 5.08E-23 | 4.25E-21 | Sestd1 |
| ENSMUSG00( | 939.764108 | 1.16762853 | 0.16065933 | 4.25E-14 | 1.28E-12 | Mvb12b |
| ENSMUSG00( | 4531.4038 | 1.16639074 | 0.05815204 | 1.68E-90 | 2.33E-87 | Arhgef6 |
| ENSMUSG00( | 67.4622232 | 1.16624187 | 0.28720229 | 5.18E-06 | 4.28E-05 | Cradd |
| ENSMUSG00( | 1070.30909 | 1.16534494 | 0.10233905 | 1.41E-30 | 2.24E-28 | Irag2 |
| ENSMUSG00( | 373.685559 | 1.16392432 | 0.15749107 | 1.56E-14 | 5.03E-13 | Cep78 |
| ENSMUSG00( | 403.055251 | 1.16388438 | 0.14561445 | 1.46E-16 | 6.01E-15 | Mettl8 |
| ENSMUSG00( | 83.8935599 | 1.16319493 | 0.35778381 | 1.02E-04 | 6.34E-04 | Mfap3l |
| ENSMUSG00( | 16.4016042 | 1.16283511 | 0.54798119 | 0.0023832 | 0.01029947 | Fkbp14 |
| ENSMUSG00( | 40.5022263 | 1.16143418 | 0.36891406 | 1.74E-04 | 0.00102143 | Sort1 |

| | | | | | | |
|---|---|---|---|---|---|---|
| ENSMUSG00( | 1727.44496 | 1.16083502 | 0.23783897 | 1.04E-07 | 1.17E-06 | Sema4a |
| ENSMUSG00( | 21.6867475 | 1.15946388 | 0.5396529 | 0.00250704 | 0.01076378 | |
| ENSMUSG00( | 147.40364 | 1.15828499 | 0.3360692 | 5.08E-05 | 3.39E-04 | Slc6a13 |
| ENSMUSG00( | 9.20204414 | 1.15723963 | 1.12231937 | 0.01367928 | 0.04513712 | Itga8 |
| ENSMUSG00( | 670.501709 | 1.15564579 | 0.11775307 | 1.51E-23 | 1.34E-21 | Chst10 |
| ENSMUSG00( | 919.597752 | 1.15563504 | 0.31590052 | 2.29E-05 | 1.65E-04 | Plxnc1 |
| ENSMUSG00( | 3211.97688 | 1.15193165 | 0.39377036 | 2.80E-04 | 0.00155967 | Mafb |
| ENSMUSG00( | 163.126952 | 1.14973656 | 0.23237536 | 7.88E-08 | 9.09E-07 | Plekhg5 |
| ENSMUSG00( | 4.84273625 | 1.14714607 | 0.92608135 | 0.01010892 | 0.0352078 | Gm11541 |
| ENSMUSG00( | 597.968172 | 1.14680741 | 0.0927536 | 1.61E-35 | 3.60E-33 | Zfyve19 |
| ENSMUSG00( | 100.046039 | 1.14274356 | 0.23508015 | 1.28E-07 | 1.41E-06 | Nipa1 |
| ENSMUSG00( | 40.2235815 | 1.14245348 | 0.51651321 | 0.00193146 | 0.00854752 | F830045P16F |
| ENSMUSG00( | 14.2417668 | 1.14227353 | 0.86769595 | 0.0106805 | 0.03684818 | Bach2os |
| ENSMUSG00( | 1076.03389 | 1.13930363 | 0.13481348 | 3.21E-18 | 1.62E-16 | Kif13a |
| ENSMUSG00( | 9.38287822 | 1.13518518 | 0.60328812 | 0.00427914 | 0.01702703 | Tmtc2 |
| ENSMUSG00( | 154.263922 | 1.13502686 | 0.2523666 | 7.30E-07 | 7.08E-06 | Accs |
| ENSMUSG00( | 823.02468 | 1.13438373 | 0.16087341 | 1.97E-13 | 5.32E-12 | Atr |
| ENSMUSG00( | 353.564189 | 1.12852569 | 0.20250091 | 2.70E-09 | 4.02E-08 | Tmco4 |
| ENSMUSG00( | 109.157088 | 1.12806947 | 0.28303624 | 6.88E-06 | 5.54E-05 | Mtus1 |
| ENSMUSG00( | 53.8788303 | 1.12668318 | 0.37910732 | 2.99E-04 | 0.0016478 | Or5v1b |
| ENSMUSG00( | 77.6116668 | 1.12652638 | 0.28463217 | 7.93E-06 | 6.29E-05 | Wdpcp |
| ENSMUSG00( | 134.657111 | 1.12630281 | 0.27271501 | 3.68E-06 | 3.15E-05 | Eif4e3 |
| ENSMUSG00( | 192.719471 | 1.12615974 | 0.19216998 | 5.18E-10 | 8.52E-09 | Prkra |
| ENSMUSG00( | 15.0381179 | 1.12584954 | 0.63523784 | 0.00473961 | 0.01856897 | A530064D06F |
| ENSMUSG00( | 82.2300128 | 1.12519706 | 0.34241219 | 9.71E-05 | 6.07E-04 | Mutyh |
| ENSMUSG00( | 182.822575 | 1.12480182 | 0.17574279 | 1.75E-11 | 3.59E-10 | Golim4 |
| ENSMUSG00( | 121.380693 | 1.12263608 | 0.2998944 | 1.88E-05 | 1.38E-04 | H1f0 |
| ENSMUSG00( | 20.2770141 | 1.1223811 | 0.63768331 | 0.00474415 | 0.01858271 | Serpinb12 |
| ENSMUSG00( | 671.062236 | 1.12164688 | 0.1429478 | 4.85E-16 | 1.88E-14 | 6430548M08I |
| ENSMUSG00( | 251.640168 | 1.12144494 | 0.16870909 | 3.22E-12 | 7.33E-11 | Dram1 |
| ENSMUSG00( | 957.244017 | 1.12025093 | 0.08640404 | 2.39E-39 | 7.57E-37 | Rnf220 |
| ENSMUSG00( | 1482.4659 | 1.12000065 | 0.1127911 | 3.59E-24 | 3.45E-22 | Prmt7 |
| ENSMUSG00( | 293.087258 | 1.11939426 | 0.18920598 | 3.84E-10 | 6.48E-09 | Bcl9l |
| ENSMUSG00( | 15.095539 | 1.11722015 | 0.74424396 | 0.00988842 | 0.03455306 | Rgs18 |
| ENSMUSG00( | 290.457906 | 1.11138503 | 0.15820324 | 2.51E-13 | 6.70E-12 | Adarb1 |
| ENSMUSG00( | 190.133176 | 1.10993228 | 0.29880432 | 2.06E-05 | 1.50E-04 | Hlcs |
| ENSMUSG00( | 8.48308424 | 1.10954526 | 0.93147076 | 0.01209517 | 0.04075738 | Igf1os |
| ENSMUSG00( | 54.0466759 | 1.10830029 | 0.54706935 | 0.00299985 | 0.0125531 | Ttc28 |
| ENSMUSG00( | 376.161148 | 1.10674032 | 0.14000077 | 3.23E-16 | 1.28E-14 | Nagk |
| ENSMUSG00( | 15.5706268 | 1.10122838 | 0.70301664 | 0.00766801 | 0.02786671 | Gm26964 |
| ENSMUSG00( | 545.864166 | 1.09588466 | 0.12436392 | 1.56E-19 | 8.98E-18 | Abcd1 |

| | | | | | | |
|---|---|---|---|---|---|---|
| ENSMUSG00( | 512.01074 | 1.09539342 | 0.10957639 | 1.08E-24 | 1.07E-22 | Akr1c13 |
| ENSMUSG00( | 1203.43004 | 1.09438984 | 0.22306508 | 1.00E-07 | 1.13E-06 | Nav2 |
| ENSMUSG00( | 360.247505 | 1.09342595 | 0.16232005 | 2.09E-12 | 4.87E-11 | Tmem94 |
| ENSMUSG00( | 698.138046 | 1.09183682 | 0.12383796 | 1.40E-19 | 8.14E-18 | Slc9a8 |
| ENSMUSG00( | 315.902268 | 1.08996929 | 0.12848486 | 2.73E-18 | 1.42E-16 | Cchcr1 |
| ENSMUSG00( | 223.961155 | 1.08879241 | 0.19953362 | 6.03E-09 | 8.42E-08 | Neil3 |
| ENSMUSG00( | 5.13876838 | 1.08851539 | 0.75463391 | 0.00924927 | 0.03260976 | Gm40457 |
| ENSMUSG00( | 78.5208324 | 1.08689837 | 0.27983793 | 1.20E-05 | 9.15E-05 | Jaml |
| ENSMUSG00( | 34.6906073 | 1.08686946 | 6.66463049 | 0.00839668 | 0.030127 | Rn7sk |
| ENSMUSG00( | 1336.46898 | 1.08580819 | 0.10529063 | 7.87E-26 | 8.46E-24 | Fcgr1 |
| ENSMUSG00( | 600.9637 | 1.08493942 | 0.10014402 | 3.05E-28 | 4.21E-26 | Shfl |
| ENSMUSG00( | 37.416431 | 1.08472086 | 0.34571952 | 1.73E-04 | 0.00101727 | Txlnb |
| ENSMUSG00( | 58.9093555 | 1.08442839 | 0.78899878 | 0.00850819 | 0.03050287 | Ptpn14 |
| ENSMUSG00( | 39.1315672 | 1.08423168 | 0.39351234 | 5.53E-04 | 0.00284927 | Hdgfl3 |
| ENSMUSG00( | 2219.64412 | 1.0793418 | 0.17477319 | 7.59E-11 | 1.42E-09 | Eno3 |
| ENSMUSG00( | 123.024775 | 1.07901316 | 0.22005343 | 1.09E-07 | 1.21E-06 | Lrp11 |
| ENSMUSG00( | 638.790359 | 1.07808576 | 0.10782708 | 2.20E-24 | 2.13E-22 | Kcnk6 |
| ENSMUSG00( | 893.575742 | 1.07739665 | 0.17102129 | 3.40E-11 | 6.75E-10 | Neurl3 |
| ENSMUSG00( | 194.283029 | 1.0771329 | 0.27646504 | 1.04E-05 | 7.98E-05 | Mtfp1 |
| ENSMUSG00( | 39.9668666 | 1.0765475 | 0.57202134 | 0.00547264 | 0.02099917 | Rtn4rl1 |
| ENSMUSG00( | 39.4823188 | 1.07568236 | 0.32632236 | 1.02E-04 | 6.36E-04 | 2210406H18I |
| ENSMUSG00( | 71.07446 | 1.07501196 | 0.34820258 | 2.13E-04 | 0.00122367 | Igip |
| ENSMUSG00( | 378.883372 | 1.07232134 | 0.12101772 | 8.12E-20 | 4.94E-18 | Vrk2 |
| ENSMUSG00( | 1719.88455 | 1.07167626 | 0.18345929 | 5.99E-10 | 9.79E-09 | Nrp1 |
| ENSMUSG00( | 47.0891522 | 1.06931564 | 0.3056619 | 5.38E-05 | 3.57E-04 | Gm20300 |
| ENSMUSG00( | 51.5811043 | 1.06806154 | 0.3344878 | 1.50E-04 | 8.98E-04 | Atp9a |
| ENSMUSG00( | 119.157901 | 1.06747568 | 0.18238354 | 5.95E-10 | 9.74E-09 | Ggh |
| ENSMUSG00( | 853.483056 | 1.06687349 | 0.08628343 | 5.42E-36 | 1.26E-33 | Smarca2 |
| ENSMUSG00( | 97.0564165 | 1.06245017 | 0.201503 | 1.67E-08 | 2.16E-07 | Hoxa1 |
| ENSMUSG00( | 78.5787751 | 1.06197198 | 0.3609432 | 3.15E-04 | 0.00172749 | Gm26532 |
| ENSMUSG00( | 676.70891 | 1.06185343 | 0.19857175 | 1.03E-08 | 1.38E-07 | Tcf19 |
| ENSMUSG00( | 707.037922 | 1.06167716 | 0.09777603 | 2.47E-28 | 3.48E-26 | Steap3 |
| ENSMUSG00( | 847.602671 | 1.060702 | 0.0898568 | 4.74E-33 | 9.02E-31 | Trappc6b |
| ENSMUSG00( | 460.128767 | 1.05889038 | 0.10555791 | 1.38E-24 | 1.36E-22 | Maged2 |
| ENSMUSG00( | 667.745809 | 1.0586841 | 0.12978 | 4.29E-17 | 1.91E-15 | Igf1r |
| ENSMUSG00( | 51.4521261 | 1.05860457 | 0.4528443 | 0.00166838 | 0.00753082 | Rgs11 |
| ENSMUSG00( | 1443.98281 | 1.05696484 | 0.08519695 | 3.12E-36 | 7.52E-34 | Madd |
| ENSMUSG00( | 75.1642804 | 1.05572239 | 0.36076272 | 3.43E-04 | 0.0018614 | Ypel3 |
| ENSMUSG00( | 88.2897083 | 1.05523971 | 0.26795098 | 8.97E-06 | 7.00E-05 | Col18a1 |
| ENSMUSG00( | 22.0313282 | 1.05517432 | 0.4764718 | 0.00249175 | 0.01070969 | A330074K22F |
| ENSMUSG00( | 1459.52469 | 1.05414016 | 0.1872933 | 2.14E-09 | 3.25E-08 | Apobec1 |

| | | | | | | |
|---|---|---|---|---|---|---|
| ENSMUSG00( | 136.946123 | 1.05241412 | 0.25861152 | 5.32E-06 | 4.38E-05 | Fam78a |
| ENSMUSG00( | 453.174487 | 1.05179688 | 0.14381795 | 3.22E-14 | 9.91E-13 | Enox2 |
| ENSMUSG00( | 296.584543 | 1.04887428 | 0.19446357 | 8.79E-09 | 1.20E-07 | Xylb |
| ENSMUSG00( | 85.1093331 | 1.04092221 | 0.32830722 | 1.59E-04 | 9.39E-04 | Gm35315 |
| ENSMUSG00( | 9.61706699 | 1.04000858 | 0.67804605 | 0.00910926 | 0.03222301 | Gm45151 |
| ENSMUSG00( | 212.791487 | 1.03996782 | 0.20408957 | 4.37E-08 | 5.31E-07 | Rab3d |
| ENSMUSG00( | 325.782307 | 1.03930999 | 0.14665534 | 1.89E-13 | 5.13E-12 | Dhrs7 |
| ENSMUSG00( | 53.4092909 | 1.03781194 | 0.30994673 | 8.92E-05 | 5.63E-04 | Rtn2 |
| ENSMUSG00( | 310.729694 | 1.03706094 | 0.18250433 | 1.64E-09 | 2.53E-08 | Suv39h2 |
| ENSMUSG00( | 588.789757 | 1.03573157 | 0.1672842 | 7.76E-11 | 1.45E-09 | Capn5 |
| ENSMUSG00( | 80.2627982 | 1.03521414 | 0.41087381 | 0.00107047 | 0.00514022 | Thbs1 |
| ENSMUSG00( | 67.2954178 | 1.03230554 | 0.35190433 | 3.62E-04 | 0.00195158 | Mblac2 |
| ENSMUSG00( | 3693.44484 | 1.03086205 | 0.2042086 | 4.41E-08 | 5.35E-07 | Cat |
| ENSMUSG00( | 1216.7029 | 1.03067583 | 0.20431695 | 5.24E-08 | 6.29E-07 | Stab1 |
| ENSMUSG00( | 45.3577535 | 1.02996076 | 0.35524512 | 3.88E-04 | 0.00207761 | Plekha6 |
| ENSMUSG00( | 170.908707 | 1.02984225 | 0.23219412 | 9.82E-07 | 9.31E-06 | Spire2 |
| ENSMUSG00( | 255.12676 | 1.02920614 | 0.22264433 | 4.47E-07 | 4.49E-06 | Fut7 |
| ENSMUSG00( | 310.33313 | 1.02723529 | 0.1858078 | 3.97E-09 | 5.73E-08 | Klf11 |
| ENSMUSG00( | 1072.34777 | 1.02449765 | 0.10610541 | 6.08E-23 | 4.97E-21 | Timeless |
| ENSMUSG00( | 147.054264 | 1.023643 | 0.20740065 | 1.14E-07 | 1.26E-06 | Nfia |
| ENSMUSG00( | 81.4395568 | 1.02320153 | 0.23788718 | 2.09E-06 | 1.87E-05 | Nadsyn1 |
| ENSMUSG00( | 107.731314 | 1.02012679 | 0.27498971 | 2.45E-05 | 1.75E-04 | Arhgap15 |
| ENSMUSG00( | 282.274858 | 1.01911166 | 0.13420313 | 4.26E-15 | 1.50E-13 | Septin10 |
| ENSMUSG00( | 118.644536 | 1.0187688 | 0.20041089 | 4.61E-08 | 5.59E-07 | Samd4 |
| ENSMUSG00( | 594.968392 | 1.01584517 | 0.21138531 | 1.84E-07 | 1.97E-06 | Slc30a1 |
| ENSMUSG00( | 637.071662 | 1.0154427 | 0.12159229 | 9.08E-18 | 4.26E-16 | Fbh1 |
| ENSMUSG00( | 462.723792 | 1.01462864 | 0.14595232 | 4.85E-13 | 1.24E-11 | Pomt1 |
| ENSMUSG00( | 100.912924 | 1.01332551 | 0.24600148 | 5.16E-06 | 4.27E-05 | Tamalin |
| ENSMUSG00( | 19.6606447 | 1.01301759 | 0.47855944 | 0.00326981 | 0.01351066 | Tst |
| ENSMUSG00( | 24.957675 | 1.01276849 | 0.42615631 | 0.00183301 | 0.00816383 | Cd244a |
| ENSMUSG00( | 3194.56528 | 1.01204427 | 0.17453537 | 6.02E-10 | 9.84E-09 | Slc16a10 |
| ENSMUSG00( | 128.787488 | 1.01010504 | 0.23794354 | 2.32E-06 | 2.06E-05 | Kyat3 |
| ENSMUSG00( | 376.786939 | 1.00880375 | 0.13669254 | 1.78E-14 | 5.70E-13 | Lrsam1 |
| ENSMUSG00( | 866.409471 | 1.00812909 | 0.13396055 | 7.16E-15 | 2.42E-13 | Tcf3 |
| ENSMUSG00( | 32.8065983 | 1.00736025 | 0.40595538 | 0.00139479 | 0.0064506 | Gm9920 |
| ENSMUSG00( | 806.864618 | 1.00708351 | 0.11412732 | 1.44E-19 | 8.38E-18 | Sfmbt2 |
| ENSMUSG00( | 14.6296796 | 1.00559437 | 0.62485081 | 0.00908747 | 0.03215854 | Fads6 |
| ENSMUSG00( | 3848.54886 | 1.00549764 | 0.17823236 | 2.17E-09 | 3.29E-08 | Rassf4 |
| ENSMUSG00( | 22.7979192 | 1.00533238 | 0.59771901 | 0.00755704 | 0.02751322 | Cand2 |
| ENSMUSG00( | 2036.84836 | 1.00467538 | 0.14812009 | 1.73E-12 | 4.09E-11 | Dnajc3 |
| ENSMUSG00( | 56.3989598 | 1.00431167 | 0.35440687 | 5.25E-04 | 0.00272544 | Ica1 |

| | | | | | | |
|---|---|---|---|---|---|---|
| ENSMUSG00( | 430.474922 | 1.00382064 | 0.13966116 | 8.96E-14 | 2.57E-12 | Pctp |
| ENSMUSG00( | 5.88425796 | 1.00293366 | 0.71347538 | 0.01047679 | 0.03630012 | Fam171b |
| ENSMUSG00( | 428.276326 | -1.0007162 | 0.09193961 | 1.75E-28 | 2.52E-26 | Cdc42ep4 |
| ENSMUSG00( | 68.1762209 | -1.001847 | 0.26568047 | 1.76E-05 | 1.30E-04 | |
| ENSMUSG00( | 584.796481 | -1.0036849 | 0.1849966 | 8.09E-09 | 1.10E-07 | Cd300lf |
| ENSMUSG00( | 33.6840578 | -1.0065941 | 0.29975102 | 8.68E-05 | 5.49E-04 | E230013L22R |
| ENSMUSG00( | 36.1473069 | -1.0100419 | 0.35730849 | 4.78E-04 | 0.00250745 | Gm15853 |
| ENSMUSG00( | 40.6868215 | -1.0100441 | 0.41793391 | 0.001385 | 0.00641519 | Pomc |
| ENSMUSG00( | 133.598593 | -1.0102483 | 0.21460236 | 3.00E-07 | 3.11E-06 | Zan |
| ENSMUSG00( | 7.52047194 | -1.0105386 | 1.17117977 | 0.01179544 | 0.03998722 | mt-Te |
| ENSMUSG00( | 199.11925 | -1.0114024 | 0.23665039 | 2.27E-06 | 2.01E-05 | Slc39a2 |
| ENSMUSG00( | 18.6513418 | -1.0125415 | 0.72828141 | 0.01110209 | 0.03801472 | Il1r2 |
| ENSMUSG00( | 20.8673427 | -1.0144543 | 0.42507896 | 0.00162398 | 0.00735617 | 6530409C15R |
| ENSMUSG00( | 579.75289 | -1.017316 | 0.23822043 | 2.19E-06 | 1.95E-05 | Parp14 |
| ENSMUSG00( | 10.4181382 | -1.01756 | 0.53059647 | 0.00419123 | 0.0167473 | Gm46637 |
| ENSMUSG00( | 6.58582546 | -1.0178711 | 0.81870784 | 0.01080306 | 0.0372023 | Gm12089 |
| ENSMUSG00( | 157.396644 | -1.0185572 | 0.2187151 | 3.84E-07 | 3.90E-06 | Card14 |
| ENSMUSG00( | 8.99418463 | -1.0186846 | 0.68698143 | 0.0087278 | 0.03114805 | 9530056E24R |
| ENSMUSG00( | 5.50452619 | -1.0189345 | 1.05862284 | 0.01417493 | 0.04645783 | Gm38365 |
| ENSMUSG00( | 5.92678264 | -1.0193259 | 0.82515092 | 0.01200967 | 0.04056124 | Gm44567 |
| ENSMUSG00( | 25.7397634 | -1.0206368 | 0.37649277 | 6.48E-04 | 0.00328117 | Tnfsf14 |
| ENSMUSG00( | 54.8668875 | -1.021127 | 0.21948203 | 3.79E-07 | 3.85E-06 | Ptgs2os |
| ENSMUSG00( | 665.092145 | -1.0257352 | 0.16439105 | 5.58E-11 | 1.07E-09 | Gcnt1 |
| ENSMUSG00( | 15.5267902 | -1.0266229 | 0.48497891 | 0.00302491 | 0.01263455 | 9330161L09R |
| ENSMUSG00( | 1178.49581 | -1.0268115 | 0.08623336 | 1.10E-33 | 2.23E-31 | Osgin2 |
| ENSMUSG00( | 461.594294 | -1.0274401 | 0.1562551 | 5.78E-12 | 1.27E-10 | Bcl2a1b |
| ENSMUSG00( | 35.1256599 | -1.0282146 | 0.31422006 | 1.10E-04 | 6.81E-04 | Smarcd3 |
| ENSMUSG00( | 14.7787249 | -1.0287222 | 0.4792995 | 0.00254239 | 0.01088587 | |
| ENSMUSG00( | 7.65583488 | -1.0290487 | 0.66167202 | 0.00744522 | 0.02718279 | Gm29367 |
| ENSMUSG00( | 8.42280861 | -1.029925 | 0.73030073 | 0.00941123 | 0.03310326 | Klhl35 |
| ENSMUSG00( | 25.9481522 | -1.0303021 | 0.428048 | 0.00140563 | 0.00649573 | Tcim |
| ENSMUSG00( | 12.1577052 | -1.0323453 | 0.55970205 | 0.00464968 | 0.01825224 | D130007C19I |
| ENSMUSG00( | 1447.96875 | -1.0326194 | 0.06904648 | 2.08E-51 | 1.10E-48 | Rsbn1 |
| ENSMUSG00( | 702.906546 | -1.0344648 | 0.10102105 | 1.27E-25 | 1.35E-23 | Vps37b |
| ENSMUSG00( | 4.68380441 | -1.0356001 | 1.34585116 | 0.01352893 | 0.04478005 | Gm47826 |
| ENSMUSG00( | 7.91238107 | -1.0356919 | 0.79624263 | 0.01102049 | 0.03777825 | Lag3 |
| ENSMUSG00( | 1175.7982 | -1.0383488 | 0.09531898 | 1.67E-28 | 2.43E-26 | Ndst1 |
| ENSMUSG00( | 24.0681709 | -1.0407081 | 0.46082043 | 0.00194312 | 0.00859493 | 4932441J04R |
| ENSMUSG00( | 328.138793 | -1.0424948 | 0.10234029 | 2.64E-25 | 2.71E-23 | Sema4b |
| ENSMUSG00( | 35.7043992 | -1.0429624 | 0.29564228 | 4.75E-05 | 3.19E-04 | Gm47813 |
| ENSMUSG00( | 803.358811 | -1.0446653 | 0.20955715 | 6.75E-08 | 7.95E-07 | Oas3 |

| | | | | | | |
|---|---|---|---|---|---|---|
| ENSMUSG00( | 58.5830471 | -1.04689 | 0.25460906 | 4.38E-06 | 3.67E-05 | |
| ENSMUSG00( | 427.962356 | -1.0475018 | 0.15080056 | 4.65E-13 | 1.19E-11 | Rigi |
| ENSMUSG00( | 175.309173 | -1.0480859 | 0.17289725 | 1.59E-10 | 2.83E-09 | Nr1d1 |
| ENSMUSG00( | 3.94745162 | -1.0488059 | 1.11287372 | 0.01291402 | 0.0430126 | Lhx9 |
| ENSMUSG00( | 391.292623 | -1.05155 | 0.18685502 | 2.27E-09 | 3.42E-08 | Mndal |
| ENSMUSG00( | 19.8213455 | -1.0532358 | 0.43803614 | 0.00144602 | 0.00665854 | 9130024F11F |
| ENSMUSG00( | 505.906869 | -1.0582414 | 0.09476751 | 6.32E-30 | 9.68E-28 | Zfp318 |
| ENSMUSG00( | 34.2504697 | -1.0619571 | 0.27444515 | 1.17E-05 | 8.91E-05 | Il12rb1 |
| ENSMUSG00( | 12.1394757 | -1.0620825 | 0.60679114 | 0.00556831 | 0.02128941 | Gm15787 |
| ENSMUSG00( | 44.2490182 | -1.0651847 | 0.29629573 | 3.37E-05 | 2.33E-04 | Gm17709 |
| ENSMUSG00( | 619.363438 | -1.0662431 | 0.13042482 | 3.44E-17 | 1.53E-15 | Tlr9 |
| ENSMUSG00( | 28.7029987 | -1.0680484 | 0.31150349 | 5.48E-05 | 3.62E-04 | Gm17435 |
| ENSMUSG00( | 21.5834506 | -1.0682443 | 0.37641099 | 4.57E-04 | 0.00240628 | Map3k13 |
| ENSMUSG00( | 2068.47208 | -1.0692949 | 0.1488047 | 6.16E-14 | 1.79E-12 | Lyst |
| ENSMUSG00( | 22.7228458 | -1.0699714 | 0.33724529 | 1.51E-04 | 8.99E-04 | Gm9888 |
| ENSMUSG00( | 325.874331 | -1.0710319 | 0.17220971 | 5.71E-11 | 1.09E-09 | Chn2 |
| ENSMUSG00( | 2665.41073 | -1.0711336 | 0.14381764 | 7.64E-15 | 2.56E-13 | Arhgef7 |
| ENSMUSG00( | 58.698094 | -1.0729365 | 0.33251489 | 1.22E-04 | 7.46E-04 | Ankrd37 |
| ENSMUSG00( | 108.772447 | -1.0730101 | 0.15807335 | 1.30E-12 | 3.10E-11 | Irgm2 |
| ENSMUSG00( | 3816.98703 | -1.073476 | 0.08059174 | 2.12E-41 | 7.37E-39 | Ccdc88a |
| ENSMUSG00( | 83.2936686 | -1.0740651 | 0.2133111 | 5.56E-08 | 6.66E-07 | Zfp113 |
| ENSMUSG00( | 408.490509 | -1.0764203 | 0.19114434 | 1.99E-09 | 3.04E-08 | Cd274 |
| ENSMUSG00( | 637.967513 | -1.0771431 | 0.16928331 | 2.53E-11 | 5.10E-10 | Tuba1a |
| ENSMUSG00( | 41.7329307 | -1.0788878 | 0.28285106 | 1.46E-05 | 1.09E-04 | Leng9 |
| ENSMUSG00( | 12.9539175 | -1.0848899 | 0.85610198 | 0.01055764 | 0.03649466 | Gpr31c |
| ENSMUSG00( | 21.2887659 | -1.0869272 | 0.42321908 | 9.05E-04 | 0.0044436 | Gabbr1 |
| ENSMUSG00( | 2990.18826 | -1.089734 | 0.22196148 | 9.53E-08 | 1.08E-06 | Ubc |
| ENSMUSG00( | 54.4934509 | -1.0913247 | 0.23853455 | 5.21E-07 | 5.17E-06 | Gm11696 |
| ENSMUSG00( | 566.22943 | -1.0968512 | 0.1394568 | 4.16E-16 | 1.63E-14 | Mllt11 |
| ENSMUSG00( | 17.4891779 | -1.0998039 | 0.40998833 | 6.65E-04 | 0.00336075 | Gm45515 |
| ENSMUSG00( | 265.723754 | -1.100202 | 0.10782465 | 1.99E-25 | 2.08E-23 | Rgs3 |
| ENSMUSG00( | 27.3254238 | -1.1019346 | 0.3249138 | 6.74E-05 | 4.37E-04 | 9130230N09F |
| ENSMUSG00( | 4.83962221 | -1.1073793 | 0.8261705 | 0.00927403 | 0.03267157 | Rapgef3 |
| ENSMUSG00( | 5842.22435 | -1.1080227 | 0.4334251 | 8.08E-04 | 0.00401362 | Plk2 |
| ENSMUSG00( | 5.3231822 | -1.1087923 | 0.8783255 | 0.0096403 | 0.03377751 | |
| ENSMUSG00( | 8.78018707 | -1.1091095 | 0.70655126 | 0.00698136 | 0.0257757 | Prr36 |
| ENSMUSG00( | 149.325805 | -1.1094603 | 0.28110972 | 7.79E-06 | 6.19E-05 | Epha2 |
| ENSMUSG00( | 273.7453 | -1.1097303 | 0.12648706 | 1.99E-19 | 1.14E-17 | Gm10052 |
| ENSMUSG00( | 29.9974692 | -1.112164 | 0.85383325 | 0.00755108 | 0.02750262 | Gm18445 |
| ENSMUSG00( | 40.2983985 | -1.1123768 | 0.27871069 | 6.90E-06 | 5.55E-05 | Asic1 |
| ENSMUSG00( | 21.265991 | -1.113502 | 0.41936579 | 6.99E-04 | 0.00351981 | Tfap2a |

| | | | | | | |
|---|---|---|---|---|---|---|
| ENSMUSG00( | 2013.05589 | -1.1147094 | 0.10974684 | 3.30E-25 | 3.35E-23 | Fem1b |
| ENSMUSG00( | 1037.87108 | -1.1148055 | 0.18669416 | 3.60E-10 | 6.11E-09 | Egln3 |
| ENSMUSG00( | 602.138139 | -1.1156984 | 0.13308893 | 6.25E-18 | 3.03E-16 | Slc5a3 |
| ENSMUSG00( | 540.630098 | -1.1160033 | 0.1204502 | 2.23E-21 | 1.61E-19 | Tbc1d9 |
| ENSMUSG00( | 1369.29376 | -1.1169545 | 0.17277186 | 9.36E-12 | 1.99E-10 | Tent5c |
| ENSMUSG00( | 17.8379753 | -1.1178978 | 0.4059743 | 5.05E-04 | 0.002632 | Gm41361 |
| ENSMUSG00( | 83.7034837 | -1.1186258 | 0.24655483 | 6.76E-07 | 6.59E-06 | P2ry1 |
| ENSMUSG00( | 96.1687049 | -1.1201576 | 0.19393879 | 8.03E-10 | 1.29E-08 | 1500015A07F |
| ENSMUSG00( | 6.34910826 | -1.12131 | 1.38360954 | 0.01403324 | 0.04607789 | Bex3 |
| ENSMUSG00( | 13.7874135 | -1.1236284 | 0.61175284 | 0.00435924 | 0.01728488 | Gm17147 |
| ENSMUSG00( | 157.032103 | -1.12501 | 0.21315936 | 1.44E-08 | 1.88E-07 | Jrk |
| ENSMUSG00( | 11.4733792 | -1.1254253 | 0.44704058 | 9.77E-04 | 0.00474446 | |
| ENSMUSG00( | 9.77556191 | -1.1269233 | 0.75453872 | 0.00720095 | 0.02645669 | Gm55066 |
| ENSMUSG00( | 158.818681 | -1.1280436 | 0.17584397 | 1.47E-11 | 3.07E-10 | Serac1 |
| ENSMUSG00( | 3.36931369 | -1.1280696 | 1.56926945 | 0.01437076 | 0.0469303 | Gm5828 |
| ENSMUSG00( | 18.793003 | -1.1287905 | 0.51666199 | 0.00213185 | 0.00933149 | Gm45477 |
| ENSMUSG00( | 3.35321498 | -1.1296705 | 1.99654683 | 0.01313894 | 0.04368123 | Lrrc18 |
| ENSMUSG00( | 10.4424336 | -1.1297587 | 0.55910567 | 0.00283151 | 0.01193436 | Gm50268 |
| ENSMUSG00( | 23.0084813 | -1.1355048 | 0.38449189 | 2.87E-04 | 0.00158927 | Gpd1 |
| ENSMUSG00( | 2139.46268 | -1.1400115 | 0.10750625 | 5.56E-27 | 6.48E-25 | Slc39a10 |
| ENSMUSG00( | 9.87582636 | -1.1412009 | 0.59369019 | 0.00353175 | 0.01442152 | 4930595D18F |
| ENSMUSG00( | 68.8861499 | -1.141641 | 0.21152125 | 7.16E-09 | 9.84E-08 | Hectd2 |
| ENSMUSG00( | 82.7980052 | -1.1423246 | 0.37806023 | 2.26E-04 | 0.00128507 | Etv4 |
| ENSMUSG00( | 344.783149 | -1.1435813 | 0.16422674 | 3.66E-13 | 9.56E-12 | Fat3 |
| ENSMUSG00( | 19.9128586 | -1.1437686 | 0.45378184 | 9.20E-04 | 0.00450835 | 5031434O11F |
| ENSMUSG00( | 1036.87252 | -1.1474611 | 0.21962278 | 1.84E-08 | 2.36E-07 | Gstp1 |
| ENSMUSG00( | 170.470487 | -1.1520724 | 0.21190912 | 5.45E-09 | 7.68E-08 | Arid3b |
| ENSMUSG00( | 3329.5694 | -1.1525536 | 0.28248827 | 4.10E-06 | 3.45E-05 | Tnf |
| ENSMUSG00( | 593.850205 | -1.1530726 | 0.17235147 | 2.64E-12 | 6.06E-11 | Gadd45b |
| ENSMUSG00( | 11.7791111 | -1.1544632 | 0.6209102 | 0.00381526 | 0.01544654 | Gm47246 |
| ENSMUSG00( | 45.6781133 | -1.1555351 | 0.37211729 | 1.62E-04 | 9.56E-04 | |
| ENSMUSG00( | 11.0600158 | -1.155698 | 0.58343656 | 0.00310342 | 0.01292947 | Armcx3 |
| ENSMUSG00( | 60.5170265 | -1.159406 | 0.23260073 | 6.06E-08 | 7.20E-07 | |
| ENSMUSG00( | 155.975958 | -1.1596542 | 0.17108472 | 1.25E-12 | 3.00E-11 | 5830432E09F |
| ENSMUSG00( | 11.1248372 | -1.1603888 | 0.60570357 | 0.00349017 | 0.01427754 | Gm17590 |
| ENSMUSG00( | 42.7099392 | -1.1609491 | 0.24138048 | 1.49E-07 | 1.62E-06 | 4833427F10F |
| ENSMUSG00( | 25.7737757 | -1.1648346 | 0.30164127 | 1.02E-05 | 7.87E-05 | Mid1-ps1 |
| ENSMUSG00( | 7.94886874 | -1.1650652 | 0.65908439 | 0.004987 | 0.01938684 | Gm42502 |
| ENSMUSG00( | 401.183883 | -1.1695483 | 0.17308004 | 1.40E-12 | 3.33E-11 | Aqp9 |
| ENSMUSG00( | 1677.08249 | -1.1726216 | 0.12443716 | 3.08E-22 | 2.37E-20 | Igf2r |
| ENSMUSG00( | 70.2741936 | -1.1747708 | 0.20576668 | 1.12E-09 | 1.77E-08 | Gm39469 |

| | | | | | | |
|---|---|---|---|---|---|---|
| ENSMUSG00( | 7.85482383 | -1.1766724 | 0.71506628 | 0.00540324 | 0.02075491 | Gm50080 |
| ENSMUSG00( | 8.47988092 | -1.1779179 | 0.62256754 | 0.00384175 | 0.01553637 | Ccdc17 |
| ENSMUSG00( | 3104.52151 | -1.1785009 | 0.2274127 | 2.09E-08 | 2.66E-07 | Cpeb4 |
| ENSMUSG00( | 12.0516668 | -1.1787694 | 0.48737878 | 0.00123066 | 0.00579535 | Gm15764 |
| ENSMUSG00( | 36.8643316 | -1.1791843 | 0.32378026 | 2.51E-05 | 1.79E-04 | A430057M04I |
| ENSMUSG00( | 13.5982855 | -1.183914 | 0.40424742 | 2.72E-04 | 0.00151827 | |
| ENSMUSG00( | 609.237783 | -1.1850987 | 0.1409807 | 4.67E-18 | 2.30E-16 | Ckb |
| ENSMUSG00( | 16.0101708 | -1.1878263 | 0.54289881 | 0.00204984 | 0.00901399 | 2510046G10F |
| ENSMUSG00( | 4.11185155 | -1.1881835 | 1.05069872 | 0.01073416 | 0.03700028 | 4930596I21Ri |
| ENSMUSG00( | 40.296223 | -1.1891374 | 0.29494666 | 5.43E-06 | 4.47E-05 | Ms4a6c |
| ENSMUSG00( | 15.4951461 | -1.1907139 | 0.45102327 | 6.89E-04 | 0.0034758 | Plcxd1 |
| ENSMUSG00( | 26.7596114 | -1.1943014 | 0.38167904 | 1.48E-04 | 8.86E-04 | 4933431K14F |
| ENSMUSG00( | 2403.19821 | -1.1986311 | 0.35754833 | 6.43E-05 | 4.19E-04 | Hilpda |
| ENSMUSG00( | 372.04963 | -1.1992929 | 0.25939212 | 4.23E-07 | 4.27E-06 | |
| ENSMUSG00( | 223.334885 | -1.201461 | 0.24271783 | 7.20E-08 | 8.43E-07 | Gatm |
| ENSMUSG00( | 132.607882 | -1.2038694 | 0.18459818 | 7.02E-12 | 1.52E-10 | Bcl2a1a |
| ENSMUSG00( | 376.174125 | -1.2056997 | 0.11629263 | 2.94E-26 | 3.26E-24 | Pdgfa |
| ENSMUSG00( | 17.7930452 | -1.2086182 | 0.41713222 | 2.96E-04 | 0.00163561 | Gm26811 |
| ENSMUSG00( | 11.4249599 | -1.2099302 | 0.57159869 | 0.00237544 | 0.01027082 | Gm6296 |
| ENSMUSG00( | 21.793691 | -1.2102781 | 0.40212867 | 2.21E-04 | 0.00126139 | 2900005J15R |
| ENSMUSG00( | 59.4193617 | -1.2106539 | 0.18805205 | 1.07E-11 | 2.26E-10 | G530011O06I |
| ENSMUSG00( | 38.5213663 | -1.212466 | 0.29015023 | 2.65E-06 | 2.31E-05 | |
| ENSMUSG00( | 806.952134 | -1.2174968 | 0.08059863 | 8.53E-53 | 4.97E-50 | Arhgap25 |
| ENSMUSG00( | 20.565247 | -1.2175824 | 0.41403028 | 2.63E-04 | 0.00147621 | Slc16a11 |
| ENSMUSG00( | 392.758995 | -1.2188884 | 0.08711803 | 1.63E-45 | 6.71E-43 | Icosl |
| ENSMUSG00( | 4.22480152 | -1.2205077 | 1.26178721 | 0.01020656 | 0.03547948 | Gm37962 |
| ENSMUSG00( | 14.7816969 | -1.2225708 | 0.52586093 | 0.00151887 | 0.00694795 | Gm50387 |
| ENSMUSG00( | 4519.85293 | -1.2234951 | 0.46071544 | 5.52E-04 | 0.00284314 | Ier3 |
| ENSMUSG00( | 30.3068585 | -1.2250295 | 0.29992072 | 4.02E-06 | 3.40E-05 | Gm9115 |
| ENSMUSG00( | 7.3587366 | -1.2286607 | 1.8444906 | 0.00837202 | 0.03005642 | Nbl1 |
| ENSMUSG00( | 29.0413386 | -1.2293799 | 0.36924778 | 6.77E-05 | 4.39E-04 | Slc6a19 |
| ENSMUSG00( | 14.6060942 | -1.2309675 | 0.45061735 | 4.69E-04 | 0.00246784 | |
| ENSMUSG00( | 28.6761098 | -1.23107 | 0.36258638 | 5.78E-05 | 3.80E-04 | 9330162G02F |
| ENSMUSG00( | 6.26445499 | -1.2347154 | 0.71831101 | 0.00472546 | 0.01852556 | |
| ENSMUSG00( | 9.77402531 | -1.2356558 | 0.64702826 | 0.00383452 | 0.01551062 | Gm14321 |
| ENSMUSG00( | 109.236157 | -1.2376578 | 0.22121681 | 1.98E-09 | 3.03E-08 | Btg3 |
| ENSMUSG00( | 44.1540127 | -1.2405803 | 0.28823654 | 1.46E-06 | 1.34E-05 | Rbpms2 |
| ENSMUSG00( | 50.2012103 | -1.2406679 | 0.28903576 | 1.63E-06 | 1.48E-05 | Cox6a2 |
| ENSMUSG00( | 8.02013298 | -1.241019 | 0.63594391 | 0.00325195 | 0.01344919 | Klf1 |
| ENSMUSG00( | 11.1534129 | -1.2422184 | 0.59748795 | 0.00228421 | 0.00992389 | D930030I03R |
| ENSMUSG00( | 120.073713 | -1.243067 | 0.20205382 | 7.99E-11 | 1.49E-09 | Plekhg2 |

| | | | | | | |
|---|---|---|---|---|---|---|
| ENSMUSG00( | 43402.2273 | -1.2455348 | 0.17618431 | 1.45E-13 | 4.04E-12 | Il1rn |
| ENSMUSG00( | 22650.3653 | -1.2456998 | 0.40301589 | 1.41E-04 | 8.50E-04 | Ptgs2 |
| ENSMUSG00( | 51.2919868 | -1.2509326 | 0.25201382 | 6.45E-08 | 7.62E-07 | Gm15726 |
| ENSMUSG00( | 12.5268102 | -1.2516981 | 0.53360629 | 0.00142317 | 0.00656504 | Gm12992 |
| ENSMUSG00( | 159.937337 | -1.256185 | 0.14266216 | 1.31E-19 | 7.69E-18 | Igtp |
| ENSMUSG00( | 52.3505521 | -1.2563124 | 0.25552131 | 8.62E-08 | 9.85E-07 | Gm17092 |
| ENSMUSG00( | 14.1806612 | -1.2596686 | 0.51744113 | 0.00103649 | 0.00499557 | 4930515G01I |
| ENSMUSG00( | 18.0654445 | -1.2637486 | 0.41759891 | 1.93E-04 | 0.00111526 | Jph3 |
| ENSMUSG00( | 469.482635 | -1.2656005 | 0.12694478 | 2.05E-24 | 2.00E-22 | Irgm1 |
| ENSMUSG00( | 93.6106657 | -1.2662658 | 0.21487394 | 3.82E-10 | 6.45E-09 | Tlr3 |
| ENSMUSG00( | 8.37735322 | -1.2672879 | 0.89005487 | 0.00677225 | 0.0251011 | Gm13369 |
| ENSMUSG00( | 846.455021 | -1.2709421 | 0.17203196 | 1.74E-14 | 5.59E-13 | Iffo2 |
| ENSMUSG00( | 18.3368406 | -1.2762727 | 0.46449708 | 4.07E-04 | 0.00216924 | Mx1 |
| ENSMUSG00( | 36.953323 | -1.2783199 | 0.29729876 | 1.54E-06 | 1.41E-05 | |
| ENSMUSG00( | 210.333417 | -1.2796501 | 0.44241034 | 2.42E-04 | 0.00136957 | Dusp14 |
| ENSMUSG00( | 44.1293965 | -1.2804096 | 0.34744694 | 1.84E-05 | 1.35E-04 | Gm26520 |
| ENSMUSG00( | 26230.5058 | -1.2808164 | 0.20744107 | 7.05E-11 | 1.33E-09 | Ccl3 |
| ENSMUSG00( | 15.9069051 | -1.2811072 | 0.54260878 | 0.00114473 | 0.00544608 | A930037H05I |
| ENSMUSG00( | 6294.84583 | -1.2867067 | 0.35391665 | 2.06E-05 | 1.50E-04 | Plau |
| ENSMUSG00( | 1272.23949 | -1.2870546 | 0.06519333 | 2.41E-87 | 2.72E-84 | Rybp |
| ENSMUSG00( | 27.7298691 | -1.2897352 | 0.36208849 | 2.86E-05 | 2.01E-04 | Cd69 |
| ENSMUSG00( | 144.196124 | -1.2920888 | 0.15931952 | 2.32E-17 | 1.06E-15 | Xaf1 |
| ENSMUSG00( | 6.12315069 | -1.2984343 | 1.03732099 | 0.00901702 | 0.03198441 | Gm45501 |
| ENSMUSG00( | 5.34547674 | -1.2988348 | 0.77463764 | 0.00493001 | 0.0191942 | Gm44974 |
| ENSMUSG00( | 10.9850716 | -1.2993248 | 0.66381433 | 0.00272699 | 0.01155851 | Gimap9 |
| ENSMUSG00( | 230.379765 | -1.3003844 | 0.21891355 | 2.52E-10 | 4.36E-09 | Zfp462 |
| ENSMUSG00( | 83.7974553 | -1.3012329 | 0.20842425 | 3.94E-11 | 7.70E-10 | |
| ENSMUSG00( | 17.6448256 | -1.3031578 | 0.43389935 | 2.05E-04 | 0.00117681 | Zfpm2 |
| ENSMUSG00( | 247.840637 | -1.3049583 | 0.1856667 | 2.32E-13 | 6.22E-12 | Myh10 |
| ENSMUSG00( | 9.14480187 | -1.3051592 | 0.50160413 | 6.20E-04 | 0.00315626 | A630072M18I |
| ENSMUSG00( | 13.6026581 | -1.3072919 | 0.47531908 | 4.21E-04 | 0.00223738 | Gm37139 |
| ENSMUSG00( | 14.2252348 | -1.3081467 | 0.66890505 | 0.0028346 | 0.01194182 | Gm37309 |
| ENSMUSG00( | 892.124617 | -1.3099703 | 0.16658168 | 3.30E-16 | 1.30E-14 | Jarid2 |
| ENSMUSG00( | 163.23289 | -1.3101921 | 0.13273296 | 6.07E-24 | 5.60E-22 | Mxd1 |
| ENSMUSG00( | 10.8609295 | -1.3114497 | 0.49529826 | 5.38E-04 | 0.00278582 | Xcr1 |
| ENSMUSG00( | 92.1778806 | -1.3139066 | 0.20347641 | 1.00E-11 | 2.13E-10 | Maml2 |
| ENSMUSG00( | 7.66252579 | -1.3140609 | 0.82056357 | 0.00493722 | 0.01921399 | |
| ENSMUSG00( | 41.7876527 | -1.3169729 | 0.31625329 | 2.48E-06 | 2.18E-05 | Gm49249 |
| ENSMUSG00( | 6.94612882 | -1.317874 | 0.92731172 | 0.00608741 | 0.02297273 | |
| ENSMUSG00( | 91.0694701 | -1.3200718 | 0.23516929 | 1.68E-09 | 2.59E-08 | Mrgpre |
| ENSMUSG00( | 193.287377 | -1.3237981 | 0.13766507 | 5.82E-23 | 4.78E-21 | Tmem156 |

| | | | | | | |
|---|---|---|---|---|---|---|
| ENSMUSG00( | 11.0130572 | -1.3269045 | 0.55021055 | 0.00103217 | 0.00497607 | Gm37933 |
| ENSMUSG00( | 18.0819156 | -1.3280166 | 0.51498109 | 6.27E-04 | 0.00318977 | |
| ENSMUSG00( | 21.9322069 | -1.3290251 | 0.56767899 | 0.00117392 | 0.00556441 | Prss16 |
| ENSMUSG00( | 19493.5122 | -1.3331185 | 0.45882216 | 2.30E-04 | 0.00130569 | Ccl2 |
| ENSMUSG00( | 40.9929537 | -1.3339003 | 0.69497987 | 0.00253221 | 0.01084744 | Egr2 |
| ENSMUSG00( | 21.5688983 | -1.3355027 | 0.59159976 | 0.00135379 | 0.00628836 | Glcci1 |
| ENSMUSG00( | 873.155916 | -1.3365708 | 0.12073932 | 2.89E-29 | 4.28E-27 | Iqsec1 |
| ENSMUSG00( | 3.80318195 | -1.3400792 | 1.76058569 | 0.01269582 | 0.04242662 | Gm11410 |
| ENSMUSG00( | 115.96121 | -1.3411199 | 0.2025487 | 3.01E-12 | 6.88E-11 | Raet1e |
| ENSMUSG00( | 7.58302861 | -1.3420916 | 0.73807991 | 0.0034996 | 0.01430638 | Gm16315 |
| ENSMUSG00( | 244.593267 | -1.3441492 | 0.20318652 | 3.32E-12 | 7.53E-11 | Vopp1 |
| ENSMUSG00( | 426.304844 | -1.3501252 | 0.10751038 | 2.88E-37 | 7.77E-35 | St7l |
| ENSMUSG00( | 85.6735532 | -1.350614 | 0.18800853 | 6.05E-14 | 1.76E-12 | Tfcp2l1 |
| ENSMUSG00( | 12.8054984 | -1.3526251 | 0.59042507 | 0.00128441 | 0.0060093 | Gm5112 |
| ENSMUSG00( | 23.9604368 | -1.3548165 | 0.37835364 | 2.62E-05 | 1.86E-04 | Gm45719 |
| ENSMUSG00( | 2.07681618 | -1.3551686 | 2.95176635 | 0.01487996 | 0.04832242 | Gm55508 |
| ENSMUSG00( | 182.271726 | -1.3578185 | 0.16658995 | 3.35E-17 | 1.50E-15 | 8430429K09F |
| ENSMUSG00( | 3.16702876 | -1.3579323 | 1.7550847 | 0.01148396 | 0.03907054 | Gm12227 |
| ENSMUSG00( | 89.9652667 | -1.3591045 | 0.20958222 | 7.44E-12 | 1.60E-10 | Gm7993 |
| ENSMUSG00( | 14.5729224 | -1.3593699 | 0.47261657 | 2.67E-04 | 0.0014951 | Gm26981 |
| ENSMUSG00( | 1290.75528 | -1.3602269 | 0.22872935 | 2.29E-10 | 3.99E-09 | Cd28 |
| ENSMUSG00( | 8.39744895 | -1.3608654 | 0.68811122 | 0.00251098 | 0.0107718 | Mmp8 |
| ENSMUSG00( | 3.25156137 | -1.3631675 | 1.42023609 | 0.00902105 | 0.03199243 | Gm47765 |
| ENSMUSG00( | 201.679548 | -1.364765 | 0.17098241 | 1.58E-16 | 6.46E-15 | Abcb1a |
| ENSMUSG00( | 2197.75726 | -1.3650286 | 0.21475816 | 1.73E-11 | 3.56E-10 | Dcstamp |
| ENSMUSG00( | 36.0397119 | -1.3660202 | 0.36940956 | 1.66E-05 | 1.23E-04 | 4921536K21F |
| ENSMUSG00( | 55.238563 | -1.36684 | 0.28121506 | 9.56E-08 | 1.08E-06 | Gm43197 |
| ENSMUSG00( | 6.2946821 | -1.3722915 | 0.95929245 | 0.00726385 | 0.02666068 | Igfbp6 |
| ENSMUSG00( | 6.0147848 | -1.3726755 | 0.84331089 | 0.00476981 | 0.01865087 | BC030343 |
| ENSMUSG00( | 44.707348 | -1.376005 | 0.27742153 | 5.84E-08 | 6.96E-07 | Gm10425 |
| ENSMUSG00( | 4.45780457 | -1.3765085 | 0.92621158 | 0.00578553 | 0.02200816 | Ms4a4b |
| ENSMUSG00( | 10.866533 | -1.3769362 | 0.51594517 | 4.95E-04 | 0.00258616 | Gm44116 |
| ENSMUSG00( | 125.591612 | -1.3776183 | 0.16531456 | 6.97E-18 | 3.34E-16 | Lims2 |
| ENSMUSG00( | 374.498484 | -1.3779724 | 0.14324858 | 9.17E-23 | 7.39E-21 | Ifit2 |
| ENSMUSG00( | 4.09134196 | -1.3803614 | 1.32429851 | 0.00881727 | 0.03138676 | Gm10390 |
| ENSMUSG00( | 291.883257 | -1.3819786 | 0.17879197 | 8.86E-16 | 3.34E-14 | Oas2 |
| ENSMUSG00( | 165.604844 | -1.3833514 | 0.18047538 | 1.54E-15 | 5.63E-14 | Rtp4 |
| ENSMUSG00( | 174.46478 | -1.3845331 | 0.13100145 | 5.03E-27 | 5.94E-25 | Cd80 |
| ENSMUSG00( | 10.1929201 | -1.3884376 | 0.61888214 | 0.00146361 | 0.00673099 | Gm4793 |
| ENSMUSG00( | 6.1030919 | -1.3914075 | 0.6457832 | 0.00178801 | 0.00799574 | |
| ENSMUSG00( | 6.88581035 | -1.3948972 | 0.58375295 | 0.00106297 | 0.00511365 | Il1bos |

| | | | | | | |
|---|---|---|---|---|---|---|
| ENSMUSG00( | 6.58406771 | -1.3963397 | 0.93601116 | 0.00604812 | 0.0228483 | Acat3 |
| ENSMUSG00( | 28.4296895 | -1.3975827 | 0.34204286 | 3.37E-06 | 2.89E-05 | Gm10252 |
| ENSMUSG00( | 114.101366 | -1.3978501 | 0.19673101 | 1.06E-13 | 3.01E-12 | Ccr1 |
| ENSMUSG00( | 7.42381547 | -1.3988194 | 0.62255061 | 0.00149803 | 0.00686996 | |
| ENSMUSG00( | 117.282737 | -1.398985 | 0.17987651 | 8.24E-16 | 3.11E-14 | Rbp4 |
| ENSMUSG00( | 130.828707 | -1.4014293 | 0.1843191 | 2.57E-15 | 9.15E-14 | Rpgrip1 |
| ENSMUSG00( | 364.784327 | -1.4038894 | 0.23866319 | 3.27E-10 | 5.59E-09 | Adora2b |
| ENSMUSG00( | 17.6321777 | -1.4053607 | 0.47470991 | 1.99E-04 | 0.00115121 | Tmem71 |
| ENSMUSG00( | 4680.66242 | -1.4064519 | 0.11889385 | 2.38E-33 | 4.67E-31 | Procr |
| ENSMUSG00( | 145.314332 | -1.4105952 | 0.31039253 | 4.06E-07 | 4.11E-06 | 6330403L08R |
| ENSMUSG00( | 78.8252044 | -1.4109199 | 0.26337718 | 7.13E-09 | 9.82E-08 | Mir155hg |
| ENSMUSG00( | 9.21711972 | -1.4140318 | 0.57594 | 9.27E-04 | 0.0045375 | 5830454E08F |
| ENSMUSG00( | 36.7041716 | -1.4141478 | 0.31141197 | 4.52E-07 | 4.53E-06 | AV356131 |
| ENSMUSG00( | 77.3232494 | -1.4167573 | 0.23070824 | 7.43E-11 | 1.39E-09 | Gm16001 |
| ENSMUSG00( | 1070.37865 | -1.4195012 | 0.09803985 | 1.44E-48 | 6.65E-46 | Grap |
| ENSMUSG00( | 90.9552755 | -1.4206564 | 0.25492993 | 2.09E-09 | 3.18E-08 | |
| ENSMUSG00( | 148.609074 | -1.4210466 | 0.17264912 | 1.55E-17 | 7.20E-16 | Zfp811 |
| ENSMUSG00( | 3.97235601 | -1.4214129 | 1.37651505 | 0.01110597 | 0.03802048 | Usp50 |
| ENSMUSG00( | 659.312525 | -1.4219008 | 0.1493388 | 1.56E-22 | 1.22E-20 | Rhou |
| ENSMUSG00( | 5.89564125 | -1.4229588 | 0.97999791 | 0.00604247 | 0.02283172 | Gpr18 |
| ENSMUSG00( | 8.0908135 | -1.4249577 | 0.57293149 | 7.17E-04 | 0.00360275 | Gm30873 |
| ENSMUSG00( | 18.0467056 | -1.4252422 | 0.4559929 | 1.23E-04 | 7.51E-04 | Gm34425 |
| ENSMUSG00( | 61.7549874 | -1.4268795 | 0.31094652 | 3.16E-07 | 3.25E-06 | Cdcp1 |
| ENSMUSG00( | 23.6344916 | -1.4297786 | 0.40243202 | 2.75E-05 | 1.94E-04 | Pla2g2a |
| ENSMUSG00( | 4.02245964 | -1.4303256 | 1.44281439 | 0.00777616 | 0.02819736 | Gm47705 |
| ENSMUSG00( | 201.305988 | -1.4318799 | 0.14821574 | 5.68E-23 | 4.69E-21 | Sema4c |
| ENSMUSG00( | 121.550689 | -1.4334537 | 0.38553221 | 1.42E-05 | 1.07E-04 | Arrdc3 |
| ENSMUSG00( | 7.9893763 | -1.4362082 | 0.83876089 | 0.00382348 | 0.01547636 | Gm17034 |
| ENSMUSG00( | 7.83472383 | -1.4369653 | 0.80277662 | 0.00374128 | 0.01518785 | 2310034O05I |
| ENSMUSG00( | 23020.1442 | -1.4398325 | 0.0764339 | 1.96E-80 | 1.97E-77 | Odc1 |
| ENSMUSG00( | 175.975641 | -1.441315 | 0.20647694 | 2.38E-13 | 6.38E-12 | Gbp7 |
| ENSMUSG00( | 10.6962591 | -1.4446748 | 0.5332806 | 4.15E-04 | 0.00220622 | Gfra1 |
| ENSMUSG00( | 26.3967611 | -1.4456348 | 0.37800393 | 9.80E-06 | 7.57E-05 | Lncpint |
| ENSMUSG00( | 108.541821 | -1.4466184 | 0.18165371 | 1.35E-16 | 5.57E-15 | Mtmr7 |
| ENSMUSG00( | 7.24168558 | -1.4488143 | 0.60823933 | 0.00109287 | 0.00523102 | |
| ENSMUSG00( | 9.00928358 | -1.4522302 | 0.7592568 | 0.00288671 | 0.01212175 | Plekhh1 |
| ENSMUSG00( | 8.63178026 | -1.4533108 | 0.72643858 | 0.0021444 | 0.00937962 | Sema3c |
| ENSMUSG00( | 20.2949836 | -1.4566476 | 0.45448679 | 9.50E-05 | 5.95E-04 | Gm8251 |
| ENSMUSG00( | 1662.27238 | -1.4568884 | 0.30401658 | 1.18E-07 | 1.31E-06 | Cdkn1a |
| ENSMUSG00( | 22.401987 | -1.4573049 | 0.37857268 | 8.28E-06 | 6.51E-05 | |
| ENSMUSG00( | 4.8444266 | -1.457663 | 0.97582355 | 0.00654095 | 0.02441407 | Smim6 |

| | | | | | | |
|---|---|---|---|---|---|---|
| ENSMUSG00( | 29.9866258 | -1.4594787 | 0.40956873 | 2.66E-05 | 1.88E-04 | Gm36874 |
| ENSMUSG00( | 4.09950838 | -1.4682045 | 1.06928989 | 0.0063944 | 0.02396608 | |
| ENSMUSG00( | 900.388441 | -1.4728426 | 0.12556812 | 4.42E-33 | 8.49E-31 | Bcl2a1d |
| ENSMUSG00( | 13.6880363 | -1.4755053 | 0.71253864 | 0.00211195 | 0.00925333 | Gm42984 |
| ENSMUSG00( | 10.6571792 | -1.4768059 | 0.79818751 | 0.00278417 | 0.011754 | Gm13748 |
| ENSMUSG00( | 9.41003692 | -1.4811932 | 0.6815429 | 0.00141475 | 0.00653285 | Gm12867 |
| ENSMUSG00( | 351.478053 | -1.4835303 | 0.49892015 | 1.70E-04 | 9.97E-04 | Mmp9 |
| ENSMUSG00( | 13.40715 | -1.4843397 | 0.63077718 | 0.00103012 | 0.00496949 | Ly6a |
| ENSMUSG00( | 579.429368 | -1.4848999 | 0.16021207 | 1.52E-21 | 1.11E-19 | Samd9l |
| ENSMUSG00( | 92.6877122 | -1.4873953 | 0.21771574 | 6.81E-13 | 1.71E-11 | Tnfrsf8 |
| ENSMUSG00( | 199.327924 | -1.4886232 | 0.14276427 | 2.23E-26 | 2.49E-24 | Trmt12 |
| ENSMUSG00( | 37.5274918 | -1.4900734 | 0.40767006 | 1.74E-05 | 1.28E-04 | Pkdcc |
| ENSMUSG00( | 29.1565266 | -1.4996369 | 0.33640998 | 6.56E-07 | 6.41E-06 | Baiap2l1 |
| ENSMUSG00( | 1571.59127 | -1.5116329 | 0.39663163 | 9.46E-06 | 7.34E-05 | Dusp5 |
| ENSMUSG00( | 6.60623832 | -1.5133516 | 0.70496243 | 0.00176602 | 0.00790637 | Espn |
| ENSMUSG00( | 13.69864 | -1.5153771 | 0.52257801 | 2.40E-04 | 0.00135877 | Gm15122 |
| ENSMUSG00( | 18.5054967 | -1.5164324 | 0.33583352 | 4.85E-07 | 4.84E-06 | Gm21860 |
| ENSMUSG00( | 79.5362883 | -1.519966 | 0.26361428 | 6.42E-10 | 1.05E-08 | Mxd3 |
| ENSMUSG00( | 870.057202 | -1.5200624 | 0.2961934 | 1.96E-08 | 2.51E-07 | Il6 |
| ENSMUSG00( | 242.25075 | -1.5272597 | 0.12272862 | 1.27E-36 | 3.10E-34 | Rps6ka2 |
| ENSMUSG00( | 5.48098652 | -1.5277947 | 0.95170908 | 0.00475328 | 0.0186104 | Caprin2 |
| ENSMUSG00( | 4.99970125 | -1.5294635 | 1.2858119 | 0.01171815 | 0.03975501 | |
| ENSMUSG00( | 12.2720163 | -1.5318725 | 0.58246694 | 5.38E-04 | 0.00278691 | |
| ENSMUSG00( | 793.429875 | -1.5409332 | 0.15823715 | 1.66E-23 | 1.47E-21 | Hmga2 |
| ENSMUSG00( | 5.86777294 | -1.5449977 | 1.07612626 | 0.00613244 | 0.02312334 | Gm46367 |
| ENSMUSG00( | 54.615961 | -1.5468668 | 0.2711067 | 8.78E-10 | 1.40E-08 | Ryr1 |
| ENSMUSG00( | 22.3586104 | -1.5489337 | 0.3835949 | 4.08E-06 | 3.44E-05 | Gm32401 |
| ENSMUSG00( | 14.6014626 | -1.5502652 | 1.11533417 | 0.01054319 | 0.03647424 | Endod1 |
| ENSMUSG00( | 36.6870974 | -1.5529078 | 0.44712876 | 3.21E-05 | 2.23E-04 | B330016D10I |
| ENSMUSG00( | 4.9500496 | -1.5603175 | 1.28841425 | 0.00654751 | 0.02442339 | |
| ENSMUSG00( | 37.9948787 | -1.5633422 | 0.43411027 | 2.26E-05 | 1.63E-04 | Ifnlr1 |
| ENSMUSG00( | 5.18996467 | -1.5785905 | 0.8637875 | 0.00243644 | 0.01049941 | Adprhl1 |
| ENSMUSG00( | 7.70517543 | -1.580423 | 0.76666181 | 0.00187135 | 0.00831001 | Gm44699 |
| ENSMUSG00( | 19.2554425 | -1.5878987 | 0.43231475 | 1.67E-05 | 1.24E-04 | Gm10138 |
| ENSMUSG00( | 4.43885216 | -1.5929207 | 1.21830537 | 0.00722858 | 0.026542 | Gm42820 |
| ENSMUSG00( | 12.9865488 | -1.5977534 | 0.4606338 | 3.76E-05 | 2.57E-04 | Gm4285 |
| ENSMUSG00( | 46.2728653 | -1.602702 | 0.27453198 | 3.75E-10 | 6.35E-09 | C430049B03I |
| ENSMUSG00( | 5.96152883 | -1.6030827 | 1.01535208 | 0.00503222 | 0.01952064 | Kazald1 |
| ENSMUSG00( | 180.365451 | -1.603429 | 0.15522727 | 5.86E-26 | 6.38E-24 | Spn |
| ENSMUSG00( | 14.0756791 | -1.6058239 | 0.50500469 | 9.09E-05 | 5.71E-04 | Gm9887 |
| ENSMUSG00( | 77.3366769 | -1.6118595 | 0.38332221 | 1.69E-06 | 1.53E-05 | Irf4 |

| | | | | | | |
|---|---|---|---|---|---|---|
| ENSMUSG00( | 20.6729394 | -1.6158773 | 0.34657952 | 2.20E-07 | 2.32E-06 | B930059L03F |
| ENSMUSG00( | 18.6518615 | -1.6160111 | 0.52691701 | 1.28E-04 | 7.78E-04 | Rab30 |
| ENSMUSG00( | 28.1540583 | -1.6176392 | 0.30613028 | 1.01E-08 | 1.36E-07 | |
| ENSMUSG00( | 617.430774 | -1.6225592 | 0.13662596 | 1.25E-33 | 2.51E-31 | |
| ENSMUSG00( | 30.7986039 | -1.626124 | 0.4025985 | 3.75E-06 | 3.20E-05 | Arhgdig |
| ENSMUSG00( | 3.72687119 | -1.6301376 | 1.27800531 | 0.00765849 | 0.02783772 | Gpr31a |
| ENSMUSG00( | 86.9968447 | -1.6338491 | 0.21939147 | 7.89E-15 | 2.64E-13 | Mipol1 |
| ENSMUSG00( | 3.19105355 | -1.6346806 | 1.64913457 | 0.00877732 | 0.03128153 | Gm44553 |
| ENSMUSG00( | 4.15019964 | -1.6354213 | 1.13852945 | 0.00601632 | 0.0227597 | A930035D04F |
| ENSMUSG00( | 400.466679 | -1.644546 | 0.10833098 | 6.31E-53 | 3.80E-50 | Gramd3 |
| ENSMUSG00( | 11.5119109 | -1.6473829 | 0.57225691 | 2.76E-04 | 0.00153727 | Ston2 |
| ENSMUSG00( | 63.8894351 | -1.6524638 | 0.24313265 | 7.95E-13 | 1.97E-11 | Slc45a3 |
| ENSMUSG00( | 9.51645228 | -1.6625124 | 0.54341797 | 1.39E-04 | 8.36E-04 | Gm37170 |
| ENSMUSG00( | 8.48893027 | -1.6628379 | 0.58407103 | 2.48E-04 | 0.0013988 | Slc24a1 |
| ENSMUSG00( | 2.78851696 | -1.6634274 | 1.50538136 | 0.00771492 | 0.02801464 | Tbr1 |
| ENSMUSG00( | 13.5379258 | -1.6732922 | 0.46887801 | 2.39E-05 | 1.71E-04 | |
| ENSMUSG00( | 384.639099 | -1.6738104 | 0.15518346 | 3.55E-28 | 4.74E-26 | Creb5 |
| ENSMUSG00( | 22.653074 | -1.6767026 | 0.3915928 | 1.27E-06 | 1.18E-05 | Ifi205 |
| ENSMUSG00( | 7.44843896 | -1.6801146 | 0.63455011 | 4.28E-04 | 0.00226952 | Gm30262 |
| ENSMUSG00( | 75.1454596 | -1.6868596 | 0.2188116 | 1.28E-15 | 4.74E-14 | F730311O21F |
| ENSMUSG00( | 3.80186988 | -1.6884098 | 0.99724974 | 0.00411308 | 0.01649339 | Gm17359 |
| ENSMUSG00( | 4.63766827 | -1.6896056 | 0.81770126 | 0.00172255 | 0.00774441 | Gm11563 |
| ENSMUSG00( | 13.3523505 | -1.6931236 | 0.66683052 | 7.98E-04 | 0.00397125 | Esr1 |
| ENSMUSG00( | 51.6626859 | -1.6945958 | 0.27498715 | 5.54E-11 | 1.06E-09 | H2-Q5 |
| ENSMUSG00( | 5.27865781 | -1.7016764 | 0.85128677 | 0.00225034 | 0.00979084 | Gm31462 |
| ENSMUSG00( | 16.9100055 | -1.7040164 | 0.56239896 | 1.31E-04 | 7.96E-04 | Gm2619 |
| ENSMUSG00( | 8.34902601 | -1.7059693 | 0.80609949 | 0.00151508 | 0.00693592 | Gm45407 |
| ENSMUSG00( | 105.679653 | -1.7060226 | 0.24311015 | 1.64E-13 | 4.53E-12 | Lix1 |
| ENSMUSG00( | 13.6334538 | -1.7080522 | 0.49875841 | 3.75E-05 | 2.57E-04 | |
| ENSMUSG00( | 22.8436544 | -1.7116965 | 0.36202676 | 1.51E-07 | 1.64E-06 | C230013L11F |
| ENSMUSG00( | 179.145899 | -1.7129332 | 0.23113709 | 8.49E-15 | 2.83E-13 | |
| ENSMUSG00( | 3.58029354 | -1.7152886 | 1.77533449 | 0.00721254 | 0.02649387 | Gm5544 |
| ENSMUSG00( | 5.55523623 | -1.7238228 | 0.90429394 | 0.00209003 | 0.00916843 | Gm7582 |
| ENSMUSG00( | 40.7481383 | -1.7243774 | 0.29716405 | 4.88E-10 | 8.10E-09 | Oas1b |
| ENSMUSG00( | 14.699104 | -1.7252572 | 0.50104765 | 3.90E-05 | 2.67E-04 | Tomt |
| ENSMUSG00( | 415.450825 | -1.7357619 | 0.14133087 | 8.83E-36 | 1.99E-33 | Ifi203 |
| ENSMUSG00( | 9.09037391 | -1.7432259 | 0.92342949 | 0.00300997 | 0.01258671 | BC046401 |
| ENSMUSG00( | 16.7923172 | -1.7455535 | 0.53574117 | 6.29E-05 | 4.11E-04 | Gm26549 |
| ENSMUSG00( | 18.3659616 | -1.7477547 | 0.59779718 | 1.77E-04 | 0.00103753 | Cda |
| ENSMUSG00( | 13.8382512 | -1.7489794 | 0.53089988 | 6.86E-05 | 4.44E-04 | Gm26724 |
| ENSMUSG00( | 14.9505927 | -1.75204 | 0.60614464 | 2.25E-04 | 0.00128311 | 9430034N14F |

| | | | | | | |
|---|---|---|---|---|---|---|
| ENSMUSG00( | 7.01476487 | -1.7629083 | 0.6417898 | 3.49E-04 | 0.00189507 | Gm16181 |
| ENSMUSG00( | 63.4345971 | -1.7665331 | 0.2853758 | 4.04E-11 | 7.89E-10 | Plekhh2 |
| ENSMUSG00( | 7.25092954 | -1.7685547 | 0.85752762 | 0.00157831 | 0.00718532 | Cd22 |
| ENSMUSG00( | 15.7375117 | -1.7730034 | 0.57155609 | 1.32E-04 | 8.01E-04 | Gm10357 |
| ENSMUSG00( | 411.77443 | -1.7847052 | 0.19928359 | 2.30E-20 | 1.50E-18 | Gprc5a |
| ENSMUSG00( | 58.7312245 | -1.7859139 | 0.26183418 | 6.73E-13 | 1.69E-11 | Dclk2 |
| ENSMUSG00( | 1364.46695 | -1.7884301 | 0.22434562 | 1.02E-16 | 4.33E-15 | Eps8 |
| ENSMUSG00( | 3.78825626 | -1.7926555 | 1.41648779 | 0.00831086 | 0.02986653 | Gm10643 |
| ENSMUSG00( | 2.55023084 | -1.7934377 | 1.4587359 | 0.00632219 | 0.02374469 | Gm26616 |
| ENSMUSG00( | 220.062147 | -1.7936901 | 0.2240419 | 8.23E-17 | 3.53E-15 | Ifit1 |
| ENSMUSG00( | 76.2727965 | -1.7962355 | 0.2463607 | 2.53E-14 | 7.90E-13 | Ifi209 |
| ENSMUSG00( | 7139.60807 | -1.8003093 | 0.15098134 | 3.00E-34 | 6.23E-32 | Il1a |
| ENSMUSG00( | 191.967971 | -1.8074157 | 0.16132483 | 2.65E-30 | 4.12E-28 | Tspan13 |
| ENSMUSG00( | 15.8709905 | -1.8085805 | 0.5105117 | 3.02E-05 | 2.11E-04 | Dnajc28 |
| ENSMUSG00( | 426.939439 | -1.8092172 | 0.23338621 | 6.41E-16 | 2.45E-14 | Cd5 |
| ENSMUSG00( | 12.5972667 | -1.8230577 | 0.48674137 | 1.26E-05 | 9.55E-05 | Prr5l |
| ENSMUSG00( | 210.580388 | -1.828328 | 0.42322402 | 8.38E-07 | 8.05E-06 | Fosl1 |
| ENSMUSG00( | 388.365923 | -1.8286043 | 0.14666616 | 9.47E-37 | 2.36E-34 | Oasl2 |
| ENSMUSG00( | 6.27404218 | -1.8371532 | 1.47270891 | 0.00615884 | 0.02320896 | 1700023B13F |
| ENSMUSG00( | 3.71940066 | -1.8413359 | 1.26257757 | 0.00451749 | 0.01777583 | Gm25280 |
| ENSMUSG00( | 24.7166632 | -1.8433592 | 0.38960971 | 1.70E-07 | 1.83E-06 | Hgf |
| ENSMUSG00( | 109.932722 | -1.853253 | 0.26227477 | 9.21E-14 | 2.63E-12 | Zfp469 |
| ENSMUSG00( | 23.3813662 | -1.8532644 | 0.3939735 | 1.67E-07 | 1.81E-06 | Gm9260 |
| ENSMUSG00( | 49.4975572 | -1.8534912 | 0.40068517 | 2.34E-07 | 2.46E-06 | Slc28a2 |
| ENSMUSG00( | 13.1063158 | -1.8600061 | 0.5374469 | 3.36E-05 | 2.32E-04 | Gbp6 |
| ENSMUSG00( | 16.2823333 | -1.8606418 | 0.47642226 | 6.06E-06 | 4.93E-05 | Slc26a1 |
| ENSMUSG00( | 4.8593661 | -1.8679041 | 1.01868843 | 0.00315869 | 0.01312544 | |
| ENSMUSG00( | 3.89467269 | -1.8718651 | 2.05962689 | 0.00967511 | 0.0338732 | Gm33280 |
| ENSMUSG00( | 4.35423287 | -1.8739609 | 1.34526802 | 0.00736031 | 0.02693265 | Gm12059 |
| ENSMUSG00( | 10.8394517 | -1.8828248 | 0.55546636 | 3.66E-05 | 2.51E-04 | Gm5475 |
| ENSMUSG00( | 86.8020425 | -1.8883204 | 0.20751285 | 6.87E-21 | 4.75E-19 | Wwtr1 |
| ENSMUSG00( | 4.73653823 | -1.8926296 | 0.99990407 | 0.00273048 | 0.01156788 | Gm13207 |
| ENSMUSG00( | 176.487542 | -1.9056526 | 0.21227179 | 2.00E-20 | 1.31E-18 | Celsr1 |
| ENSMUSG00( | 19.7651128 | -1.9059778 | 0.56433927 | 3.73E-05 | 2.56E-04 | Tgm1 |
| ENSMUSG00( | 875.080735 | -1.912599 | 0.24496257 | 3.89E-16 | 1.52E-14 | Serpinb1a |
| ENSMUSG00( | 12.8591611 | -1.9129185 | 0.59548135 | 7.00E-05 | 4.52E-04 | |
| ENSMUSG00( | 13.0068641 | -1.9213945 | 0.62146007 | 1.24E-04 | 7.61E-04 | Ccr5 |
| ENSMUSG00( | 173.635236 | -1.9221742 | 0.15608113 | 4.11E-36 | 9.77E-34 | Gask1b |
| ENSMUSG00( | 12.868518 | -1.9249215 | 0.66665897 | 1.76E-04 | 0.00103203 | Pdpn |
| ENSMUSG00( | 13.482004 | -1.9256334 | 0.49799805 | 7.76E-06 | 6.17E-05 | C130013H08I |
| ENSMUSG00( | 5.70880723 | -1.9293066 | 0.79093336 | 7.96E-04 | 0.00396488 | Gm42930 |

| | | | | | | |
|---|---|---|---|---|---|---|
| ENSMUSG00( | 19.7095779 | -1.9305591 | 0.39967111 | 9.62E-08 | 1.09E-06 | Rnf43 |
| ENSMUSG00( | 5.23135974 | -1.9311618 | 0.81172835 | 8.84E-04 | 0.00434642 | Gm28731 |
| ENSMUSG00( | 65.1272482 | -1.9337993 | 0.27599653 | 1.88E-13 | 5.13E-12 | Atp6v0d2 |
| ENSMUSG00( | 6.18452756 | -1.9501791 | 0.70922052 | 3.64E-04 | 0.00196011 | |
| ENSMUSG00( | 22.3128461 | -1.9518117 | 0.39770636 | 5.83E-08 | 6.95E-07 | Trim66 |
| ENSMUSG00( | 22.7770185 | -1.9535899 | 0.46646475 | 1.49E-06 | 1.37E-05 | 4921529L05R |
| ENSMUSG00( | 3.40780571 | -1.9554796 | 1.44212505 | 0.00704346 | 0.02596771 | Gm13833 |
| ENSMUSG00( | 11.6553149 | -1.9557774 | 0.5970099 | 6.28E-05 | 4.10E-04 | A430108G06I |
| ENSMUSG00( | 7.28688022 | -1.9562359 | 0.8806414 | 0.00134579 | 0.00625602 | Zeb1 |
| ENSMUSG00( | 3.12005879 | -1.9575595 | 1.55569762 | 0.00735211 | 0.02690966 | Rny1 |
| ENSMUSG00( | 75.852775 | -1.9582018 | 0.27909579 | 1.47E-13 | 4.07E-12 | Ccl22 |
| ENSMUSG00( | 22.8783338 | -1.9612018 | 0.49889736 | 6.74E-06 | 5.43E-05 | Mras |
| ENSMUSG00( | 3.10607708 | -1.9632002 | 1.54157994 | 0.00722356 | 0.02652897 | Gm3625 |
| ENSMUSG00( | 65.7004009 | -1.9715752 | 0.27962848 | 1.41E-13 | 3.94E-12 | Unc13a |
| ENSMUSG00( | 4.37788333 | -1.9736504 | 1.0479046 | 0.00202236 | 0.00889749 | |
| ENSMUSG00( | 1326.00604 | -2.0035327 | 0.16014877 | 4.48E-37 | 1.17E-34 | Bhlhe40 |
| ENSMUSG00( | 9.57276175 | -2.0114558 | 0.62831126 | 8.15E-05 | 5.18E-04 | Gm12250 |
| ENSMUSG00( | 42.6056105 | -2.0224855 | 0.38157539 | 6.98E-09 | 9.63E-08 | Sh3bp4 |
| ENSMUSG00( | 142.355917 | -2.0246041 | 0.23256963 | 2.03E-19 | 1.16E-17 | Gimap6 |
| ENSMUSG00( | 19.2760278 | -2.0299742 | 0.40186474 | 2.47E-08 | 3.12E-07 | Itgb3 |
| ENSMUSG00( | 4107.19636 | -2.0423725 | 0.14707867 | 4.97E-45 | 1.83E-42 | Furin |
| ENSMUSG00( | 22.8163333 | -2.045858 | 0.40698313 | 3.36E-08 | 4.16E-07 | Gcnt2 |
| ENSMUSG00( | 472.677916 | -2.0538565 | 0.13785095 | 2.27E-51 | 1.17E-48 | Usp18 |
| ENSMUSG00( | 15.3927681 | -2.0677881 | 0.737103 | 4.56E-04 | 0.00240093 | Gm39458 |
| ENSMUSG00( | 2625.35214 | -2.0795303 | 0.08517393 | 1.27E-132 | 1.15E-128 | Nckap5l |
| ENSMUSG00( | 720.158166 | -2.0946141 | 0.2128953 | 5.84E-24 | 5.41E-22 | Lgals3bp |
| ENSMUSG00( | 26.883157 | -2.0980983 | 0.40341774 | 1.71E-08 | 2.22E-07 | Ceacam19 |
| ENSMUSG00( | 2.44521952 | -2.1009159 | 1.24957409 | 0.00378479 | 0.01533349 | Cpvl |
| ENSMUSG00( | 4.05242588 | -2.1067641 | 1.02591625 | 0.00178044 | 0.00796642 | Gm38161 |
| ENSMUSG00( | 2.54457505 | -2.10951 | 1.29940783 | 0.00394093 | 0.01589122 | Gm32552 |
| ENSMUSG00( | 32.5374827 | -2.1173188 | 0.35771981 | 2.49E-10 | 4.32E-09 | Flot2 |
| ENSMUSG00( | 3.12556253 | -2.1183874 | 1.62632667 | 0.0039963 | 0.01608221 | Gm14188 |
| ENSMUSG00( | 19.5329143 | -2.1186828 | 0.42490229 | 3.60E-08 | 4.42E-07 | A1cf |
| ENSMUSG00( | 57.0849715 | -2.1232553 | 0.25506163 | 6.54E-18 | 3.15E-16 | Ifit3b |
| ENSMUSG00( | 10.472551 | -2.137031 | 0.59200214 | 1.81E-05 | 1.33E-04 | Gm47175 |
| ENSMUSG00( | 27.7610411 | -2.1377842 | 0.43194046 | 5.26E-08 | 6.32E-07 | Gm7598 |
| ENSMUSG00( | 22.1143627 | -2.1406846 | 0.57402769 | 9.18E-06 | 7.15E-05 | H2-Q6 |
| ENSMUSG00( | 9.67642934 | -2.146042 | 0.71692312 | 1.33E-04 | 8.07E-04 | Gm44851 |
| ENSMUSG00( | 7.94024168 | -2.1542142 | 0.62178731 | 3.56E-05 | 2.45E-04 | |
| ENSMUSG00( | 5.07695175 | -2.1544692 | 1.27387579 | 0.0073171 | 0.02682344 | Kcnd1 |
| ENSMUSG00( | 8.8579512 | -2.1644197 | 0.97811823 | 0.00158595 | 0.00720918 | Tdrd9 |

| | | | | | | |
|---|---|---|---|---|---|---|
| ENSMUSG00( | 4.68923713 | -2.170528 | 0.98473008 | 0.00101647 | 0.00491481 | Tm4sf4 |
| ENSMUSG00( | 5.53179591 | -2.1745198 | 1.51685043 | 0.00380091 | 0.01539534 | Gm37558 |
| ENSMUSG00( | 293.755827 | -2.1784685 | 0.17485444 | 7.44E-37 | 1.89E-34 | Ifit3 |
| ENSMUSG00( | 5.03645683 | -2.1790093 | 0.92042649 | 0.00118399 | 0.00560185 | Gm45220 |
| ENSMUSG00( | 17.1837146 | -2.1813445 | 0.43577799 | 3.56E-08 | 4.39E-07 | Pira2 |
| ENSMUSG00( | 42.4862617 | -2.1819944 | 0.90951667 | 5.41E-04 | 0.0027998 | AA467197 |
| ENSMUSG00( | 7.77592664 | -2.1908008 | 0.74713066 | 1.78E-04 | 0.00103948 | 2810029C07I |
| ENSMUSG00( | 35.5389069 | -2.2036764 | 0.34456251 | 1.16E-11 | 2.44E-10 | Gbp9 |
| ENSMUSG00( | 4.35504616 | -2.2046402 | 1.07635446 | 0.00223358 | 0.00972495 | Gm15512 |
| ENSMUSG00( | 4.13951526 | -2.2049011 | 1.24410686 | 0.00312863 | 0.01302261 | P2rx1 |
| ENSMUSG00( | 36.228519 | -2.2062229 | 0.36420157 | 7.80E-11 | 1.45E-09 | Mx2 |
| ENSMUSG00( | 2813.75778 | -2.2108891 | 0.60744471 | 1.14E-05 | 8.74E-05 | Mmp13 |
| ENSMUSG00( | 109.422329 | -2.2168591 | 0.20898748 | 2.66E-27 | 3.27E-25 | Angptl2 |
| ENSMUSG00( | 112.440731 | -2.2216437 | 0.30870058 | 3.92E-14 | 1.19E-12 | Nrcam |
| ENSMUSG00( | 2129.23764 | -2.2245502 | 0.15951986 | 2.13E-45 | 8.37E-43 | Cd83 |
| ENSMUSG00( | 4.49251875 | -2.2250505 | 0.79323392 | 2.88E-04 | 0.00159575 | |
| ENSMUSG00( | 27.3010827 | -2.2312418 | 0.36765198 | 7.37E-11 | 1.38E-09 | Chd5 |
| ENSMUSG00( | 3.50490892 | -2.2319631 | 1.23044785 | 0.00385594 | 0.01558331 | Gm42941 |
| ENSMUSG00( | 39.5581889 | -2.2381446 | 0.39143958 | 5.90E-10 | 9.68E-09 | Lif |
| ENSMUSG00( | 3.33526248 | -2.2386472 | 1.15896461 | 0.00255663 | 0.01094166 | Gm43010 |
| ENSMUSG00( | 5.12486269 | -2.2487973 | 0.73137458 | 1.16E-04 | 7.16E-04 | Pira12 |
| ENSMUSG00( | 4.2247728 | -2.2525704 | 1.09178701 | 0.00178819 | 0.00799574 | Nox3 |
| ENSMUSG00( | 2.5787615 | -2.2634606 | 1.56943687 | 0.00736629 | 0.02694907 | Gm36933 |
| ENSMUSG00( | 8.02680946 | -2.2644305 | 0.77716154 | 2.42E-04 | 0.00136908 | 4933427D14I |
| ENSMUSG00( | 902.998677 | -2.2743454 | 0.32772765 | 1.55E-13 | 4.27E-12 | Cmpk2 |
| ENSMUSG00( | 63.7867934 | -2.2792725 | 0.21146822 | 3.12E-28 | 4.27E-26 | H2-Q7 |
| ENSMUSG00( | 38.5296034 | -2.2807155 | 0.31557536 | 3.66E-14 | 1.11E-12 | Sema5a |
| ENSMUSG00( | 6280.84599 | -2.2856937 | 0.11640045 | 4.59E-87 | 4.87E-84 | Il1b |
| ENSMUSG00( | 4.33022969 | -2.3118872 | 0.89137722 | 5.67E-04 | 0.00291432 | Gm47920 |
| ENSMUSG00( | 166.022252 | -2.3185286 | 0.31231717 | 6.62E-15 | 2.25E-13 | Bcar3 |
| ENSMUSG00( | 4.56073953 | -2.3327705 | 1.09682176 | 0.00189482 | 0.00840185 | Gm47552 |
| ENSMUSG00( | 132.245783 | -2.339301 | 0.2547133 | 2.33E-21 | 1.68E-19 | Ccbe1 |
| ENSMUSG00( | 3.18627767 | -2.3395535 | 1.6030969 | 0.00691839 | 0.02557453 | Gm53056 |
| ENSMUSG00( | 237.49999 | -2.3426451 | 0.30554085 | 1.01E-15 | 3.78E-14 | Lat |
| ENSMUSG00( | 51.0480042 | -2.3565404 | 0.4387339 | 3.96E-09 | 5.72E-08 | Gm46189 |
| ENSMUSG00( | 2.76331582 | -2.3616354 | 1.70540633 | 0.0065435 | 0.02441645 | Gm17088 |
| ENSMUSG00( | 3.18872353 | -2.3635574 | 1.17237868 | 0.00160341 | 0.00727393 | Tmem82 |
| ENSMUSG00( | 6.10266733 | -2.3720565 | 0.68670224 | 3.73E-05 | 2.56E-04 | Gm1141 |
| ENSMUSG00( | 4.9567781 | -2.38244 | 0.82252286 | 2.07E-04 | 0.00118703 | Gm14186 |
| ENSMUSG00( | 12.8832529 | -2.387448 | 0.54362603 | 1.03E-06 | 9.72E-06 | 1700001G11I |
| ENSMUSG00( | 61.7924773 | -2.3952962 | 0.47835329 | 2.92E-08 | 3.65E-07 | Ifitm3 |

| | | | | | | |
|---|---|---|---|---|---|---|
| ENSMUSG00( | 459.100853 | -2.4011558 | 0.12167401 | 1.01E-87 | 1.21E-84 | Padi2 |
| ENSMUSG00( | 24.3326151 | -2.4015532 | 0.39396344 | 6.75E-11 | 1.28E-09 | Il7r |
| ENSMUSG00( | 31.2903564 | -2.4094618 | 0.40321348 | 1.30E-10 | 2.35E-09 | Rgs9 |
| ENSMUSG00( | 154.520391 | -2.4203377 | 0.22743424 | 1.01E-27 | 1.29E-25 | Ifi44 |
| ENSMUSG00( | 791.421736 | -2.4203879 | 0.13085197 | 1.62E-77 | 1.54E-74 | Card11 |
| ENSMUSG00( | 4.10250986 | -2.4246419 | 1.38032753 | 0.00191245 | 0.00847172 | Gm4610 |
| ENSMUSG00( | 86.9928064 | -2.4458611 | 0.19830948 | 5.87E-36 | 1.34E-33 | Tnfsf12 |
| ENSMUSG00( | 8.1192554 | -2.4520135 | 1.03065685 | 0.00159202 | 0.00723495 | Foxd2 |
| ENSMUSG00( | 2473.22746 | -2.466149 | 0.46162616 | 4.46E-09 | 6.36E-08 | Ccl7 |
| ENSMUSG00( | 11.570161 | -2.4668718 | 0.46729639 | 8.98E-09 | 1.22E-07 | Tnnc1 |
| ENSMUSG00( | 39.9028581 | -2.4890494 | 0.25133254 | 2.61E-24 | 2.52E-22 | |
| ENSMUSG00( | 7.7457879 | -2.5043923 | 0.71149513 | 2.56E-05 | 1.82E-04 | Gm45548 |
| ENSMUSG00( | 82.7728987 | -2.5084217 | 0.26359192 | 1.36E-22 | 1.07E-20 | Acp5 |
| ENSMUSG00( | 21.826918 | -2.5304749 | 0.46018466 | 1.97E-09 | 3.02E-08 | Gdnf |
| ENSMUSG00( | 12.1831284 | -2.5427332 | 0.65547396 | 7.32E-06 | 5.85E-05 | Csn3 |
| ENSMUSG00( | 2.45392984 | -2.5466736 | 1.38207059 | 0.00461683 | 0.01813513 | Dlx4 |
| ENSMUSG00( | 13.7569255 | -2.5482649 | 0.5023854 | 3.23E-08 | 4.01E-07 | Klk1b11 |
| ENSMUSG00( | 5.43397672 | -2.5546611 | 0.77223226 | 5.66E-05 | 3.73E-04 | 9130016M20I |
| ENSMUSG00( | 590.24728 | -2.5622231 | 0.16427586 | 4.26E-56 | 3.08E-53 | Hbegf |
| ENSMUSG00( | 12.4436405 | -2.5648142 | 0.60385898 | 1.28E-06 | 1.19E-05 | Gm43430 |
| ENSMUSG00( | 4.25926637 | -2.5683868 | 0.83473026 | 1.21E-04 | 7.40E-04 | C630004M23 |
| ENSMUSG00( | 7.36078469 | -2.5743218 | 0.73116426 | 2.73E-05 | 1.92E-04 | Gm39460 |
| ENSMUSG00( | 188.748478 | -2.5772324 | 0.28798469 | 1.95E-20 | 1.28E-18 | Scimp |
| ENSMUSG00( | 125.417516 | -2.6210374 | 0.26820827 | 8.28E-24 | 7.55E-22 | Epdr1 |
| ENSMUSG00( | 1539.26769 | -2.6234171 | 0.23061228 | 2.79E-31 | 4.71E-29 | Rsad2 |
| ENSMUSG00( | 2.57086541 | -2.6249213 | 1.64311704 | 0.00449796 | 0.01771441 | Gm43915 |
| ENSMUSG00( | 348.792929 | -2.631896 | 0.40251978 | 3.15E-12 | 7.17E-11 | Gm49774 |
| ENSMUSG00( | 60.7668602 | -2.6324866 | 0.29622601 | 4.09E-20 | 2.57E-18 | Card6 |
| ENSMUSG00( | 39.7769034 | -2.6416512 | 0.27768942 | 1.50E-22 | 1.18E-20 | Gm12968 |
| ENSMUSG00( | 43.0791498 | -2.6444271 | 0.29365263 | 1.53E-20 | 1.02E-18 | Cish |
| ENSMUSG00( | 3.74219327 | -2.660732 | 1.53691374 | 0.001854 | 0.00824714 | Gm13836 |
| ENSMUSG00( | 6.06249034 | -2.6661734 | 1.10698929 | 0.00128876 | 0.00602657 | Timp1 |
| ENSMUSG00( | 18.3867728 | -2.6846234 | 0.75959923 | 1.66E-05 | 1.23E-04 | Mmp3 |
| ENSMUSG00( | 12.0754021 | -2.6899153 | 0.60841269 | 7.00E-07 | 6.81E-06 | Ccl17 |
| ENSMUSG00( | 7.26771426 | -2.7078741 | 0.63254256 | 1.50E-06 | 1.38E-05 | Cth |
| ENSMUSG00( | 28.6576011 | -2.7260525 | 0.32396898 | 3.20E-18 | 1.62E-16 | F3 |
| ENSMUSG00( | 168.031867 | -2.7383816 | 0.61190973 | 3.21E-07 | 3.30E-06 | Dok7 |
| ENSMUSG00( | 30.7762044 | -2.7397905 | 0.37983001 | 4.46E-14 | 1.33E-12 | Kdm5b |
| ENSMUSG00( | 3.33660411 | -2.7524661 | 1.15002974 | 0.00123754 | 0.00582472 | Mir1957a |
| ENSMUSG00( | 130.510103 | -2.7531586 | 0.20646644 | 2.02E-41 | 7.15E-39 | Smtn |
| ENSMUSG00( | 3.59417852 | -2.7718283 | 1.36972919 | 0.0030141 | 0.01260107 | Gpr17 |

| | | | | | | |
|---|---|---|---|---|---|---|
| ENSMUSG00( | 291.121122 | -2.772818 | 0.17195627 | 1.04E-59 | 7.82E-57 | Tspan7 |
| ENSMUSG00( | 45.7512785 | -2.7886529 | 0.30219573 | 1.56E-21 | 1.14E-19 | Gm2774 |
| ENSMUSG00( | 4.02127928 | -2.789308 | 1.00530951 | 3.91E-04 | 0.00208939 | |
| ENSMUSG00( | 6585.2084 | -2.7962907 | 0.21286394 | 1.09E-40 | 3.71E-38 | Mmp12 |
| ENSMUSG00( | 5.04313252 | -2.8050446 | 1.15821665 | 6.23E-04 | 0.00317282 | |
| ENSMUSG00( | 80.0651833 | -2.8125621 | 0.31713469 | 3.25E-20 | 2.06E-18 | Rdm1 |
| ENSMUSG00( | 12.2151223 | -2.8152269 | 0.73606253 | 5.93E-06 | 4.85E-05 | Csta2 |
| ENSMUSG00( | 13.7668604 | -2.8221011 | 0.52483552 | 4.24E-09 | 6.08E-08 | Pgf |
| ENSMUSG00( | 29.9839242 | -2.8333524 | 0.3856564 | 1.99E-14 | 6.34E-13 | Cd200r1 |
| ENSMUSG00( | 27.2763567 | -2.8403286 | 0.46621638 | 5.98E-11 | 1.14E-09 | Gm35996 |
| ENSMUSG00( | 4.98361439 | -2.8404102 | 1.04047934 | 6.06E-04 | 0.00309673 | Ptgds |
| ENSMUSG00( | 13.4700385 | -2.8472648 | 0.49602255 | 5.00E-10 | 8.24E-09 | Muc3 |
| ENSMUSG00( | 2.94930818 | -2.8508538 | 1.66745192 | 0.00353988 | 0.01445143 | Tekt3 |
| ENSMUSG00( | 92.6854887 | -2.8540316 | 0.19938057 | 1.80E-47 | 7.93E-45 | Zfp827 |
| ENSMUSG00( | 75.1523573 | -2.8548789 | 0.27993984 | 1.49E-25 | 1.58E-23 | C1qtnf6 |
| ENSMUSG00( | 6.14695452 | -2.8707643 | 0.79996864 | 2.50E-05 | 1.78E-04 | Gm18422 |
| ENSMUSG00( | 182.810933 | -2.8988255 | 0.3274301 | 4.68E-20 | 2.94E-18 | Upp1 |
| ENSMUSG00( | 10.5317482 | -2.9378971 | 0.65697603 | 4.90E-07 | 4.89E-06 | Serpind1 |
| ENSMUSG00( | 12.9623078 | -2.9494431 | 0.52674384 | 2.09E-09 | 3.18E-08 | Cpn1 |
| ENSMUSG00( | 49.1125753 | -2.9513668 | 0.38001332 | 7.61E-16 | 2.90E-14 | Cx3cl1 |
| ENSMUSG00( | 18.2609247 | -2.9658543 | 0.64820106 | 2.05E-07 | 2.18E-06 | |
| ENSMUSG00( | 12.4862183 | -2.9709778 | 0.5424427 | 3.22E-09 | 4.74E-08 | Tnfsfm13 |
| ENSMUSG00( | 1019.73198 | -2.9749367 | 0.13280289 | 8.30E-113 | 2.50E-109 | S100a6 |
| ENSMUSG00( | 4.68069811 | -2.9935788 | 1.20130182 | 0.00157066 | 0.0071577 | Tmem98 |
| ENSMUSG00( | 12.3094377 | -2.9967513 | 0.54130232 | 2.67E-09 | 3.99E-08 | Prkg1 |
| ENSMUSG00( | 5.30467581 | -2.9988428 | 0.89389796 | 6.23E-05 | 4.07E-04 | Gm28874 |
| ENSMUSG00( | 1.80973541 | -3.0157064 | 1.79437988 | 0.00316229 | 0.01313549 | Gm13940 |
| ENSMUSG00( | 11.6224064 | -3.0462725 | 0.66209644 | 4.85E-07 | 4.84E-06 | Erbb3 |
| ENSMUSG00( | 2.39944645 | -3.0561862 | 1.27748489 | 0.00167405 | 0.00755077 | Abcb9 |
| ENSMUSG00( | 56.0155758 | -3.0663297 | 0.3198817 | 1.21E-22 | 9.63E-21 | Entrep1 |
| ENSMUSG00( | 708.132257 | -3.085499 | 0.15099056 | 6.16E-94 | 1.01E-90 | Cd93 |
| ENSMUSG00( | 31.1856859 | -3.0951953 | 0.46044424 | 1.19E-12 | 2.87E-11 | Disp3 |
| ENSMUSG00( | 10.3459427 | -3.1228129 | 0.62150646 | 3.57E-08 | 4.40E-07 | Rgs16 |
| ENSMUSG00( | 93.456745 | -3.1425427 | 0.25055016 | 3.44E-37 | 9.13E-35 | Ehd2 |
| ENSMUSG00( | 10.1315627 | -3.1482119 | 0.92613446 | 1.19E-04 | 7.33E-04 | Zcchc24 |
| ENSMUSG00( | 7.8325117 | -3.1498302 | 0.87783818 | 2.80E-05 | 1.96E-04 | Gbp4 |
| ENSMUSG00( | 27.41959 | -3.155296 | 0.38898696 | 4.57E-17 | 2.02E-15 | Spry4 |
| ENSMUSG00( | 37.865681 | -3.1605164 | 0.49277535 | 1.83E-11 | 3.74E-10 | F11r |
| ENSMUSG00( | 9.73663889 | -3.1617877 | 0.55878182 | 1.02E-09 | 1.62E-08 | Rnu5g |
| ENSMUSG00( | 28.0730712 | -3.181488 | 0.39063809 | 2.50E-17 | 1.13E-15 | Serpina3f |
| ENSMUSG00( | 2.75697447 | -3.1876005 | 1.52422787 | 0.00167892 | 0.00756709 | Golga7b |

| | | | | | | |
|---|---|---|---|---|---|---|
| ENSMUSG00( | 8.5209313 | -3.211169 | 1.21054964 | 0.00103096 | 0.00497155 | Gm13091 |
| ENSMUSG00( | 3.0847238 | -3.2464631 | 1.3967127 | 9.46E-04 | 0.00462076 | Csta3 |
| ENSMUSG00( | 15.9855779 | -3.2776932 | 0.56245326 | 9.01E-10 | 1.44E-08 | Hap1 |
| ENSMUSG00( | 31.3746547 | -3.2839981 | 0.48903025 | 2.38E-12 | 5.50E-11 | Mylk |
| ENSMUSG00( | 16.4376089 | -3.2869287 | 0.44116073 | 6.15E-15 | 2.10E-13 | Cyp1a2 |
| ENSMUSG00( | 5.23416426 | -3.2884647 | 1.0858157 | 2.43E-04 | 0.00137249 | Scml4 |
| ENSMUSG00( | 50.44684 | -3.3349631 | 0.3446206 | 3.19E-23 | 2.76E-21 | Gm9844 |
| ENSMUSG00( | 7.21863189 | -3.3366835 | 0.91062095 | 1.67E-05 | 1.23E-04 | Rtn4r |
| ENSMUSG00( | 281.856925 | -3.3504582 | 0.15482385 | 2.95E-105 | 6.65E-102 | Ndrg1 |
| ENSMUSG00( | 10.8366729 | -3.3586489 | 0.64814229 | 2.64E-08 | 3.31E-07 | Nox1 |
| ENSMUSG00( | 3.2798158 | -3.3953128 | 1.52287392 | 9.64E-04 | 0.00469333 | Gm23444 |
| ENSMUSG00( | 51.6197138 | -3.4024086 | 0.26718026 | 4.74E-38 | 1.36E-35 | Edn1 |
| ENSMUSG00( | 11.1179305 | -3.4614364 | 0.66387035 | 1.85E-08 | 2.38E-07 | Ifi206 |
| ENSMUSG00( | 36.1786741 | -3.4771008 | 0.29949053 | 2.58E-32 | 4.67E-30 | Prkg2 |
| ENSMUSG00( | 13.653457 | -3.4793882 | 0.6262196 | 1.44E-09 | 2.26E-08 | Gm23849 |
| ENSMUSG00( | 4.62281295 | -3.5028549 | 1.24470111 | 5.07E-04 | 0.00264003 | Gm45879 |
| ENSMUSG00( | 20.9443674 | -3.5057071 | 0.48538611 | 4.00E-14 | 1.21E-12 | Csf2 |
| ENSMUSG00( | 2.44160069 | -3.5644204 | 1.67283295 | 0.00133247 | 0.00620208 | Rpl10a-ps3 |
| ENSMUSG00( | 8.93035502 | -3.5662042 | 0.72391099 | 1.21E-07 | 1.34E-06 | 1700006J14R |
| ENSMUSG00( | 3.82095865 | -3.5878203 | 1.28411769 | 2.28E-04 | 0.0013003 | Gm25099 |
| ENSMUSG00( | 2.50898309 | -3.5963119 | 1.70422987 | 0.00135162 | 0.00627987 | A930023M06I |
| ENSMUSG00( | 64.7140049 | -3.6133036 | 0.31473315 | 1.11E-31 | 1.97E-29 | Gm6093 |
| ENSMUSG00( | 50.3992669 | -3.6566385 | 0.45353662 | 1.73E-16 | 7.05E-15 | F5 |
| ENSMUSG00( | 995.074294 | -3.6786306 | 0.15951736 | 8.29E-119 | 3.75E-115 | Tmsb10 |
| ENSMUSG00( | 7.12607925 | -3.6920174 | 0.871095 | 1.44E-06 | 1.32E-05 | Rnu1a1 |
| ENSMUSG00( | 3.73590881 | -3.7269243 | 1.22050127 | 3.19E-04 | 0.00174479 | Gm6637 |
| ENSMUSG00( | 26.7818854 | -3.7443946 | 0.67753473 | 6.69E-09 | 9.26E-08 | Nsmaf |
| ENSMUSG00( | 4.33610158 | -3.8837141 | 1.05898499 | 2.66E-05 | 1.88E-04 | Rpl31-ps17 |
| ENSMUSG00( | 62.6507769 | -3.9367639 | 0.30787864 | 1.02E-38 | 3.08E-36 | Gpr68 |
| ENSMUSG00( | 232.856393 | -4.0711669 | 0.19380685 | 7.40E-99 | 1.34E-95 | Fabp3 |
| ENSMUSG00( | 73.222274 | -4.1715453 | 0.28792821 | 1.85E-48 | 8.34E-46 | Ass1 |
| ENSMUSG00( | 7.57963686 | -4.2295456 | 0.89485449 | 3.46E-07 | 3.55E-06 | Enthd1 |
| ENSMUSG00( | 25.8032859 | -4.238943 | 0.59618652 | 4.26E-13 | 1.10E-11 | Lipg |
| ENSMUSG00( | 3.18604494 | -4.2585479 | 2.41232297 | 0.00118684 | 0.00561238 | Gm24830 |
| ENSMUSG00( | 424.637126 | -4.3319794 | 0.18853927 | 3.29E-117 | 1.19E-113 | Axl |
| ENSMUSG00( | 3.49950583 | -4.3459741 | 1.44683699 | 2.18E-04 | 0.00124496 | Gm22973 |
| ENSMUSG00( | 8.32424645 | -4.4014458 | 1.0329395 | 2.91E-06 | 2.52E-05 | Gm15978 |
| ENSMUSG00( | 28.6627552 | -4.409933 | 0.44054485 | 1.25E-24 | 1.23E-22 | B930036N10I |
| ENSMUSG00( | 10.8293119 | -4.4223447 | 0.64349975 | 9.33E-13 | 2.28E-11 | Gm22513 |
| ENSMUSG00( | 9.92069839 | -4.591183 | 0.84291679 | 3.20E-09 | 4.73E-08 | Gm23971 |
| ENSMUSG00( | 7.67095727 | -4.593809 | 0.95981363 | 1.21E-07 | 1.34E-06 | Snord118 |

| | | | | | | |
|---|---|---|---|---|---|---|
| ENSMUSG00( | 15.9807453 | -4.6009265 | 0.68970218 | 6.11E-12 | 1.33E-10 | Gm3788 |
| ENSMUSG00( | 90.0128367 | -4.6696542 | 0.31067608 | 5.88E-52 | 3.22E-49 | Ahr |
| ENSMUSG00( | 3.73831596 | -4.8450105 | 2.64987213 | 4.34E-04 | 0.00229913 | Rgs8 |
| ENSMUSG00( | 10.8932933 | -5.2440629 | 0.83923762 | 4.01E-11 | 7.85E-10 | Rnu2-10 |
| ENSMUSG00( | 3.38890975 | -5.2890497 | 1.53421729 | 2.33E-05 | 1.67E-04 | Gm22265 |
| ENSMUSG00( | 21.0773837 | -5.3117453 | 0.62370919 | 5.55E-18 | 2.70E-16 | Nyap2 |
| ENSMUSG00( | 15.1938851 | -5.3840321 | 0.6909877 | 2.30E-15 | 8.23E-14 | Cstdc4 |
| ENSMUSG00( | 7.59233705 | -5.4185328 | 0.96631769 | 5.82E-09 | 8.17E-08 | Rnu12 |
| ENSMUSG00( | 15.7019005 | -5.4561646 | 2.40833392 | 1.71E-05 | 1.26E-04 | Ighm |
| ENSMUSG00( | 15.3804954 | -5.7232346 | 1.01289946 | 1.95E-08 | 2.49E-07 | Trim63 |
| ENSMUSG00( | 144.76251 | -5.7976773 | 0.26158135 | 1.02E-109 | 2.63E-106 | Grcc10 |
| ENSMUSG00( | 13.4146752 | -5.9029591 | 2.44106657 | 3.77E-06 | 3.22E-05 | Ablim1 |
| ENSMUSG00( | 2.36954567 | -6.1030968 | 2.85311474 | 1.00E-04 | 6.25E-04 | Col1a1 |
| ENSMUSG00( | 5.86367263 | -6.2054625 | 1.44766716 | 5.32E-07 | 5.27E-06 | 2900060B14F |
| ENSMUSG00( | 6.00297631 | -6.2543385 | 1.3834216 | 1.79E-07 | 1.93E-06 | Gm26316 |
| ENSMUSG00( | 875.089232 | -6.8821877 | 0.27412491 | 1.08E-138 | 1.95E-134 | Hspb7 |
| ENSMUSG00( | 49.2832222 | -8.0407456 | 0.73518134 | 4.34E-27 | 5.19E-25 | |
| ENSMUSG00( | 111.959434 | -8.7485793 | 1.29541599 | 3.54E-14 | 1.08E-12 | Mcoln3 |
| ENSMUSG00( | 20.4914399 | -10.799749 | 3.02559307 | 1.40E-15 | 5.15E-14 | Gm24407 |

e_name
Rik

Rik

Rik

Rik

ῤik

ℜik

Rik

Rik

Rik

Rik

Rik

Rik

ik

Rik

Rik

ik

Rik

ik

Rik

Rik

Rik

Rik

Rik

Rik

Rik

Rik

Rik
ik

Rik

ik
Rik

Rik

ik

Rik

Rik

Rik

ʀik

ʀik

Rik

Rik

Rik

Rik

Rik

Rik

Rik

Rik

Rik

Rik

Rik

Rik

Rik

Rik

Rik

Rik

Rik

Rik

Rik

ik

Rik

Rik

Rik
