## Supplemental information, figure compressed, table 1-5 for "Role of copper during microglial inflammation": Supplemental Table 4-compressed.pdf

| | |
|---|---|
| *Hprt* _F | TCAGTCAACGGGGGACATAAA |
| *Hprt* _R | GGGGCTGTACTGCTTAACCAG |
| *Il-1b* _F | GCAACTGTTCCTGAACTCAACT |
| *Il-1b* _R | ATCTTTTGGGGTCCGTCAACT |
| *Il-6* _F | CCACTTCACAAGTCGGAGGCTT |
| *Il-6* _R | CCAGCTTATCTGTTAGGAGA |
| *Nos2* _F | GTTCTCAGCCCAACAATACAAGA |
| *Nos2* _R | GTGGACGGGTCGATGTCAC |
| *Il-1a* _F | CGAAGACTACAGTTCTGCCATT |
| *Il-1a* _R | GACGTTTCAGAGGTTCTCAGAG |
| *Tnfa* _F | CCCTCACACTCAGATCATCTTCT |
| *Tnfa* _R | GCTACGACGTGGGCTACAG |
| *Slc31a1* _F | TATGAACCACACGGACGACAA |
| *Slc31a1* _R | GCCATTTCTCCAGGTGTATTGA |
| *Atp7a* _F | TGGGAAAGTGAATGGTGTCCA |
| *Atp7a* _R | ACGGTATTGGTTAAGACAGGGA |
