## Supplemental information, figure compressed, table 1-5 for "Role of copper during microglial inflammation": Supplemental Table 5-compressed.pdf

shRNA Sample
Atp7a. KD1
Atp7a KD2
Slc31a1 KD1
Slc31a1 KD2
CTRL-Luci.1309

shRNA Oligo Sequence

atTGCTGTTGACAGTGAGCGCAAGGTGTTTGCTGAAGTTCTATAGTGAAGCCACAGATGTATAGAACTTCAGCAAACACCTTAT

atTGCTGTTGACAGTGAGCGAAACAGCGTTGTTACTAGTGAATAGTGAAGCCACAGATGTATTCACTAGTAACAACGCTGTTCT

atTGCTGTTGACAGTGAGCGCAAGTTCCTAGGTGATGTTCTATAGTGAAGCCACAGATGTATAGAACATCACCTAGGAACTTAT

atTGCTGTTGACAGTGAGCGCCCCAGGTAGCTTAGTGTTTAATAGTGAAGCCACAGATGTATTAAACACTAAGCTACCTGGGAT

atTGCTGTTGACAGTGAGCGCCCGCCTGAAGTCTCTGATTAATAGTGAAGCCACAGATGTATTAATCAGAGACTTCAGGCGGT[illegible]
