## Supplementary figures and images for "Role of copper during microglial inflammation"

### SupplementalFigures_Copper-compressed.pdf

**Supplemental Figure 1**

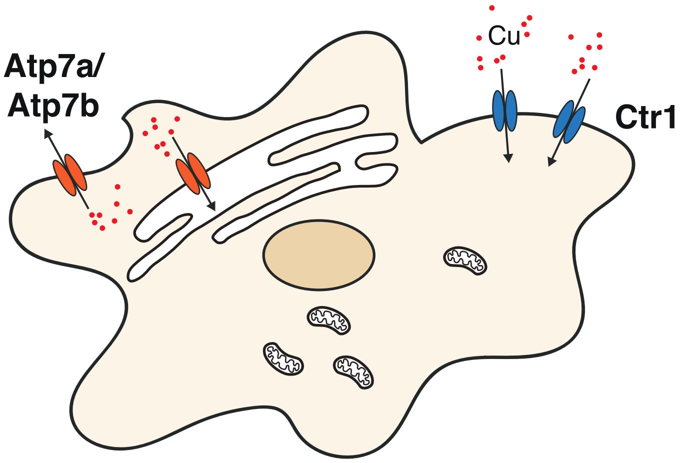

**Supplemental Figure 2**

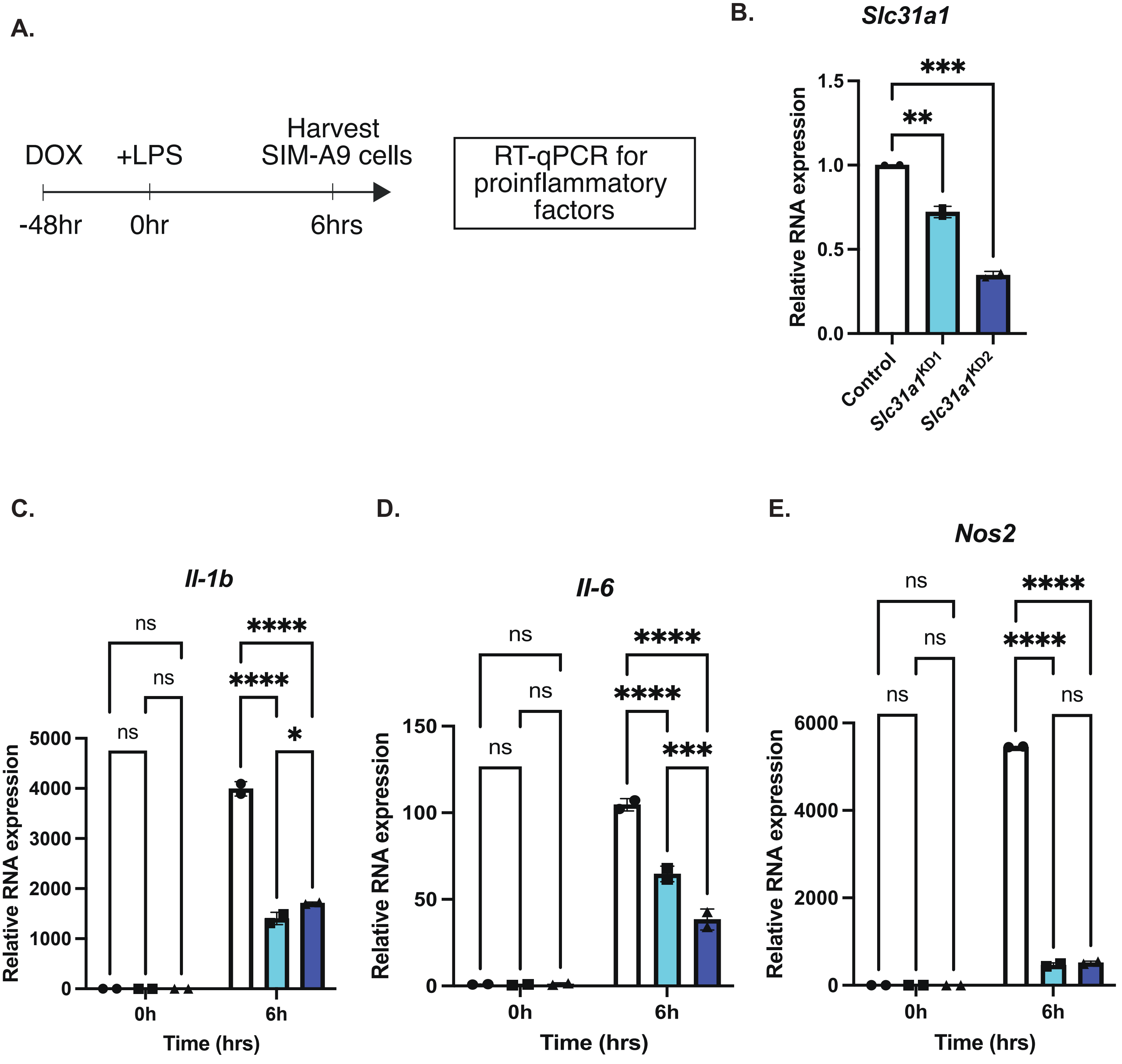

Supplemental Figure 3

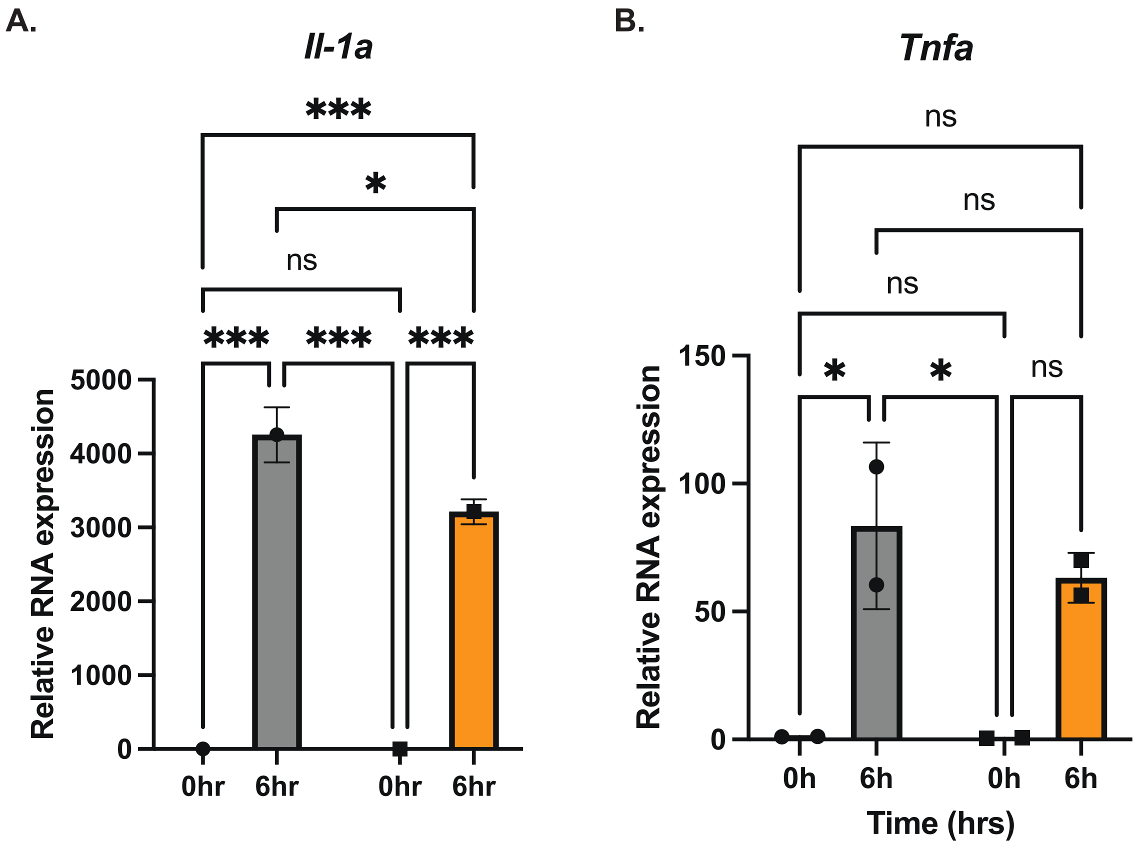

**Supplemental Figure 4**

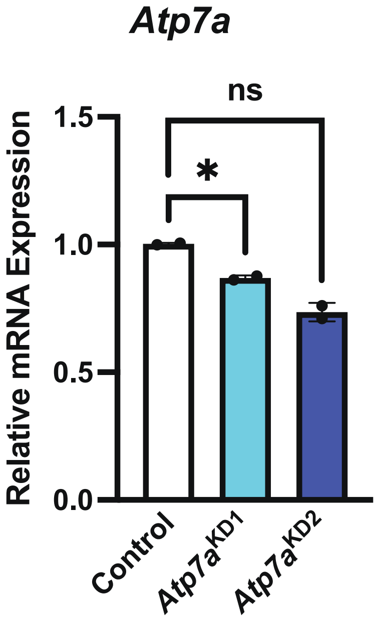
